# Modeling epithelial tissue and cell deformation dynamics using a viscoelastic slab sculpted by surface forces

**DOI:** 10.1101/2022.12.23.521823

**Authors:** XinXin Du, Michael Shelley

**Affiliations:** Center for Computational Biology, Flatiron Institute, New York, New York 10010, USA; Courant Institute of Mathematical Sciences, New York University, New York, New York 10012, USA

## Abstract

During morphogenesis, epithelial monolayers actively alter their shape to create future body parts of the animal; this makes the epithelium one of the most active and critical structures in early animal development. Even though epithelial cells exist and move in three dimensions, mathematical models frequently describe them as merely two-dimensional. However, recent imaging technology has begun to reveal pertinent dynamics in the third dimension of the tissue. With the importance of the third dimension in mind, we have developed a self-sculpting, three-dimensional model of epithelia whose dynamics are driven by active forces on its surface. We present a first, fundamental study for a reduced version of epithelia that investigates how surface forces affect its internal dynamics. Our model captures the 3D slab-like geometry of epithelia, viscoelasticity of tissue response, fluid surroundings, and driving from active surface forces. We represent epithelial tissue as a thick slab, a 3D continuum comprised of a Stokes fluid with an extra viscoelastic stress. Employing this model, we present both analytical and numerical solutions of the system and make quantitative predictions about cell shapes, cell dynamics, and the tissue’s response to surface force in a three-dimensional setting. In particular, we elucidate the initiation of ventral furrow invagination and T1 transitions in *Drosophila* embryogenesis. In the former, we demonstrate the importance of fluid and geometric surroundings to drive invagination. In the latter, we show the limitations of surface forces alone to drive T1 transitions.

## II. INTRODUCTION

Epithelial cells change shape and rearrange while staying connected in a planar geometry. While many events in morphogenesis, such as tube formation [1] or epithelial invagination [2, 3], involve deforming the planar geometry of the epithelium, many of these deformations are preceded by morphological changes in which cells move merely in-plane to the epithelium [4]. Moreover, many events in morphogenesis do not deform the epithelial plane at all, as they consist only of in-plane movement; examples of this include convergent extension in both *Drosophila* [5–7] and vertebrates [8] and ommatidia rotation in *Drosophila* [9].

Whenever epithelial cells move mostly in-plane to the epithelium, authors often use 2D vertex models [10–14] or particle models [15, 16] to describe them. These models are sufficient if the epithelial cells they describe are thin. However, recent developments in microscopy have revealed that when cells have significant height, movements become complex [5], and a three-dimensional approach may be required. To this end, 3D vertex models [17–20] and 3D finite element models [21–23] have been used. While these models have produced effective results to describe both cell rearrangements and tissue deformations, they do not take into account the fluid surroundings of the tissue, nor do they track internal velocity fields and stresses. Moreover, these approaches require many phenomenological parameters to be specified and tuned. The thin viscoelastic shell models of [24, 25] treat the mechanics and activities of the actin cortex near cell surfaces as continuum fields and are able to describe in detail the 3D shapes of individual cells, however the influence of fluid surroundings on the active layer are not modeled. Our approach restricts to modeling phenomena in which cells move mostly in-plane to the epithelium; however, our model epithelium is fully three dimensional, and we consider its interaction with both fluid/material surroundings and boundary constraints.

We present a mathematical model for epithelial tissue in which the tissue is described by a three-dimensional continuum viscoelastic fluid, geometrically, a flat 3D slab, surrounded by viscous fluids. The epithelium is driven by active surface forces, for example due to populations of actomyosin that exist on the apical or basal surface of cells but not on their lateral sides [26]. One question that our model treats explicitly is how the thickness of the epithelium plays a role in how forces alter cell shape: since forces of contractility are specified only at the surface of the tissue, non-zero thickness means that forces specified on the apical (basal) side do not fully propagate to the basal (apical) side. This would imply that the topology of cells can look different on the apical versus basal surface due to the attenuation of forces through the thickness of the tissue. Our mathematical description quantitatively comments on the nature of force transfer from one side of the epithelium to the other.

Finally, because we model the tissue as a continuum, velocity fields are used to track material elements. If we assume that cell membranes do not exert significant force, that is, that membranes are merely carried by the flow of cytoplasm [26], then cell boundaries, tracked as material elements, provide simulated data of 3D cell shape change over time. Figure 1 shows the results of thus simulated 3D cell deformations under different driving surface forces (elaborated in Fig. 7). In this paper, we simulate both the onset of ventral furrow invagination and convergent-extension in *Drosophila*. We elucidate these events using 3D solutions to the system, accounting for the fluid environment and other geometric constraints.

**FIG. 1.**
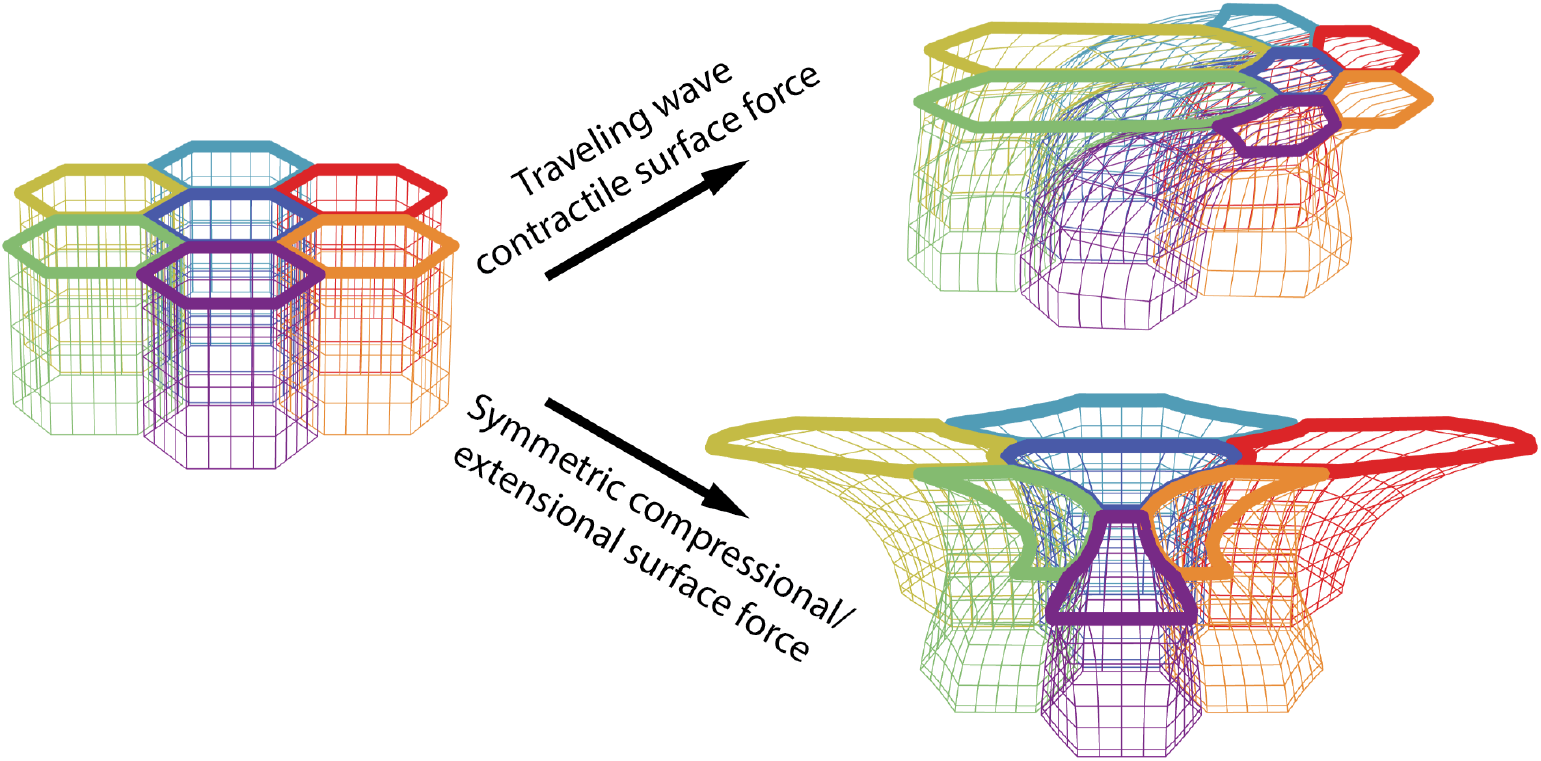
Simulated 3D cell deformations under two different applied surface forces.

## III. DEFINITION AND SOLUTION TO THE MODEL

### A. Biological motivation for model geometry

Biologically and stereotypically, epithelia are flat tissues that, on one side, are separated from a solid-like structure by a layer of fluid, and on the other side, adjacent to a fluidic bath. In the fly embryo, for example, the apical side of the epithelium is located next the perivitelline fluid layer which borders the vitelline membrane, while the basal side faces the fluid at the center of the embryo (Fig. 2(a)) [2, 3].

**FIG. 2.**
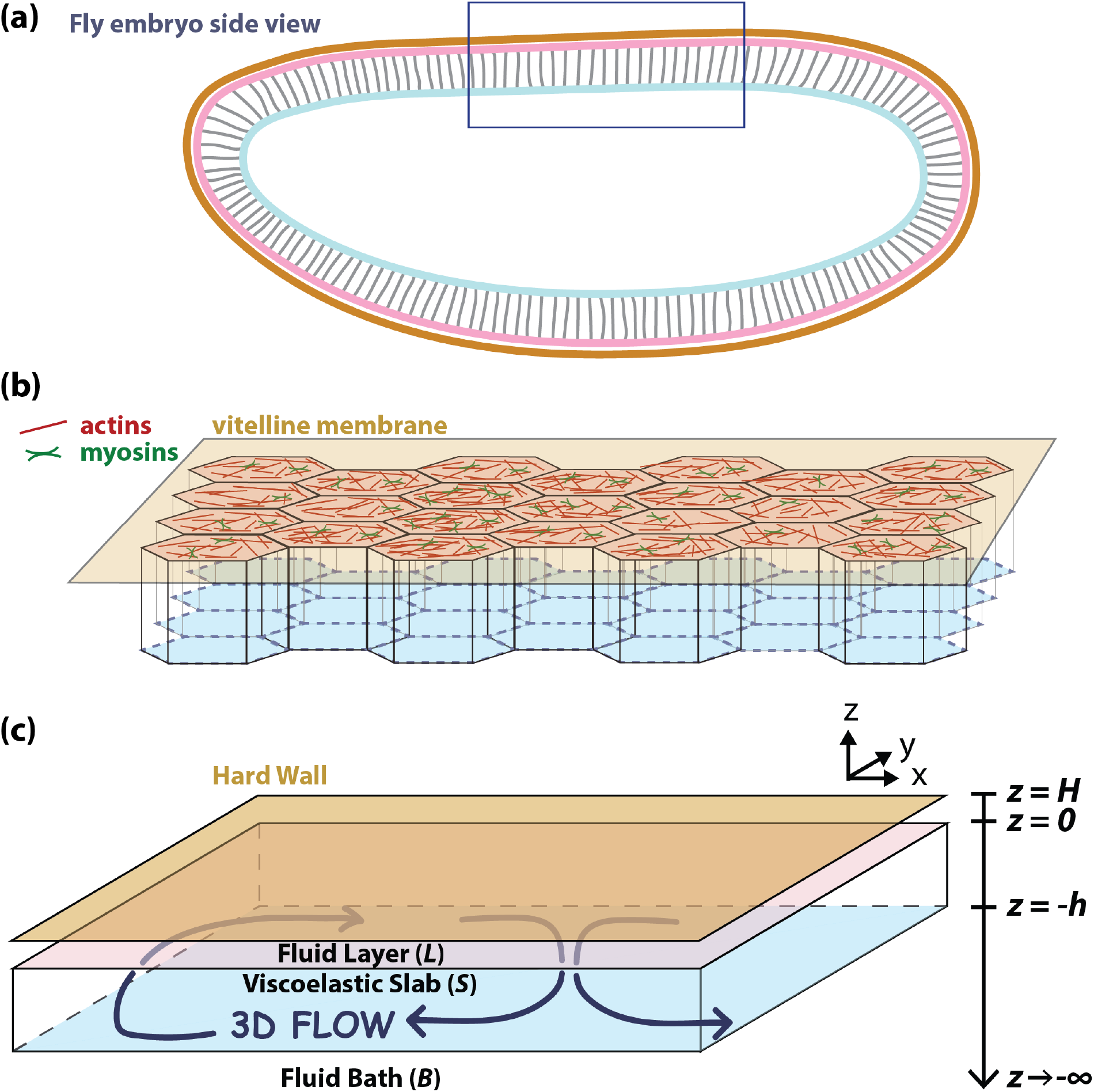
System geometry. (a) Stereotypical epithelial geometry of the *Drosophila* embryo: one side of the epithelium is adjacent to a fluid layer (perivitelline fluid) and a hard wall (vitelline membrane); the other side faces a fluid bath. The box indicates a region of approximately flat epithelium. (b). Schematic of epithelium with an active surface containing actomyosin adjacent to the perivitelline fluid and vitelline membrane. (c) Diagram of model epithelium with definitions of regions *L, S, B*, and the coordinate system; the two surfaces of the slab are “top” (*z* = 0) and “bottom” (*z* = −*h*).

To describe the epithelium mathematically, as a mechanical object, we use a viscoelastic fluid model to represent the epithelium [27–29]. Because an epithelium is longer in two directions and shorter in the third, we describe it as a 3D slab that is periodic in the *x* and *y* directions with finite thickness *h* in the *z* direction. Above the slab we include a layer of Newtonian fluid (region *L*), simulating the layer between the epithelium and a solid-like structure, typically a vitelline membrane or extracellular matrix. The solid-like structure itself is represented by a hard wall at *z* = *H*. The viscoelastic slab occupies the domain −*h* ≤ *z* ≤ 0 (region *S*). Below the slab, for *z <* −*h*, we include a fluid bath of infinite depth (region *B*) to simulate the admittedly finite but large-scale fluidic bath that often exists on the other side of the epithelium. We refer to the side of the slab adjacent to the fluid layer at *z* = 0 as the “top”, and the side adjacent to the semi-infinite fluid bath at *z* = −*h* as the “bottom”. Figure 2(c) shows regions *L, S*, and *B* in a schematic diagram.

Active forces are applied along the surface of the slab in the planar directions (*x* and *y*). These active forces typically arise from populations of myosin motors moving on an actin mesh that exists on the surface of the tissue (Fig. 2(b)). While the forces themselves are two-dimensional, they create a three-dimensional flow inside the tissue because of the internal mechanics of the slab (Fig. 2(c)). In this paper, we analyze these 3D flows given the particular form of the surface force. Assuming that cell boundaries are carried by the flow, we also characterize cell deformations that arise from this force.

### B. The Model

#### 1. Bulk equations

Let the fields ***σ***^*L*^, ***σ***^*S*^, and ***σ***^*B*^ denote the total stress tensors in the *L, S*, and *B* regions of the system:

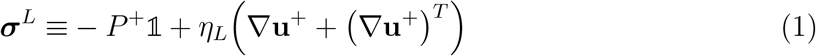

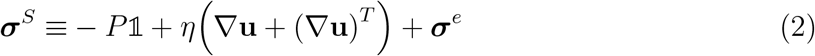

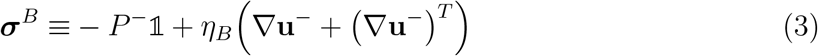

where **u, u**^±^ are the velocities, *P, P*^±^ are the pressures, and *η, η*_*L,B*_ are the viscosities in each region. Here, ***σ***^*L,B*^ indicate linear Newtonian solvents and ***σ***^*e*^ indicates an extra stress. We model the tissue as an Oldroyd-B fluid because Oldroyd-B is perhaps the simplest model of a viscoelastic fluid. Requiring:

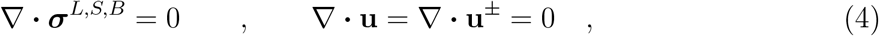

we obtain the Stokes or forced Stokes equations of motion for fields **u, u**^±^, *P, P*^±^. The extra stress ***σ***^*e*^ in Eq. (2) is evolved by the equation:

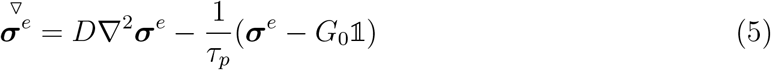

where

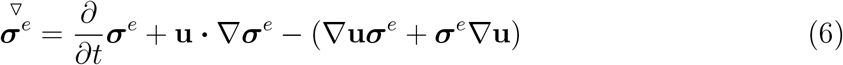

is the upper-convected time derivative. Further, the origins of Oldroyd-B in a microscopic description gives that there is no diffusive flux at *z* = 0 and *z* = −*h*, that is:

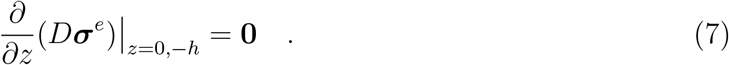

To non-dimensionalize Eqns. (1-5), we rescale time, length, and force as:

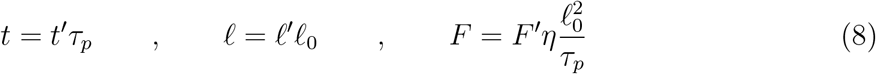

where ℓ_0_ is a yet-unspecified characteristic length scale. We additionally rescale ***σ***^*e*^ by a dimensionless stress *α*: ***σ***^*e*^ = ***σ***^*e*^′*αη/τ*_*p*_ where *αη/τ*_*p*_ = *G*_0_. We chose dimensional units to depend on the material properties of the system (the solvent viscosity *η* and polymer relaxation time *τ*_*p*_), so that parameters specifying the driving force, e.g. its amplitude and frequency, will be free to vary. The details of non-dimensionalization are in section (SI) of the Supplement.

Using the scalings in Eq. (8), the non-dimensional equations of motion in the fluid layer (region *L*), the slab (region *S*), and the fluid bath (region *B*) are:

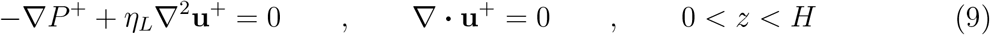

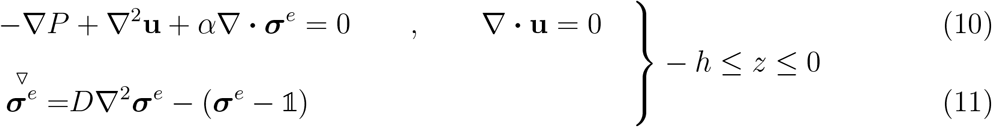

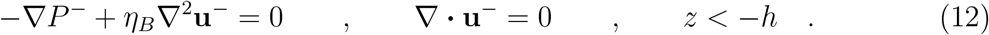

#### 2. Boundary conditions

We assume a no-slip condition on the wall:

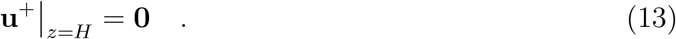

Additionally, we assume that at *z* = 0 and *z* = −*h*, we have continuity of velocity with a zero-velocity condition in the *z* direction:

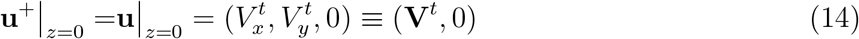

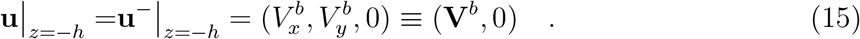

Here, the planar (*x, y*) velocity at *z* = 0 is denoted 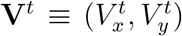 where the superscript denotes the “top” surface of the slab, and the planar velocity at *z* = −*h* is denoted 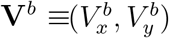 where the superscript denotes the “bottom” surface of the slab. We also demand that velocities at −∞ vanish:

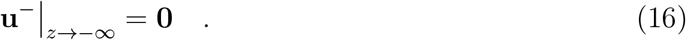

Finally, we have boundary conditions reflecting the fact that actively applied tangential forces at the top (*z* = 0) and bottom (*z* = −*h*) surfaces of the slab create stress jumps across those surfaces. Let **F**^*t*^ and **F**^*b*^ notate stresses from active driving forces applied at the top and bottom surfaces, respectively. Then:

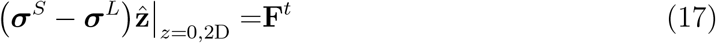

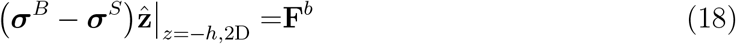

where the notation 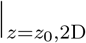 is defined by 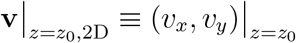 for an arbitrary 3D vector **v** = (*v*_*x*_, *v*_*y*_, *v*_*z*_). The *z* directional stress at the top and bottom surfaces are determined by the condition that *w* = 0 at *z* = 0, −*h*.

### C. Fourier representation of solution when *σ*^*e*^ = 0: the Stokes solution

If we assume that the slab is a simple Stokes fluid, setting ***σ***^*e*^ = **0** in Eq. (10) and eliminating Eq. (11) entirely, then the velocity field can be determined everywhere in the system as a nonlocal functional of the surface forces [30]. This involves solving a Neumann-to-Dirichlet mapping from surface forces to surface velocities. We Fourier transform the Stokes equations in *x* and *y*. Stress boundary conditions are applied to 3D velocity solutions such that the surface forces **F**^*t*^ and **F**^*b*^ self-consistently determine the surface velocities **V**^*t*^ and **V**^*b*^ through a Neumann-to-Dirichlet map.

A 2D Fourier Transform of an arbitrary scalar function *f* is defined as:

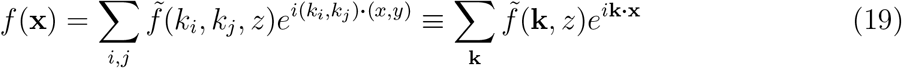

where **k** takes on values:

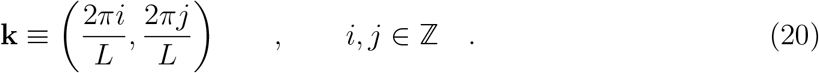

We notate 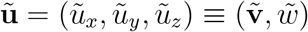 so that 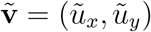. Using Eq. (19) to Fourier transform all fields in the Stokes equation 0 = −∇*P* + *η*∇^2^**u** and applying the divergence-less condition ∇ · **u** = 0 leads to the 2D transformed Stokes equations and divergence-less condition:

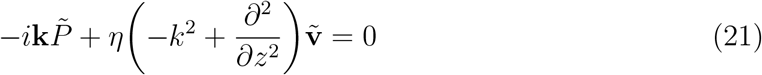

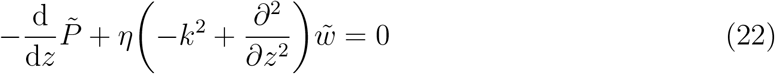

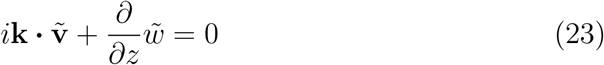

where all fields are functions of (**k**, *z*) and *k* ≡ |**k**|. The boundary conditions for 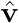 and 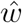, from Eqns. (14) and (15), are:

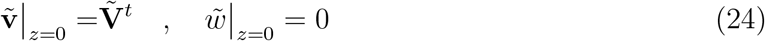

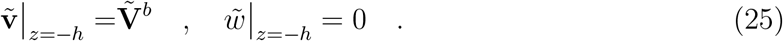

Solutions for Eqns. (21)-(23) are presented in section (SII A) of the Supplement.

### D. Neumann-to-Dirichlet map of surface forces to surface velocities

Using the expressions for 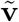 and 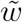 in Eqs. (S30) and (S26), the tangential shear stresses 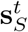 and 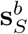 evaluated at the boundaries *z* = 0 and *z* = −*h* from region *S* are:

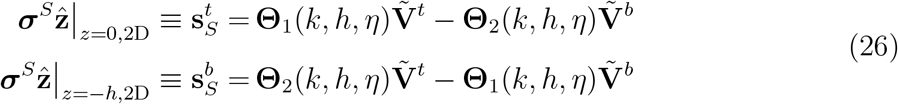

where the matrix operators **Θ**_1_ and **Θ**_2_ (given in Eq. (S49)) acting on 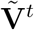 and 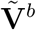 depend on the magnitude of the wavevector *k* and parameters *h* and *η* (rescaled to 1 from nondimensionalization but reinstated here for clarity). A similar calculation in the Supplement gives, for tangential shear stresses from regions *L* and *B*:

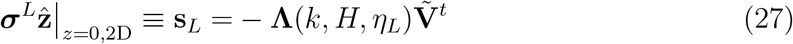

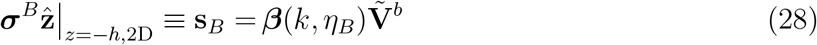

where the matrix operators **Λ** and ***β*** are given in Eq. (S51) and (S52). Assuming that the stress jumps at *z* = 0 and *z* = −*h* are created by active forces **F**^*t*^ and **F**^*b*^, respectively, we obtain:

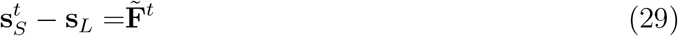

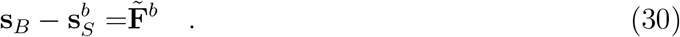

Inverting these equations, we express 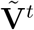 and 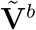 in terms of 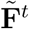 and 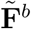:

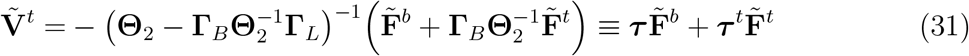

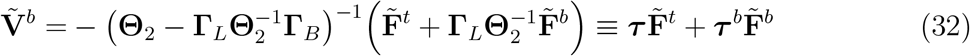

where **Γ**_*B*_ ≡ **Θ**_1_ + ***β*** and **Γ**_*L*_ ≡ **Θ**_1_ + **Λ**. Note that the matrices **Θ**_1,2_, **Γ**_*L,B*_, ***β*, Λ**, all have the form 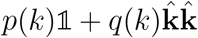 where *p* ≠ 0 and *q* ≠ −*p*. Hence each is invertible with its inverse of the form 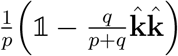 where 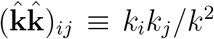; additionally, they all commute under multiplication. Hence, the same matrix ***τ*** multiplies 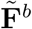 in Eq. (31) and 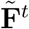 in Eq. (32), meaning that the transfer of surface forces to velocities on opposite surfaces are identical regardless of direction of transfer. Further, the transfer matrices ***τ, τ***^*t,b*^, like the other matrices in this discussion, are also nonsingular rank-one perturbations of the identity. Thus, we have determined **V**^*t,b*^ as functionals of surface forces **F**^*t,b*^. Finally, Eqns. (S26), (S28), (S30), (S44)-(S46), and (S39)-(S41) give the bulk velocities **u, u**^±^ and pressures *P, P*^±^ in terms of **V**^*t,b*^.

### E. Stokes Oldroyd-B solution

To construct the full solution to the Stokes Oldroyd-B equation (Eq. (10)) in region *S*, we need to consider the source term *α*∇ · ***σ***^*e*^. We first construct, numerically, a particular solution to Eq. (10) using zero velocity boundary conditions at *z* = −*h*, 0. We add to this a homogenous solution (as in the previous section) that corrects the boundary conditions and so obtain the full solution. This is detailed section (SIII) of the Supplement. The end result is that the quantities:

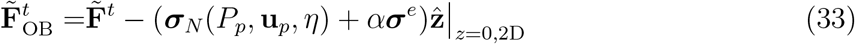

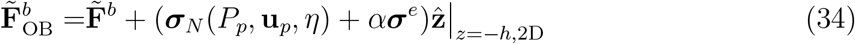

are substituted for 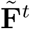 and 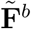 to find velocities 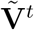 and 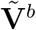 in Eqns. (31) and (32). In the above, **u**_*p*_ and *P*_*p*_ indicate the numerical particular solution and ***σ***_*N*_ (*P*, **u**, *η*) ≡ −*P* 1 + *η*(∇**u** + ∇**u**^*T*^). Explicitly, we have:

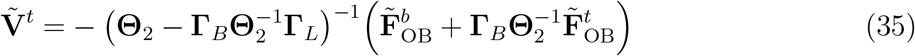

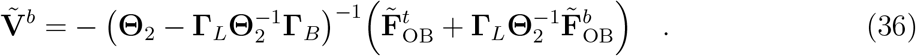

Quantities 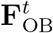 and 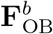 now include active forces **F**^*t*^ and **F**^*b*^ as well induced forces from the extra stress and the particular solution in the slab. As before, these 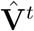 and 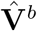 are utilized in Eqns. (S26), (S28), (S30), (S44)-(S46), and (S39)-(S41) to obtain **u**_*h*_, **u**^±^, *P*_*h*_, *P*^±^. These, along with numerical solutions *P*_*p*_ and **u**_*p*_ give the solution for the full system. Note that since the velocity boundary conditions for *P*_*p*_ and **u**_*p*_ are **0**, then Eqns. (35) and (36) indicate the entirety of the surface velocities.

## IV. THE TRANSFER MATRICES

### A. Dependencies on wavevector and other parameters

To understand how the geometry (parameters *h* and *H*) and intensive material parameters (viscosities *η*_*B,L*_) influence how surface force is converted to surface velocities, we explore how transfer matrices depend on the wavevector *k*, the thickness of the slab *h*, and other parameters. Consider Eqns. (31) and (32). Note that each of ***τ, τ***^*t*^, and ***τ***^*b*^ have the form:

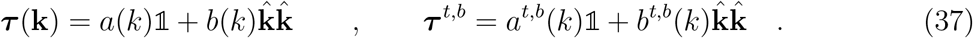

From Eq. (37), the eigenvalues and corresponding eigenvectors of ***τ*** are: *e*_1_ = *a*(*k*) + *b*(*k*), 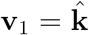 and *e*_2_ = *a*(*k*), 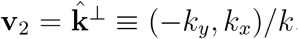. Analogous eigenvectors and eigenvalues hold for ***τ***^*t,b*^. Using the eigenvectors of ***τ***, we express the action of ***τ*** on an arbitrary vector 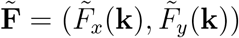 in the basis of vectors parallel and perpendicular to 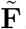, vectors 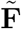 and 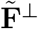:

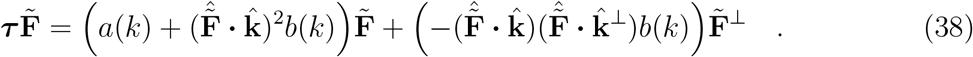

Equation (38) is derived in section (SIV A). Interpreting 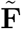 as the **k**th Fourier component of a driving force profile **F**(*x, y*), Eq. (38) indicates that the action of ***τ*** on 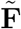, which produces a surface velocity, is proportional to 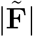 (as it should) and depends strongly and explicitly on the orientation of wavevector **k** with respect to the orientation of the driving force, evident from the factors 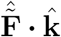 and 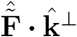. We take 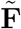 to be in the *x* direction, i.e. 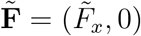, then Eq. (38) becomes:

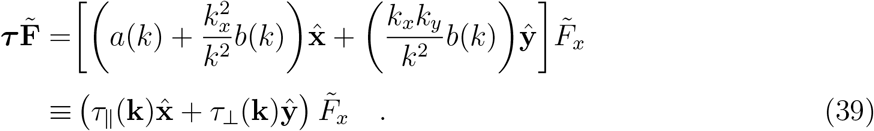

Here, we defined the coefficient of force in the *x* direction as the “parallel transfer coefficient” τ_‖_ and the coefficient of force in the *y* direction as the “perpendicular transfer coefficient” *τ*_⊥_. We assume the driving force 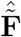 is aligned along *x* for the rest of the paper and take *k*_*x*_ and *k*_*y*_ to be positive.

Transfer functions ***τ***^*t*^ and ***τ***^*b*^ generate similar expressions to Eq. (39):

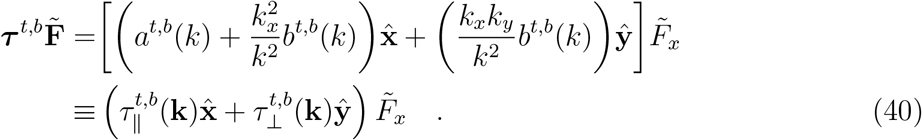

From Eqns. (39) and (40), it is evident that the force profile must vary in both *x* and *y* to produce a velocity perpendicular to the force. For example, a force profile *F*_*x*_(*x, y*) = sin(*k*_*x*_*x*) or *F*_*x*_(*x, y*) = sin(*k*_*y*_*y*) will produce no *y* velocity.

We explore how *a, b* and *a*^*t,b*^, *b*^*t,b*^ behave as *k* and other physical parameters are varied, and then comment on the consequences for parallel and perpendicular transfer coefficients 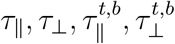.

Figure 3 shows *a*(*k*), *a*^*t,b*^(*k*) (shades of red) and *b*(*k*), *b*^*t,b*^(*k*) (shades of brown) as functions of *k* using the dimensionless (rescaled using *η* and _0_) base parameters *η*_*B*_ = *η*_*L*_ = 1, *H* = 0.5, *h* = 2, *L* = 20. Here, each graph varies one of the parameters *h, H, η*_*B*_, or *η*_*L*_ higher or lower by numerical factors and plots the result in a lighter or darker shade, respectively. The range between which each of these parameters approaches 0 and ∞ are shaded. The base parameters are chosen to mimic physical parameters in the fly embryo, where, if the length scale *ℓ*_0_, is taken as the radius of a typical cell, then the thickness of the fluid layer *H* is typically 0.1 *ℓ*_0_ to 2 *ℓ*_0_, the thickness of the epithelium *h* is typically 0.5 *ℓ*_0_ to 10 *ℓ*_0_, and the viscosities of the fluid bath and fluid layer (*η*_*B*_ and *η*_*L*_) are on the order of *η*. Figure S1 shows the same functions considered as functions of *h* using the same base parameters as Fig. 3.

**FIG. 3.**
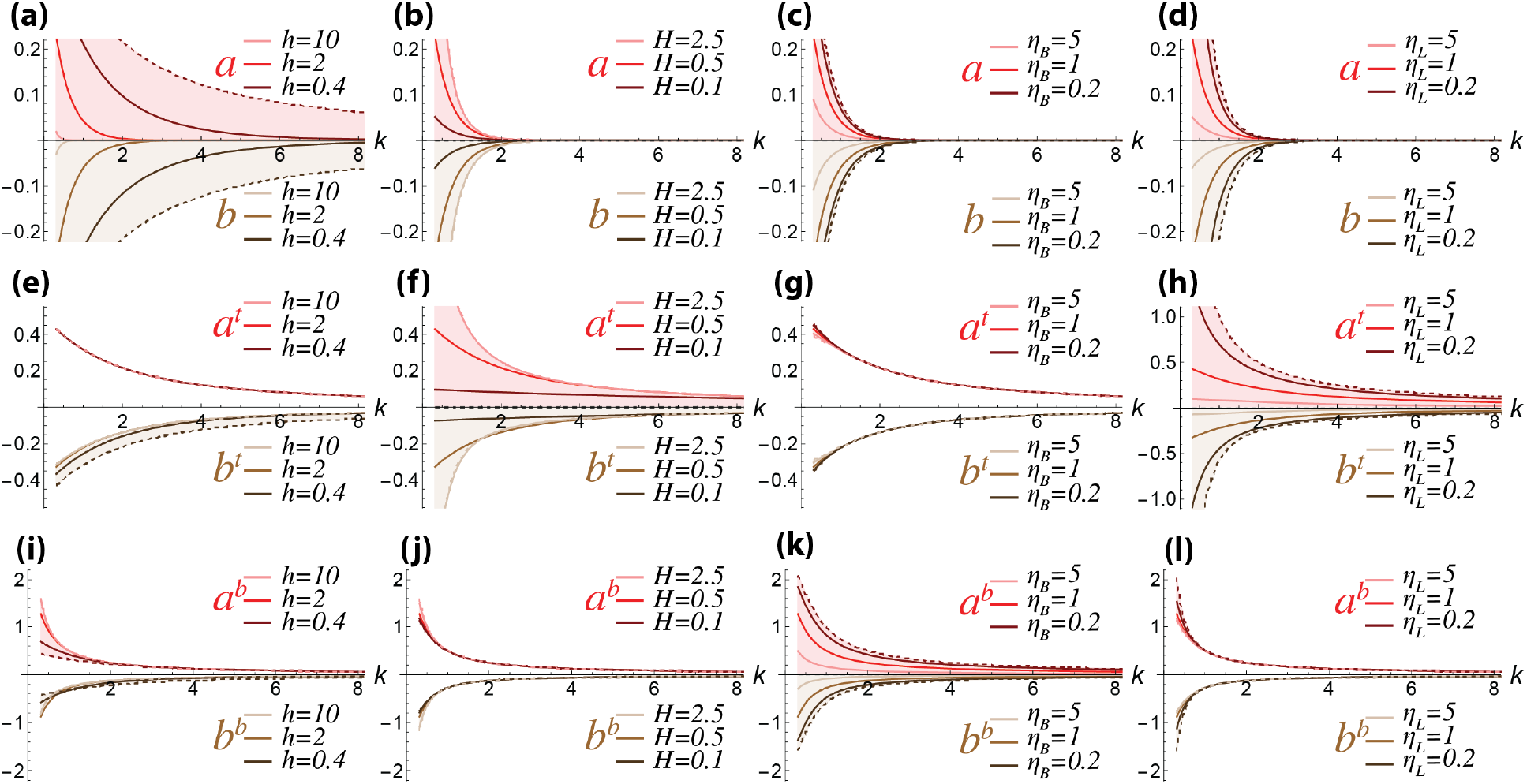
Matrix elements of ***τ, τ***^*t,b*^. Quantities *a, a*^*t,b*^ are graphed in red shades and quantities *b, b*^*t,b*^ are graphed in brown shades as functions of *k*. Base parameters are *η* = *η*_*B*_ = *η*_*L*_ = 1, *H* = 0.5, *h* = 1, *L* = 20. Each graph varies one of these parameters and plots the result, with the shaded regions denoting the 0 to ∞ limits of each of these parameters. The 0 and ∞ limits are indicated by a dark or light dotted line, respectively. (a-d) *a*(*k*) and *b*(*k*); (e-h) *a*^*t*^(*k*) and *b*^*t*^(*k*); (i-l) *a*^*b*^(*k*) and 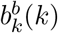.

From Figs. 3 and S1 and analysis, we make observations about the behavior of the matrix elements *a, b* and *a*^*t,b*^, *b*^*t,b*^ of the transfer matrices ***τ*** and ***τ***^*t,b*^.

I. **Perpendicular transfer coefficients** *τ*, 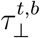 **are *always* negative**. This is because the quantities *b, b*^*t,b*^ satisfy *b, b*^*t,b*^ *<* 0 as plotted in Figs. 3 and S1 and shown analytically in section (SIV C). Since *b, b*^*t,b*^ *<* 0 and *k*_*x*_ and *k*_*y*_ are positive, then it follows from Eqns. (39) and (40) that *τ*, 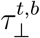 are negative. This means that if the surface force is compressional (extensional) in the *x* direction, then the velocity produced on *any* surface in the *y* direction will *always* be extensional (compressional). Notably, Figs. 3 and S1 and section (SIV C) also show that *a, a*^*t,b*^ *>* 0.
II. **The quantities** 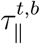 **satisfy** 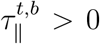. The sums *a*^*t,b*^ + *b*^*t,b*^ satisfy *a*^*t,b*^ + *b*^*t,b*^ *>* 0 from Figs. S2 and S3; these are also shown analytically in section (SIV C). From Eq. (40), the expressions *a*^*t,b*^ + *b*^*t,b*^ *>* 0 are lower bounds to 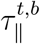 since *b*^*t,b*^ *<* 0. A positive lower bound on 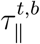 means that 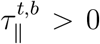, so when forces transfer to velocities on the same surface, the velocities parallel to the force are in the same direction as the force.
III. **The quantity** τ_‖_ **is positive or negative depending on k**. The sum *a* + *b* satisfies *a*+*b <* 0 from Figs. S2 and S3; these are also shown analytically in section (SIV C). Equation (39) indicates that *a* + *b <* 0 is a lower bound to τ_‖_ since *b <* 0. A negative lower bound on τ_‖_ with *a >* 0 means that *τ*_I_ may be positive or negative depending on the wavevector **k**. In particular, the transfer coefficient of surface force to surface velocity generates *velocity reversal* (*τ*_‖_ *<* 0) for some wavevectors in which a compressional (extensional) force in the *x* direction on a surface can generate an extensional (compressional) velocity in the *x* direction on the opposite surface. For example, if *F*_*x*_(*x, y*) = sin(*k*_*x*_*x*), then *τ*_‖_ = *a* + *b <* 0 and velocities will be reversed on the opposite surface. We discuss this in detail in the next section.
IV. **Transfer matrix elements decrease with** *k*. This is intuitive. Figure 3 shows that all quantities *a, b, a*^*t,b*^, *b*^*t,b*^ decrease in magnitude as a function of *k*, meaning that high spatial frequency modes are harder to transfer.
V. **The quantities** *a*^*t*^ **and** *b*^*t*^ **are insensitive to** *h* **and** *η*_*B*_**; the quantities** *a*^*b*^ **and** *b*^*b*^ **are insensitive to** *H* **and** *η*_*L*_. These observations are evident from Fig. 3(e, g, j, l). In Fig. 3(e, g), the full range of *a*^*t*^ and *b*^*t*^ for all values of *h* and *η*_*B*_ are shaded, and we see that the shaded region is small, meaning that even when *h* and *η*_*B*_ are varied widely, the values of *a*^*t*^ and *b*^*t*^ do not change significantly. This makes sense, as *h* and *η*_*B*_ are parameters relevant to physical elements that are farther from the top surface.

Similarly, from Fig. 3(j,l), the full range of *a*^*b*^ and *b*^*b*^ for all values of *H* and *η*_*L*_ are shaded, and we see that even when *H* and *η*_*L*_ are varied widely, the values of *a*^*b*^ and *b*^*b*^ do not change significantly. This again makes sense, as *H* and *η*_*L*_ are parameters relevant to the fluid layer, which is farther from the bottom surface.

#### 1. Parallel and perpendicular transfer coefficients

Given a driving force *F*_*x*_, we show how the parallel and perpendicular transfer coefficients *τ*_‖_, *τ*_⊥_, 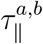, and 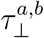 depend on the wavevector **k** of the force. Figure 4 plots *τ*_‖_ and *τ*_⊥_ for *h* = 2 *ℓ*_0_ in which the cell height and width are roughly equal (with *η*_*L,B*_ = 1, *H* = 0.5, *L* = 20). Figures S4 and S5 show similar results for *h* = 0.5, a shorter cell, and *h* = 8, a taller cell most resembling the dimensions of a *Drosophila* cell in the ventral furrow and convergent-extension phases. Consider the results for *τ*_I_. There exists a region in the lower right where *τ*_‖_ *<* 0: this is the region of velocity reversal. Since 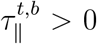 everywhere, then Fig. 4 confirms that there is never velocity reversal in the parallel direction when forces transfer to velocities on the same surface. We also see that *every* perpendicular transfer is negative, confirming that compressional (extensional) forces in *x* always lead to extensional (compressional) forces in *y*. From Fig. 4(b), we observe that large parallel transfer coefficients are generically located in the upper left in **k** space, while large perpendicular transfer coefficients are generically located near the diagonal. Hence the modes of driving that maximize magnitudes of parallel and perpendicular transfer are different.

**FIG. 4.**
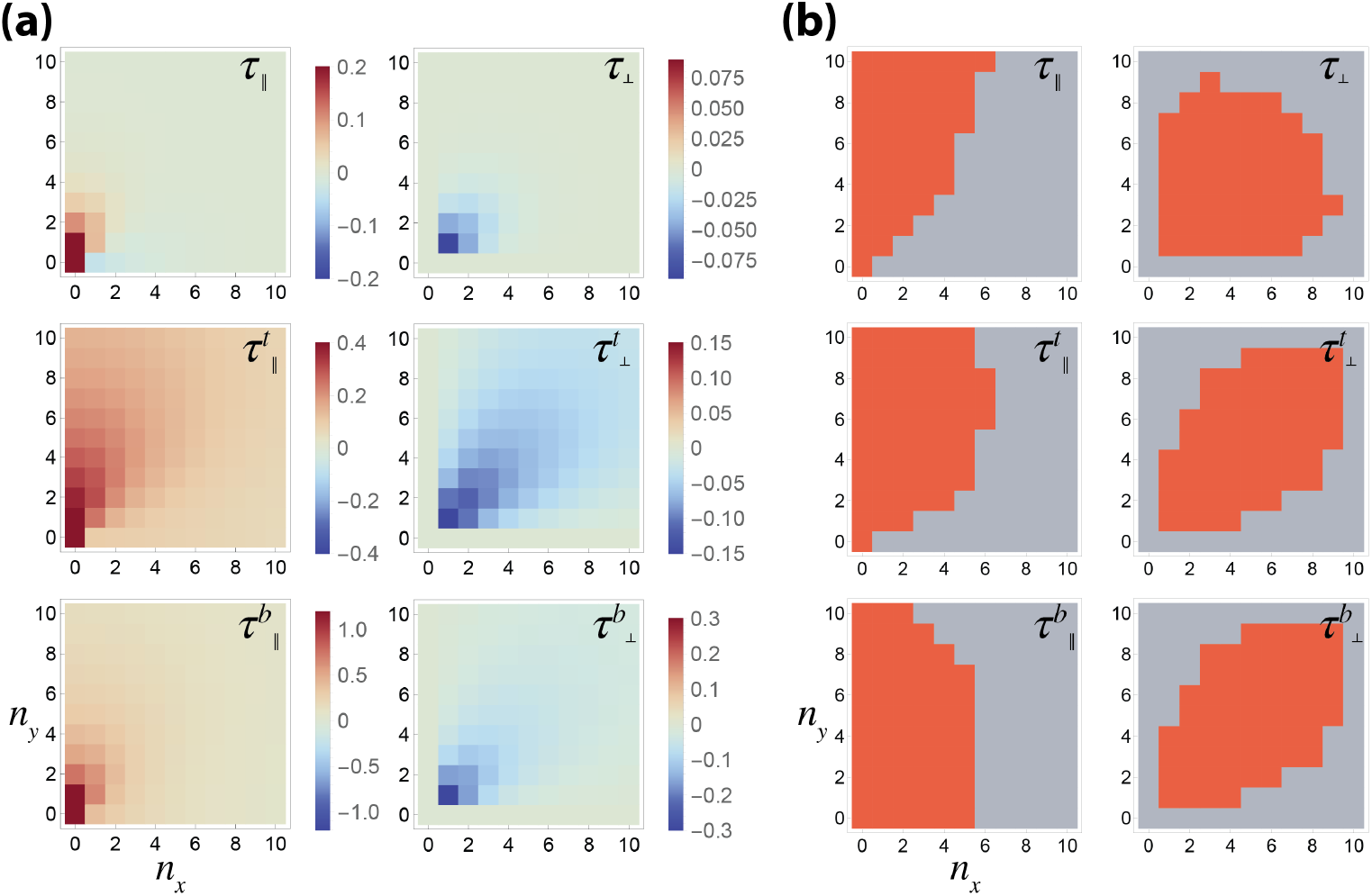
Dependence of parallel and perpendicular transfer coefficients on wavevectors for *h* = 2.0*ℓ*_0_. (a) Coefficients of parallel and perpendicular transfer *τ*_‖_, *τ*_⊥_, 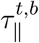, and 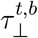 plotted (color scale) as functions of 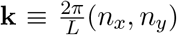. (b) Wavevectors in which |*τ*_‖_ |, |*τ*_⊥_|, 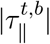, and 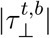 are larger than their median values are highlighted in dark orange; for *τ*_‖_, only non-negative values greater than the median are highlighted.

### B. Velocity reversal when transferring to the opposite surface

If our biological problem is to find spatial profiles of driving such that top and bottom surfaces move together, then surface force modes with velocity matching (*τ*_‖_ *>* 0) are preferred over those with velocity reversal (*τ*_‖_ *<* 0). From Eq. (39), velocity reversal occurs whenever *k*_*x*_ is sufficiently close to *k*. Figure 5(a) shows wavevectors of the driving force that produce velocity matching and reversal: generally, wavevectors with small *k*_*x*_ and large *k*_*y*_ (the upper left in **k** space) will produce velocity matched solutions; these modes have long wavelengths in *x* and short wavelengths in *y*. If the slab thickness is increased as in Fig. 5(a), then the range of **k** that produces velocity matched solutions becomes smaller.

**FIG. 5.**
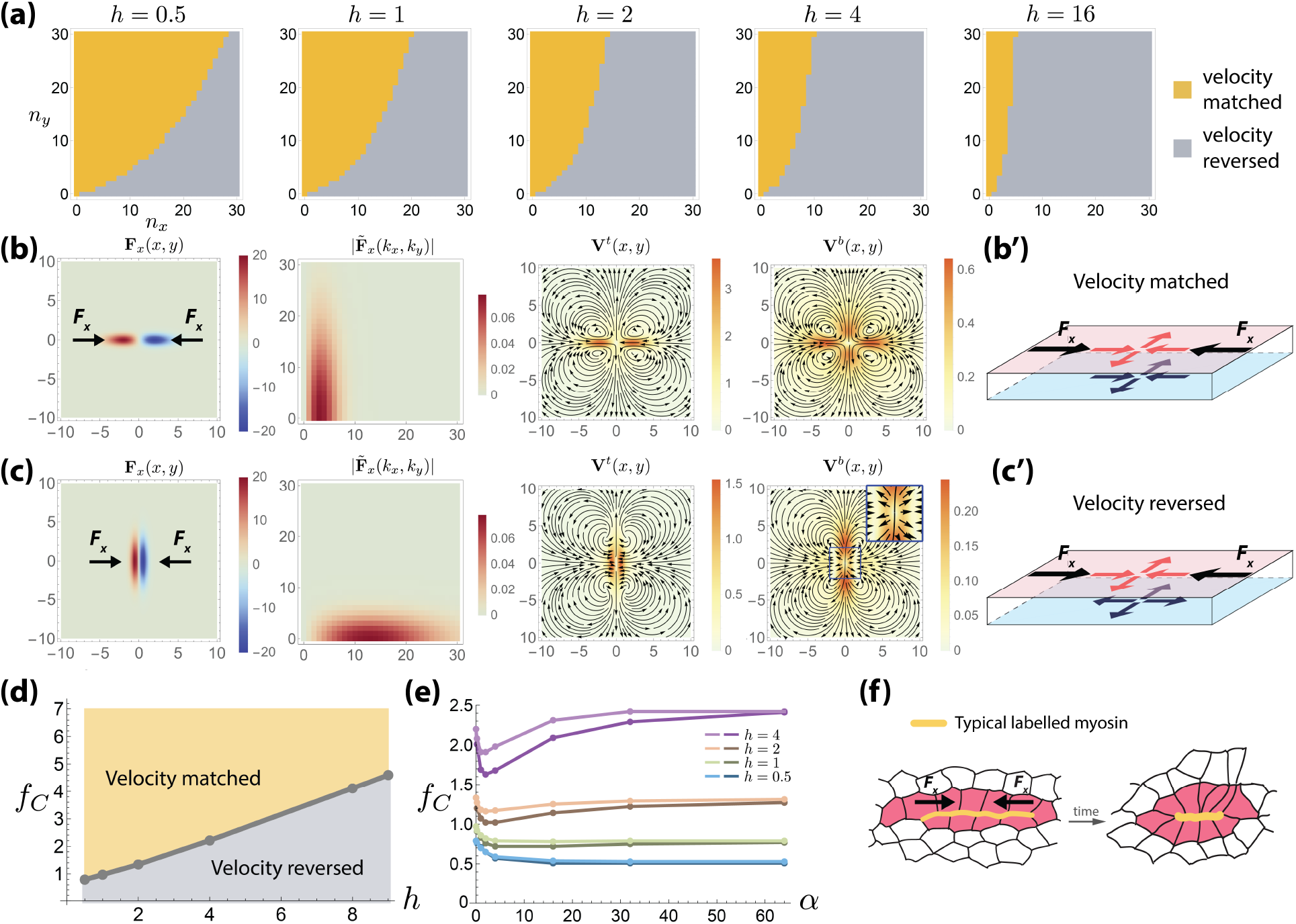
Velocity matching vs. velocity reversal. (a) Wavevectors that produce velocity matched (yellow) versus velocity reversed (gray) solutions for various *h*. As *h* increases, the range of **k** producing velocity matched solutions becomes smaller. (b) Velocity matching when *F*_*x*_(*x, y*) consists mainly of modes from the upper left in **k** space. (b’) Schematic of velocity matching. (c) Velocity reversal when *F*_*x*_(*x, y*) consists mainly of modes from the lower right in **k** space. (c’) Schematic of velocity reversal. (d) When *α* = 0, with increasing *h*, larger *f* is required for velocity matched solutions; *f*_*C*_ is the crossover value of *f*. (e) Taking *α >* 0, the crossover *f*_*C*_ for velocity matched versus velocity reversed solutions is shown for different *h* and *A*. For each *h*, light and dark lines indicate *A* = 10 and *A* = 20, respectively; velocity matched (reversed) solutions exist above (below) each line. F) Schematic of elongated myosin profile (interpreted as active stress profile) during convergent extension in *Drosophila*.

As an illustration of velocity reversal, consider the force profile **F**(*x, y*) = (*F*_*x*_(*x, y*), 0) with

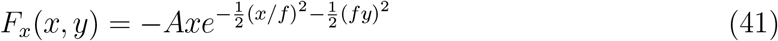

applied to the top surface. Here *f* is a numerical factor controlling the profile shape. For *h* = 1 (with *η*_*L,B*_ = 1, *H* = 0.5, *L* = 40), Figs. 5(b) and (c) examine the cases *f* = 2 and *f* = 0.5, respectively. The modal content of the first case (*f* = 2 in Fig. 5(b)) sits largely in the velocity matched region of Fig. 5(a) (second panel) and indeed **V**^*t*^ and **V**^*b*^ (third and fourth panels of (b)) both show similar surface hyperbolic flows. Conversely, the modal content of the second case (*f* = 0.5 in Fig. 5(c)) sits primarily in the velocity reversed region, and in the third and fourth panels of Fig. 5(c), we see a transition from a hyperbolic straining surface flow in **V**^*t*^ to an outwards source-like surface flow in **V**^*b*^ near the origin.

During convergent extension of the embryonic fly epithelium, labelling has revealed elongated regions enriched in myosin (“myosin cables”) aligned in the direction of convergence [6, 31] (diagrammed in Fig. 5(f)). Since these cables likely exert force profiles similar to that in Fig. 5(b), with modes largely in the velocity matched regions, we speculate that the *Drosophila* embryo may be employing a strategy of velocity matching. Since *Drosophila* tissue at this stage is very thick (around *h* = 8 − 10), very large shape factors *f* are required to achieve velocity matching (Fig. 5(a,d)), and indeed the shape of myosin cables correspond to very large *f*.

#### 1. Dependence of velocity reversal on tissue thickness and viscoelastic strength

If we consider the tissue to be purely viscous (*α* = 0), then as the tissue thickness *h* is increased, the value of the shape factor *f* from Eq. (41) needs also to be increased in order to achieve velocity matched solutions. The crossover value *f*_*C*_ of *f* between velocity matched and velocity reversed solutions is shown in Fig. 5(d).

We additionally examine how the crossover between velocity matched and velocity reversed solutions changes when the viscoelastic strength *α*, the tissue thickness *h*, and the amplitude of the driving force *A* are changed. Figure 5(e) shows *f*_*C*_ as a function of *α* for various values of *h* and *A*. There, values of *f* above (below) each line indicate force profiles creating velocity matched (reversed) solutions. We find that for large values of *h*, e.g. *h >* 0.5, the crossover *f*_*C*_ changes non-monotonically with *α*, i.e. there is an optimal, middle range of *α* for which the velocity matched parameter region is largest. The intuition is the following: as *α* is increased, the tissue becomes more elastic and surfaces tend to move together, making it easier for velocity matching; on the other hand, increased elasticity also means that the material is more resistant to compression, making it harder for hyperbolic flows to appear near the origin and harder for velocity matching. Hence, intermediate values of *α* provide the largest velocity matched parameter regions.

### C. Velocity response to periodic driving with full Oldroyd-B model

The velocity response to periodic driving in the full Stokes Oldroyd-B model is given numerically and depends on the wavevector **k** of the driving as well as on the (non-dimensional) temporal driving frequency *ω* (units of 1*/τ*_*p*_), proportional to the Deborah number *De* ≡ *ωτ*_*p*_*/*(2*π*). The driving force is applied to the top surface and has the form **F**(*x, y, t*) = (*F*_*x*_(*x, y, t*), 0) with:

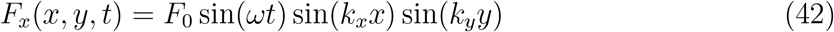

where *F*_*x*_(*x, y, t*) = −*F*_0_ sin(*ωt*) sin(*k*_*x*_*x*) if *k*_*y*_ = 0 and *F*_*x*_(*x, y, t*) = −*F*_0_ sin(*ωt*) sin(*k*_*y*_*y*) if *k*_*x*_ = 0.

The driving force is specified with a given wavevector (*k*_*x*_, *k*_*y*_) in the direction *x* on the top surface. However, the velocity response will consist, in general, of both *x* and *y* velocities, velocities on both surfaces, and, in particular, since *α* ≠ 0, velocities with components in modes other than the driving mode. In Fig. 6, we consider the component of the velocity response in the same mode as driving. In that mode, top and bottom velocities in steady state oscillation take the form:

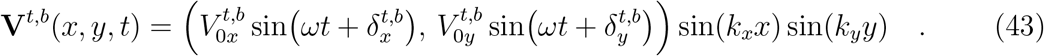

Figure 6 shows the magnitude of velocity-force responses 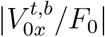 and 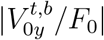 for driving wavevectors 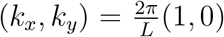 (a pure compression mode, Fig. 6(a)), 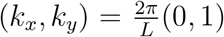 (a pure shear mode, Fig. 6(c)), and 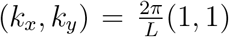 (Fig. 6(c)), simulated with *η*_*L,B*_ = 1, *h* = 2, *H* = 0.5, *L* = 20. In the pure compression and pure shear modes, the velocity response in the *y* direction is 0. When the viscoelastic contribution *α* is large enough, e.g. *α* = 4 or higher in Fig. 6, the velocity response on the bottom side 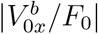 and 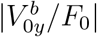 exhibit a maximum at certain values of the Deborah number. This implies that there are certain frequencies of driving for which transfer of force to the opposite surface is optimal for the velocity modes in the same mode as the force. The angular responses 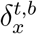 and 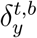 for the same driving forces are shown in Fig. S6.

**FIG. 6.**
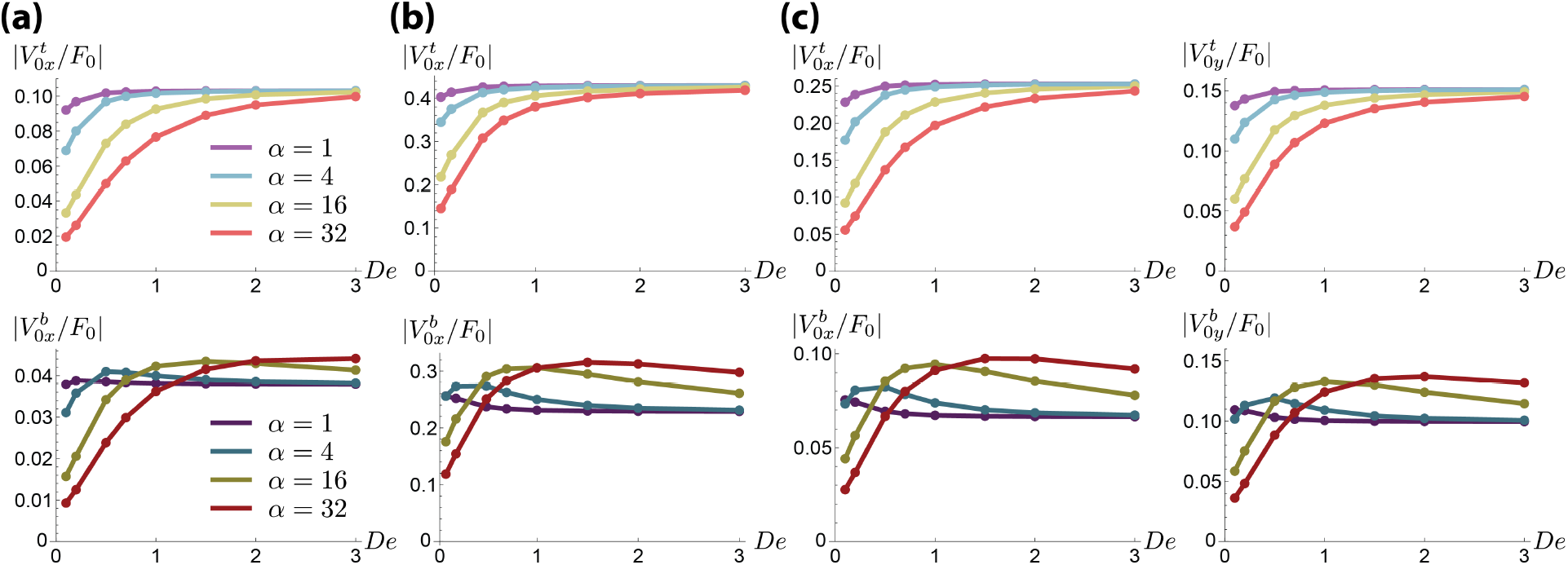
Velocity response as a function of Deborah number for the driving force in Eq. (42) with 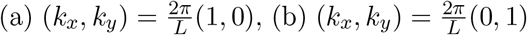, and 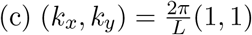. Color key for (b) and (c) are same as in (a).

## V. 3D CELL DEFORMATION

A useful feature of our model is the ability to track material elements while simulating surface forces that resemble those from live experiments. If we assume that cell membranes are not force producing objects, but merely carried by the flow of the tissue material, a roughly valid assumption for very early stages of the fly embryo [26], then the movement of markers in our velocity solutions would approximate that of cell membranes. Hence, instead of modeling membranes, we track them as material surfaces. In the following, we show two examples of quantifying cell deformation dynamics using tracked membranes. Additionally by tracking cells, we specify surface forces such that they remain localized to a particular cell as that cell moves. We find results that are suggestive in two events during *Drosophila* development: ventral furrow invagination and convergent extension.

### A. Quantifying cell deformations

Figure 7 shows two illustrative examples of how cell deformation dynamics from surface forces can be simulated, quantified, and studied. In Fig. 7(a,b) and Movie S1, we apply an isotropic, contractile, traveling wave force on the top surface of the tissue that moves to the right (*η*_*L,B*_ = 1, *h* = 2, *H* = 0.5, *L* = 20). The force is given as the divergence of the stress 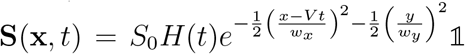 where *S*_0_ = 20, *V* = 1.0, *w*_*x*_ = 1, *w*_*y*_ = 4 (in non-dimensional units) for Fig. 7(a), and *H*(*t*) is a heaviside function that sets the force to zero when it has moved close to the edge of the simulation domain. Figure 7(b) shows the displacement of the origin point (0, 0), tracked as a function of time, for simulations with different *α* (*S*_0_ = 20, *V* = 4.0). When *α* is small, viscous response dominates, the total displacement is larger and shows no recoil; when *α* is large, elasticity plays a larger role in the response, the total displacement is smaller, and recoil from the elasticity can be observed.

**FIG. 7.**
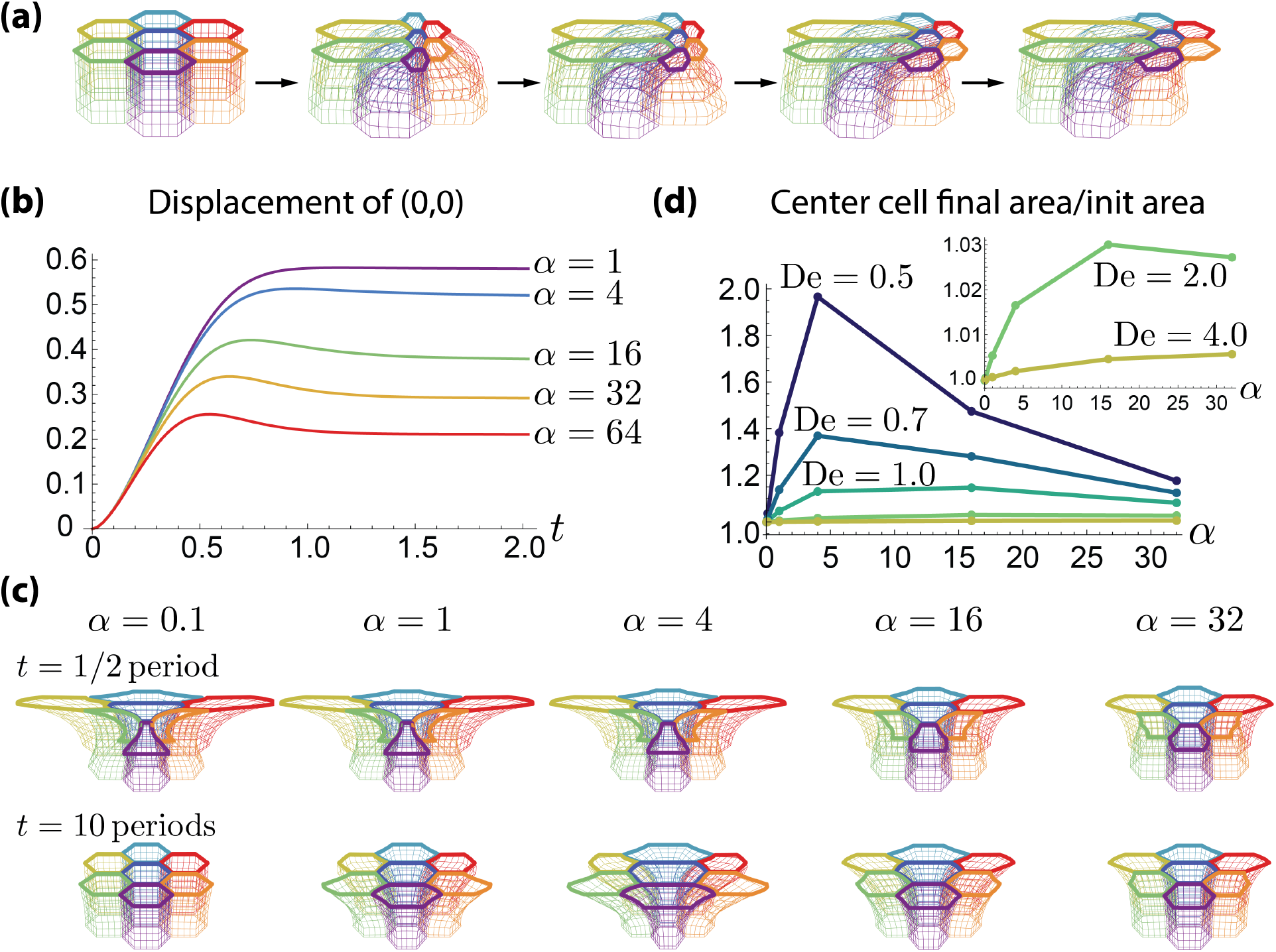
Cell deformations due to surfaces forces. (a) Time series of cell deformations from a traveling wave contractile surface force. (b) Displacement of the origin point (0, 0) tracked as a function of time for simulations with different *α*. (c) Cell shapes from an oscillatory force after half a period (top) and after 10 full periods (bottom) for different *α*. (d) Area deformation of central cell after 10 periods of oscillations; maximum deformation is achieved for different values of *α* depending on *De*.

In Fig. 7(c,d) and Movie S2, we apply the oscillatory force in Eq. 42 with *De* = 0.5, *F*_0_ = 20, and 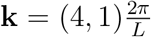 and quantify cell deformations over time. Figure 7(c) shows deformations at the timepoints of half a period and 10 periods. For small values of *α*, we see that the cells are easy to deform (left panel, top) but carry no memory of past deformations (left panel, bottom), while for large *α*, the cells are harder to deform (right panel, top) but carry more memory (right panel, bottom), and intermediate values of *α* show complex combinations of these limits. Figure 7(d) quantifies the change in the area of the central cell after 10 periods of oscillation when the net force over time is 0. We show that depending on the Deborah number of the driving force, the maximum area deformation is achieved at different values of *α*. The two examples in Figure 7 indicate the interesting and important role of viscoelastic strength in cell deformation dynamics and the range of cell shape phenomena that can be studied.

### B. Insight into ventral furrow formation

An important event in *Drosophila* development is ventral furrow formation, occurring when a group of cells invaginate from the surface to the inside of the embryo. The invagination is preceded by many cells undergoing “pulsatile apical constriction” [4] meaning that the invaginating cells actively constrict their apical (top) surfaces in a temporally periodic manner. This process is known to be driven by myosin localized to the apical surface. After 2 to 4 cycles of apical constriction, the epithelium starts to invaginate. We model these periodically constricting cells to understand the physics of invagination.

#### 1. Estimating forces and tissue properties

To model apical constriction, we applied a local, convergent force periodically in time to a cell centered at the origin (Fig. 8(a) and Movie S3). The applied force is computed as the divergence of a 2D, isotropic, radially symmetric active stress tensor **S** whose slice at *y* = 0 is shown in Fig. 8(A)). The forces from **S** model contractile forces from myosin. The tensor **S** is given by:

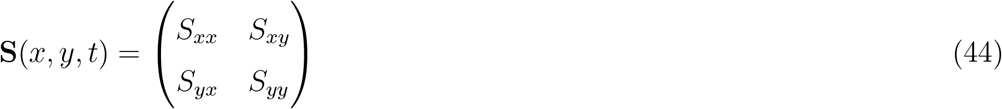

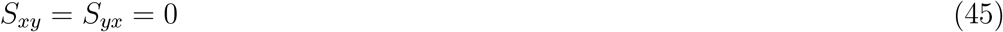

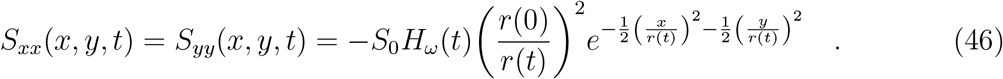

This is a simple model of force feedback since *r*(*t*) corresponds to the radius of the central cell as it is tracked in time. Here *r*(0) = *ℓ*_0_ is the radius of the cell at time 0 (rescaled so *ℓ*_0_ ≡ 1), and *S*_0_ is the amplitude of the stress. The function *H*_*ω*_(*t*) is a Heaviside function with *H*(*ω*) = 1 when sin(*ωt*) *>* 1 and *H*(*ω*) = 0 when sin(*ωt*) *<* 1, simulating the pulsatile stress being on and off at frequency *ω*. The form of **S** is chosen such that the integrals of *S*_*xx*_ and *S*_*yy*_ are constant, as 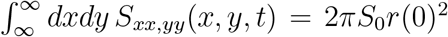. This simulates that the total amount of myosin in a cell’s apical surface is constant in time.

**FIG. 8.**
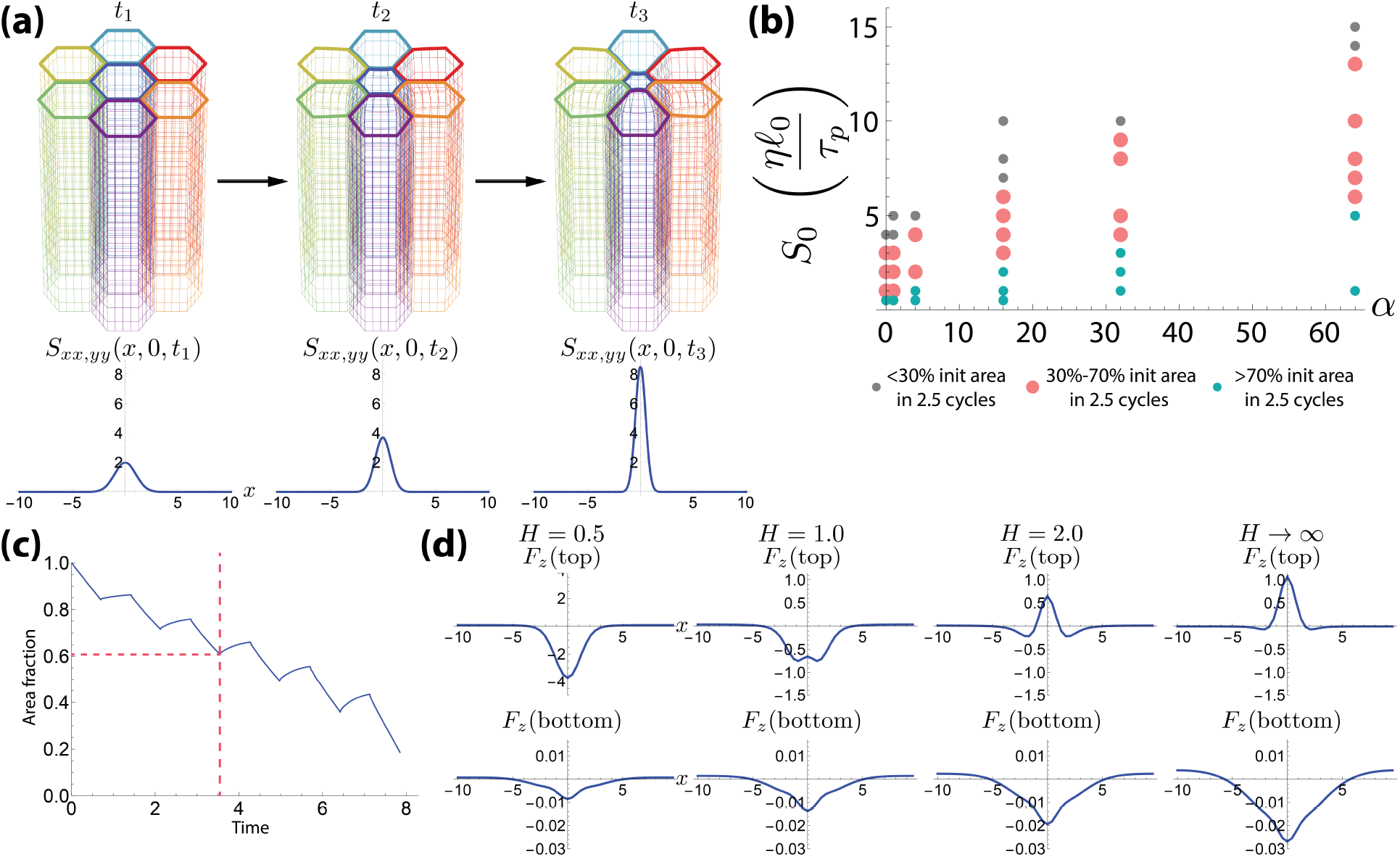
Simulation of apical constriction. (a) Frames from apical constriction simulation (top); *S*_*xx,yy*_(*x*, 0, *t*) at the timepoints of the frames. (b) Diagram of parameter space *α* and *S*_0_; pink dots indicate simulations that agree with experiments. (c) Constricting cell’s area as a fraction of its initial area during apical constriction simulation; dotted lines indicate 2.5 cycles of constriction-relaxation and corresponding area fraction. (d) The *z*-directional force *F*_*z*_ (*y* = 0 slice) on the tissue’s top and bottom surface for various thickness of the fluid layer *H*.

During ventral furrow formation, cells undergo approximately 2.5 cycles of constriction-relaxation before the epithelium starts moving out-of-plane [4]. Therefore, our simulations of apical constriction in flat epithelial tissue can be compared to experiments for the first 2.5 simulated cycles of constriction-relaxation (dotted line in Fig. 8(c)). Experiments in [4] estimate that cell apical areas shrink to approximately 50% their initial values during these 2.5 cycles, taking 2 to 3.5 minutes. Using that *τ*_*p*_ ≈ 1 minute [28], we simulated apical constriction in single cells using length scales relevant to *Drosophila* (*h* = 8*ℓ*_0_, *H* = 0.5*ℓ*_0_) with periods of forcing between 1–2*τ*_*p*_ (corresponding to 2.5 cycles of constriction-relaxation in 2–3.5 minutes). After simulating 2.5 cycles, we identified parameters *S*_0_ and *α* for which cell areas decreased to 30% to 70% (approximately 50%) their original values using *η*_*B*_ = *η*_*L*_ = 1(*η*) (Fig. 8(b)).

Our simulations show that as the viscoelastic strength *α* is increased, the amplitude of stress must also increase to achieve the same amount of area deformation in same time (Fig. 8(b)); this makes sense. Since the perivitelline fluid is poorly understood, its viscosity *η*_*L*_ is highly uncertain. Authors [32] have proposed that *η*_*L*_ could be 0.1*η* or lower. Hence we present parameter space estimation with *η*_*L*_ = 0.1 in Figure S7 where we find that the required *S*_0_ to fit experiments is decreased to roughly half of the *η*_*L*_ = 1 case.

Since dimensionless force densities are defined with units *η/τ*_*p*_, then, taking *η* as approximately 1000 times water viscosity and *τ*_*p*_ = 60 s, we have that 1 on the dimensionless force density scale is equivalent to 1/60 Pa. Hence, the range of *S*_0_ = 1–15 *ηℓ*_0_*/τ*_*p*_ that is appropriate for various values of *α* (Fig. 8(b)) correspond to force densities of approximately 1*/*60 to 1*/*4 Pa applied to the apical surface.

#### 2. Insight into epithelial invagination

Some models for ventral furrow invagination argued that by actively decreasing their apical areas, cells become individually wedge shaped; this shape could only be accommodated by an overall invaginated shape for the epithelium, and hence the epithelium becomes invaginated. Others argue that invagination occurs as a result of instability: a compressional force is created when cells apically constrict; the epithelium is unstable to compression and buckles.

Our model elucidates the phenomena of invagination quantitatively in the context of the tissue being surrounded by incompressible fluids in three dimensions. When the active stress in Eq. (46) is applied in simulations, we calculate the force densities in the *z* direction exerted on the top (*h* = 0) and bottom (*h* = −*z*) surfaces of the tissue due to both the directly applied force and forces from its surrounding environment.

When we calculate (section SII C) the *z*-directional forces that are exerted on the top and bottom surfaces of the tissue (*y* = 0 slices in Fig. 8(d)) for the timepoint at the end of constriction), we find that in our *Drosophila*-inspired geometry, negative *z*-directional forces are indeed exerted there (Fig. 8(d) left). However, if we assume that the tissue sits in an infinite bath, that is, if we take the limit *H* → ∞, then we see that the *z*-force exerted on the tissue’s top surface is positive, while the *z*-force exerted on the tissue’s bottom surface is negative (Fig. 8(d) right). This means that if we eliminate the constraint that the tissue remains flat, then convergent active stresses on the tissue’s surface would act to make it locally thicker instead of causing it to invaginate. If we consider intermediate cases where *H <* ∞ but is larger than its typical value in *Drosophila* (Fig. 8(d) middle panels), corresponding an enlarged space for the perivitelline fluid, then the *z*-forces on the top and bottom surfaces of the tissue also achieve intermediate values, showing that there exists a continuum of situations between tissue invagination and tissue thickening tuned by the parameters of the surrounding fluids and boundary geometry.

Our results show that it is the presence of the hard wall (vitelline membrane) in combination with the incompressible fluid layer in region *L* (perivitelline fluid) that leads the system to find a 3D solution that necessarily contains a downward force on both the tissue’s top and bottom surfaces when *H* = 0.5. Intuitively, the convergent stress, because it is applied at the interface of regions *L* and *S*, drives the material in both the fluid layer and the tissue toward (*x, y*) = (0, 0). Since these fluids are incompressible and constrained by the wall, both fluids exert a force on the other. Thus, whether or not the tissue invaginates is determined by which fluid pushes more strongly into the other. Another demonstration of this concept is in Fig. S8, in which as the viscosity of the fluid layer *η*_*L*_ is decreased, the fluid layer pushes less strongly into the tissue, and the resultant *z*-force no longer causes the tissue to invaginate. Our explanation for invagination is different from the notion of buckling because in this case, an explicit *z*-directional force is produced by the competition between the fluid layer and the tissue; there is no instability. For parameters of the *Drosophila* embryo, the convergent active stress causes invagination, but one may find other systems where instead of invagination, a convergent active stress causes the tissue to locally thicken.

### C. Convergent extension

Recent experiments have found that cells undergoing convergent-extension in both *Drosophila* [5] and vertebrates such as *Xenopus* [8, 33, 34] and mouse [35] exert active forces on *both* their apical and basal surfaces to create T1 transitions, a cell-level movement required for convergent-extension. We simulated T1 transitions to understand whether they can be driven *exclusively* by surface forces. We used non-zero **F**^*t*^ and **F**^*b*^ that are computed as the divergence of a 2D stress tensor **S** with components:

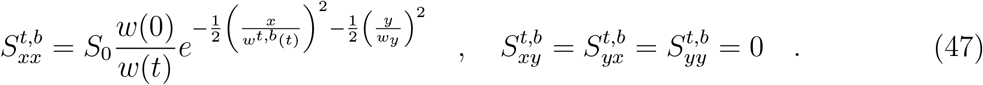

Here, *w*^*t,b*^(*t*) is the length of the junction executing the T1 transition (on either the top or bottom of the epithelium) as a function of time, *S*_0_ is the amplitude of the stress, and *w*_*y*_ is a constant parameter that specifies a delta-like profile for the stress in the *y*-direction; again, this is a simple model of force feedback.

Our simulations show that short cells (*h* = 0.5) are able to execute a T1 transition (Fig. 9(a) and Movie S4) throughout the height of the cells (other parameters *η*_*L,B*_ = 1, *H* = 0.5, *α* = 32, *S*_0_ = 5, *w*_*y*_ = 0.2, *L* = 20). However, tall cells, such as those in the fly embryo (*h* = 8), are not able to propagate the topological change from a T1 transition to the interior of the tissue: while both top and bottom surfaces of cells exchange neighbors, away from the surface, cells remain adjacent to their original neighbors (Fig. 9(d) and Movie S7). Even for cells that are relatively short (*h* = 1, 2, Fig. 9(b,c) and Movies S5-6), we see that the middle of cells cannot come together.

**FIG. 9.**
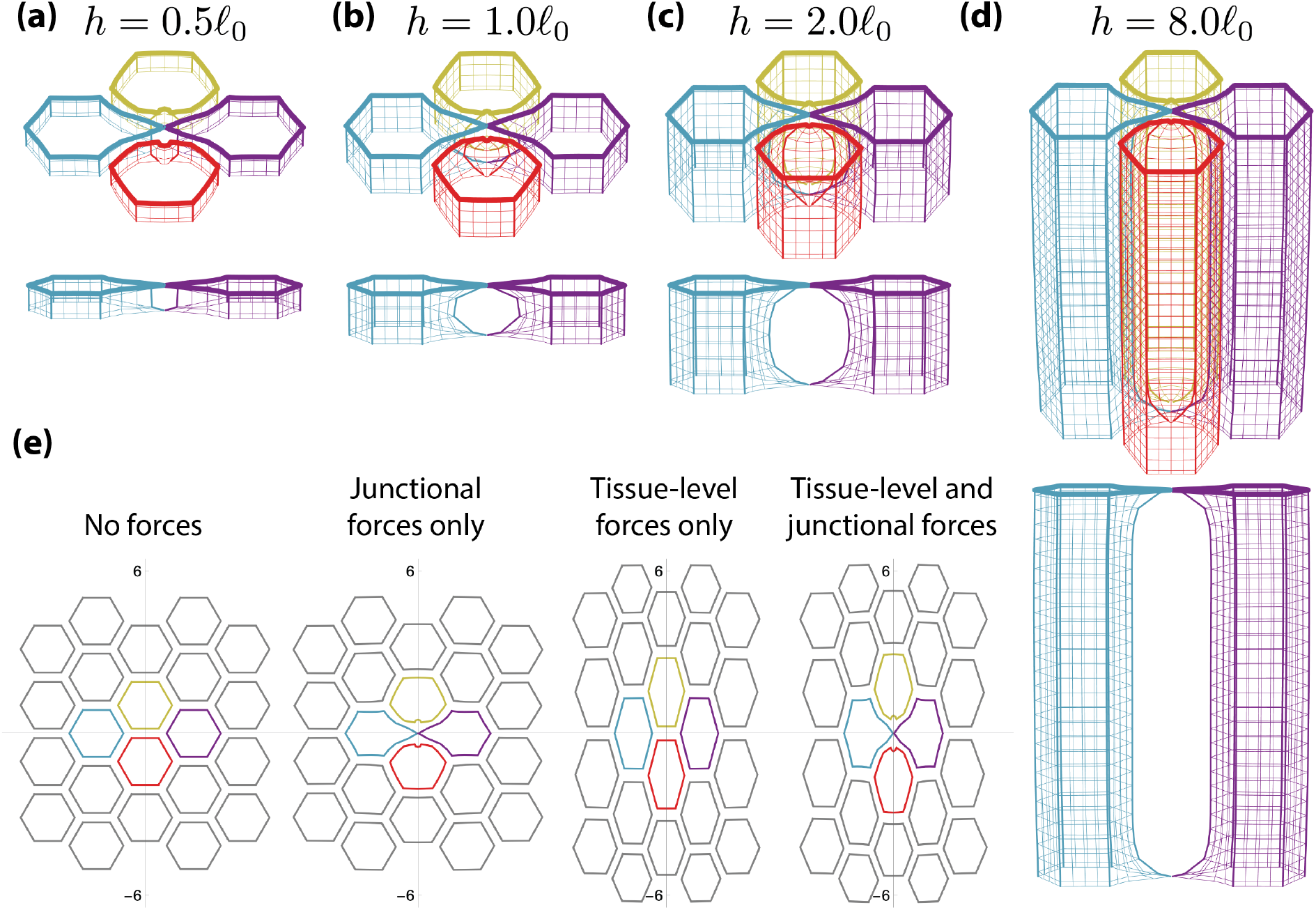
Simulation of T1 transition. (a) Short cells are able to execute a T1 transition through the full height of the cell. (b-d) Cells with intermediate and large height cannot execute T1 transitions through the full height of the cell. (e) Forces on the length scales of cell junctions are required for T1 transitions. Images show apical (top surface) cell boundaries for **F**^*t*^ = **F**^*b*^ = **0** (first panel), **F**^*t*^ = **F**^*b*^ = ∇ · **S** (Eq. 47) (second panel), 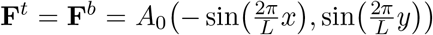 (third panel), and 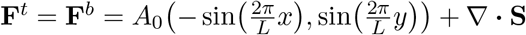 (fourth panel).

Another finding from our simulations is that forces at the length scales of cell junctions are required for T1 transitions (Fig. 9(e) second and fourth panels): overall force applied at the tissue-level (Fig. 9(e) third panel) cannot create T1 transitions even in very short cells (*h* = 0.5).

To note, authors in [5] showed that active forces from the basal side of the epithelium extend laterally beyond the basal surface by roughly 5–10 *μ*m; we do not model this. However, noting that the gap between cells in Fig. 9(d) is very large, our prediction holds that for tall cells such as those in the *Drosophila* germ band, cells would require a different mechanism to execute a T1 transition through their full height. Hence, while surface forces exclusively from the apical and basal surfaces of tall epithelia are sufficient create T1 transitions near the tissue surface, they are not sufficient to create T1 transitions for the full height of the cell. Additional forces or biological mechanisms are needed to “zip” together the remainder of the cells’ lateral sides. The biology and the mechanics of these other mechanisms may involve other molecules and are not well-understood, suggesting future directions of research.

## VI. DISCUSSION

We addressed a need for 3D modeling in thick epithelial tissues. In particular, 2D epithelial models cannot give insight to morphogenetic processes in which the apical and basal sides of cells do not move in concert. Here, we provided a framework for modeling flat, viscoelastic tissue that takes into account tissue thickness as an important parameter. We presented analytical and numerical solutions to the system, and our mathematical framework will allow us to eventually evolve active stresses as surface populations in future extensions of the model.

We analyzed how applied surfaces forces drive surface velocities by studying transfer matrices. A key result of our analysis is the existence of velocity matched versus velocity reversed solutions on the side of the tissue opposite from the force. We found that to achieve velocity matched solutions, the modal composition of the driving force should be heavily weighed with long wavelength wavevectors in the direction parallel to the force (small *k*_*x*_) and short wavelength wavevectors in the direction perpendicular to the force (large *k*_*y*_). This pattern of driving is seen in the *Drosophila* germ band during convergent extension where authors observe “myosin cables” [6, 31]. This result hints that the *Drosophila* germ band may be employing (in addition to basal forces) a strategy of velocity matching to help cells move their apical and basal surfaces in concert as much as possible across a thick epithelium.

Because our model of epithelial tissue is fully immersed in a realistic fluid environment, we are able to calculate the forces induced by other parts of the system on the epithelium when active stresses are applied. In simulations of apical constriction, we showed that when an active constricting force is applied parallel to the surface (in *x* and *y*), an orthogonal, *z*-directional force is induced on the tissue surface from adjacent fluids; and this is the force that ultimately causes invagination. The direction and amplitude of this force depends strongly on the presence and parameters of other fluids in the system and on geometric constraints. Hence, our work suggests that future studies of morphogenesis should include analysis of how geometric constraints and fluid surroundings influence the shapes of tissues.

Using apical constriction simulations, we estimated the amplitude of surface stress created by medial myosin. These estimates are only the starting point because our simulations assumed that viscosity and viscoelastic strength are spatially constant throughout the tissue. Authors [29] have shown that cells are in fact spatially heterogeneous in viscosity and viscoelastic strength even within the same cell. An elaboration of our model would be to take *α* and *η* to be space-dependent and/or to specify an evolving field *ϕ* (**x**, *t*) corresponding to the density of viscoelastic polymers in the tissue.

One key assumption in our model is that lateral membranes are not force-generating objects. In this paper, we took advantage of the fact that this assumption is roughly valid in the early *Drosophila* embryo [26] to model events during embryogenesis. However, since others have shown that there are actomyosin forces localized to lateral membranes during EMT [36], apoptosis [37], and the start of convergent-extension in the *Drosophila* mesoderm [38], a future direction will be to take into account forces from lateral membranes that may be contributing to bulk mechanics. We here showed that forces from lateral membranes likely play a role during T1 transitions in tall cells such as those in *Drosophila*. We showed that the transfer of surface force to velocities in the tissue interior is so low, that without forces from the lateral sides of cells, T1 transitions cannot propagate through the full height of the cell, and hence additional forces, perhaps due to molecules like cadherens, are required. This is another direction of future work.

Finally, a promising direction of our work is to rigorously evolve a population of active force-generators on our tissue surface alongside the bulk material. In our mathematical framework, we have the potential to incorporate any 2D active population into our system and ask: how can these populations remodel the epithelium in 3D, and how does the motion of epithelial cells couple back to the active population? Active populations include surface actomyosin, but also collections of bacteria, or a 2D model of active cellular apical surfaces and elastic membranes; our framework can evolve these populations consistently with how the tissue and other fluids move in 3D. These populations might couple to reaction-diffusion equations such as those specifying chemical oscillations, trigger waves, or genetic circuits. For example, calcium is a molecule that transports into cells under mechanical tension and feeds back to cytoskeletal forces [39, 40]: our model can simulate cellular calcium concentrations and alter material parameters based on local values of surface stress. The coupling between 2D active populations and 3D tissue will not only provide opportunities to study rich mathematical structures in these systems but also provide quantitative model predictions to connect to experiments in a wide variety of settings that involve tissues, fluid environments, and boundaries.

## Supporting information

Sopplemental Movie 1

Sopplemental Movie 2

Sopplemental Movie 3

Sopplemental Movie 4

Sopplemental Movie 5

Sopplemental Movie 6

Sopplemental Movie 7

## References

[1] Miriam Osterfield, XinXin Du, Trudi Schüpbach, Eric Wieschaus, and Stanislav Y Shvartsman. Three-dimensional epithelial morphogenesis in the developing drosophila egg. Developmental cell, 24(4):400–410, 2013.

[2] Maria Leptin and Barbara Grunewald. Cell shape changes during gastrulation in drosophila. Development, 110(1):73–84, 1990.

[3] Dari Sweeton, Suki Parks, Michael Costa, and Eric Wieschaus. Gastrulation in drosophila: the formation of the ventral furrow and posterior midgut invaginations. Development, 112(3):775– 789, 1991.

[4] Adam C Martin, Matthias Kaschube, and Eric F Wieschaus. Pulsed contractions of an actin– myosin network drive apical constriction. Nature, 457(7228):495–499, 2009.

[5] Zijun Sun, Christopher Amourda, Murat Shagirov, Yusuke Hara, Timothy E. Saunders, and Yusuke Toyama. Basolateral protrusion and apical contraction cooperatively drive Drosophila germ-band extension. Nature Cell Biology, 19(4):375–383, 2017.

[6] Jennifer A Zallen and Eric Wieschaus. Patterned gene expression directs bipolar planar polarity in drosophila. Developmental cell, 6(3):343–355, 2004.

[7] J Todd Blankenship, Stephanie T Backovic, Justina SP Sanny, Ori Weitz, and Jennifer A Zallen. Multicellular rosette formation links planar cell polarity to tissue morphogenesis. Developmental cell, 11(4):459–470, 2006.

[8] Shinuo Weng, Robert J Huebner, and John B Wallingford. Convergent extension requires adhesion-dependent biomechanical integration of cell crawling and junction contraction. Cell Reports, 39(4):110666, 2022.

[9] Nabila Founounou, Reza Farhadifar, Giovanna M Collu, Ursula Weber, Michael J Shelley, and Marek Mlodzik. Tissue fluidity mediated by adherens junction dynamics promotes planar cell polarity-driven ommatidial rotation. Nature communications, 12(1):1–16, 2021.

[10] Reza Farhadifar, Jens-Christian Röper, Benoit Aigouy, Suzanne Eaton, and Frank Jülicher. The influence of cell mechanics, cell-cell interactions, and proliferation on epithelial packing. Current Biology, 17(24):2095–2104, 2007.

[11] Hisao Honda. Geometrical models for cells in tissues. International review of cytology, 81:191– 248, 1983.

[12] Tatsuzo Nagai, Kyozi Kawasaki, and Katsuhiro Nakamura. Vertex dynamics of twodimensional cellular patterns. Journal of the physical society of Japan, 57(7):2221–2224, 1988.

[13] Tatsuzo Nagai and Hisao Honda. A dynamic cell model for the formation of epithelial tissues. Philosophical Magazine B, 81(7):699–719, 2001.

[14] Alexander G Fletcher, Miriam Osterfield, Ruth E Baker, and Stanislav Y Shvartsman. Vertex models of epithelial morphogenesis. Biophysical journal, 106(11):2291–2304, 2014.

[15] Dirk Drasdo, R Kree, and JS McCaskill. Monte carlo approach to tissue-cell populations. Physical review E, 52(6):6635, 1995.

[16] Markus Basan, Jens Elgeti, Edouard Hannezo, Wouter-Jan Rappel, and Herbert Levine. Alignment of cellular motility forces with tissue flow as a mechanism for efficient wound healing. Proceedings of the National Academy of Sciences, 110(7):2452–2459, 2013.

[17] Hisao Honda, Masaharu Tanemura, and Tatsuzo Nagai. A three-dimensional vertex dynamics cell model of space-filling polyhedra simulating cell behavior in a cell aggregate. Journal of theoretical biology, 226(4):439–453, 2004.

[18] Mahim Misra, Basile Audoly, Ioannis G Kevrekidis, and Stanislav Y Shvartsman. Shape transformations of epithelial shells. Biophysical journal, 110(7):1670–1678, 2016.

[19] Christina Bielmeier, Silvanus Alt, Vanessa Weichselberger, Marco La Fortezza, Hartmann Harz, Frank Jülicher, Guillaume Salbreux, and Anne-Kathrin Classen. Interface contractility between differently fated cells drives cell elimination and cyst formation. Current Biology, 26(5):563–574, 2016.

[20] Edouard Hannezo, Jacques Prost, and Jean-Francois Joanny. Theory of epithelial sheet morphology in three dimensions. Proceedings of the National Academy of Sciences, 111(1):27–32, 2014.

[21] Vito Conte, José J Muñoz, and Mark Miodownik. A 3d finite element model of ventral furrow invagination in the drosophila melanogaster embryo. Journal of the mechanical behavior of biomedical materials, 1(2):188–198, 2008.

[22] Natalie C Heer, Pearson W Miller, Soline Chanet, Norbert Stoop, Jörn Dunkel, and Adam C Martin. Actomyosin-based tissue folding requires a multicellular myosin gradient. Development, 144(10):1876–1886, 2017.

[23] Hannah G Yevick, Pearson W Miller, Jörn Dunkel, and Adam C Martin. Structural redundancy in supracellular actomyosin networks enables robust tissue folding. Developmental cell, 50(5):586–598, 2019.

[24] Hélène Berthoumieux, Jean-Léon Maître, Carl-Philipp Heisenberg, Ewa K Paluch, Frank Jülicher, and Guillaume Salbreux. Active elastic thin shell theory for cellular deformations. New Journal of Physics, 16(6):065005, 2014.

[25] Hudson Borja da Rocha, Jérémy Bleyer, and Hervé Turlier. A viscous active shell theory of the cell cortex. arXiv e-prints, pages arXiv–2110, 2021.

[26] Bing He, Konstantin Doubrovinski, Oleg Polyakov, and Eric Wieschaus. Apical constriction drives tissue-scale hydrodynamic flow to mediate cell elongation. Nature, 508(7496):392–396, 2014.

[27] Gabor Forgacs, Ramsey A Foty, Yinon Shafrir, and Malcolm S Steinberg. Viscoelastic properties of living embryonic tissues: a quantitative study. Biophysical journal, 74(5):2227–2234, 1998.

[28] Konstantin Doubrovinski, Michael Swan, Oleg Polyakov, and Eric F Wieschaus. Measurement of cortical elasticity in drosophila melanogaster embryos using ferrofluids. Proceedings of the National Academy of Sciences, 114(5):1051–1056, 2017.

[29] Alok D Wessel, Maheshwar Gumalla, Jörg Grosshans, and Christoph F Schmidt. The mechanical properties of early drosophila embryos measured by high-speed video microrheology. Biophysical journal, 108(8):1899–1907, 2015.

[30] Hassan Masoud and Michael J Shelley. Collective surfing of chemically active particles. Physical review letters, 112(12):128304, 2014.

[31] Claire Bertet, Lawrence Sulak, and Thomas Lecuit. Myosin-dependent junction remodelling controls planar cell intercalation and axis elongation. Nature, 429(6992):667–671, 2004.

[32] Amanda Nicole Goldner and Konstantin Doubrovinski. What basal membranes can tell us about viscous forces in drosophila ventral furrow formation. bioRxiv, 2021.

[33] Robert J Huebner and John B Wallingford. Coming to consensus: a unifying model emerges for convergent extension. Developmental cell, 46(4):389–396, 2018.

[34] John Shih and Ray Keller. Cell motility driving mediolateral intercalation in explants of xenopus laevis. Development, 116(4):901–914, 1992.

[35] Margot Williams, Weiwei Yen, Xiaowei Lu, and Ann Sutherland. Distinct apical and basolateral mechanisms drive planar cell polarity-dependent convergent extension of the mouse neural plate. Developmental cell, 29(1):34–46, 2014.

[36] Mélanie Gracia, Sophie Theis, Amsha Proag, Guillaume Gay, Corinne Benassayag, and Magali Suzanne. Mechanical impact of epithelialmesenchymal transition on epithelial morphogenesis in drosophila. Nature communications, 10(1):1–17, 2019.

[37] Bruno Monier, Melanie Gettings, Guillaume Gay, Thomas Mangeat, Sonia Schott, Ana Guarner, and Magali Suzanne. Apico-basal forces exerted by apoptotic cells drive epithelium folding. Nature, 518(7538):245–248, 2015.

[38] Alphy John and Matteo Rauzi. A two-tier junctional mechanism drives simultaneous tissue folding and extension. Developmental Cell, 56(10):1469–1483, 2021.

[39] Ginger L Hunter, Janice M Crawford, Julian Z Genkins, and Daniel P Kiehart. Ion channels contribute to the regulation of cell sheet forces during drosophila dorsal closure. Development, 141(2):325–334, 2014.

[40] Neophytos Christodoulou and Paris A Skourides. Cell-autonomous ca2+ flashes elicit pulsed contractions of an apical actin network to drive apical constriction during neural tube closure. Cell reports, 13(10):2189–2202, 2015.

