## Supplemental Documents for "Modeling epithelial tissue and cell deformation dynamics using a viscoelastic slab sculpted by surface forces"

Note: equations and figures in the main text are referred to using the letter M preceding the equation or figure number, e.g Eq. (M4) and Fig. M4.

### I. NON-DIMENSIONALIZATION OF EQUATIONS

The equations of motion in the slab (region $S$), corresponding to the dimensionless forms Eqns. (M10) and (M11), have dimensional forms given by:

$$-\nabla_{\mathbf{x}} P + \eta \nabla_{\mathbf{x}}^2 \mathbf{u} + \nabla_{\mathbf{x}} \cdot \boldsymbol{\sigma}^e = 0 \tag{1}$$

$$\overset{\triangledown}{\boldsymbol{\sigma}^e} = D \nabla_{\mathbf{x}}^2 \boldsymbol{\sigma}^e - \frac{1}{\tau_p}(\boldsymbol{\sigma}^e - \kappa G \phi \mathbb{1}) \tag{2}$$

where we have defined $G_0 \equiv \kappa G \phi$ in Eq. (M5), $D$ is the center-of-mass diffusion constant for the Oldroyd-B particles, and $\eta$ is the solvent viscosity. The other parameters in Eq. (2) are given by:

$$\tau_p = \frac{6\pi\eta a}{8k_B T\beta^2} \sim t \quad , \quad G = \frac{N_b b^2}{3} \equiv \frac{1}{2\beta^2} \sim \ell^2 \quad , \quad \kappa = 2k_B T\beta^2 \sim \frac{\text{force}}{\ell} \quad , \tag{3}$$

where we have indicated the units of each of these constants. These parameters and Eq. (2) come from a kinetic theory for a suspension of Oldroyd-B particles [1]. The microscopic parameters appearing in $\beta$, $G$, and $\kappa$ are:

| | | |
|---|---|---|
| $N_b$ | Number of links in the polymer chain of an Oldroyd-B particle | (4) |
| $b$ | Length of each link in polymer chain of an Oldroyd-B particle | (5) |
| $a$ | Radius of ball on the dumbbell shape of an Oldroyd-B particle | (6) |
| $T$ | Temperature of the system | (7) |

The parameter $\tau_p$ is interpreted as a time scale of relaxation for the Oldroyd-B polymers. The parameter $G$ measures the magnitude of the isotropic steady state stress $\boldsymbol{\sigma}^e$ normalized to particle density and divided by the spring constant when velocities are $\mathbf{0}$, i.e. $\boldsymbol{\sigma}^e = \kappa G \phi \mathbb{1}$ if $\frac{\partial}{\partial t}\boldsymbol{\sigma}^e = \mathbf{0}$ and $\mathbf{u} = \mathbf{0}$. The parameter $\kappa$ measures the effective spring constant of an Oldroyd-B polymer assuming an entropic spring force model. We keep the $\mathbf{x}$ subscript on $\nabla_{\mathbf{x}}$ in Eqns. (1) and (2) to remind us that derivatives are dimensional, in contrast to $\nabla_{\mathbf{x}'}$ which will later indicate dimensionless derivatives. Here, we defined $\phi(\mathbf{x})$ as the number of Oldroyd-B particles per volume at location $\mathbf{x}$, or the particle density at $\mathbf{x}$:

$$\phi(\mathbf{x}, t) \equiv \int d^3\mathbf{R}\, \Psi(\mathbf{x}, \mathbf{R}, t) \sim \frac{1}{\ell^3} \tag{8}$$

where $\Psi(\mathbf{x}, \mathbf{R}, t)$ is the density of particles at position $\mathbf{x}$ with orientation $\mathbf{R}$.

To non-dimensionalize, we choose time and force (effectively mass) scales to depend on the intensive properties of the material. That is, the time scale is $\tau_p$, the polymer relaxation time, and the force scale is $\eta\ell_0^2/\tau_p$, where $\eta$ is the viscosity of the solvent. The length scale, $\ell_0$, is a yet-unspecified characteristic length scale and will be determined later based on the geometry of the problem.

$$\tau_p = \text{unit of time} \qquad , \qquad \eta\frac{\ell_0^2}{\tau_p} = \text{unit of force} \qquad , \qquad \ell_0 = \text{unit of length} \quad . \tag{9}$$

The units of stress are then:

$$\text{stress} \sim \frac{\text{force}}{\ell^2} \sim \frac{\eta}{\tau_p} \quad . \tag{10}$$

To rescale Eq. (2), we replace dimensional parameters with dimensionless numbers multiplied by the correct dimensional unit. We use primes to indicate dimensionless numbers:

$$\kappa = \kappa'\eta\frac{\ell_0}{\tau_p} \qquad , \qquad D = D'\frac{\ell_0^2}{\tau_p} \qquad , \qquad G = G'\ell_0^2 \qquad , \qquad \phi = \phi'\frac{1}{\ell_0^3} \quad . \tag{11}$$

The dimensionless derivatives are:

$$\triangledown = \frac{1}{\tau_p}\triangledown' \qquad , \qquad \nabla_{\mathbf{x}}^2 = \frac{1}{\ell_0^2}\nabla_{\mathbf{x}'}^2 \quad . \tag{12}$$

In anticipation of algebraic simplification, we rescale $\boldsymbol{\sigma}^e$ with additional dimensionless factors $\kappa'$, $G'$, and $\phi'$:

$$\boldsymbol{\sigma}^e = \boldsymbol{\sigma}^{e\prime}\,\kappa'G'\phi'\,\frac{\eta}{\tau_p} \equiv \boldsymbol{\sigma}^{e\prime}\alpha'\,\frac{\eta}{\tau_p} \qquad , \qquad \alpha' \equiv \kappa'G'\phi' \tag{13}$$

Note that $\alpha'\eta/\tau_p = \kappa G\phi \equiv G_0$ where $G_0$ is the notation that appears in the main text. Substituting these into Eq. (2), we have:

$$\left(\alpha'\frac{\eta}{\tau_p}\right)\frac{1}{\tau_p}\overset{\triangledown'}{\boldsymbol{\sigma}^{e\prime}} - \left(D'\frac{\ell_0^2}{\tau_p}\right)\frac{1}{\ell_0^2}\left(\alpha'\frac{\eta}{\tau_p}\right)\nabla_{\mathbf{x}'}^2\boldsymbol{\sigma}^{e\prime} + \frac{1}{\tau_p}\left(\alpha'\frac{\eta}{\tau_p}\boldsymbol{\sigma}^{e\prime} - \alpha'\frac{\eta}{\tau_p}\mathbb{1}\right) = 0 \tag{14}$$

where in the substitution, we used that $\kappa G\phi = \alpha'\eta/\tau_p$. The above then simplifies to the dimensionless form of Eq. (2):

$$\overset{\triangledown'}{\boldsymbol{\sigma}^{e\prime}} - D'\nabla_{\mathbf{x}'}^2\boldsymbol{\sigma}^{e\prime} + (\boldsymbol{\sigma}^{e\prime} - \mathbb{1}) = 0 \tag{15}$$

Notice from Eq. (13) that $\boldsymbol{\sigma}^e$ is proportional to particle density $\phi$, so $\boldsymbol{\sigma}^e$ contains a factor that accounts for the fact that if there are more Oldroyd-B particles in solution, then the extra stress is proportionally higher. In contrast, $\boldsymbol{\sigma}^{e\prime}$ does not include a factor proportional to the particle density, instead, the particle density is incorporated into the parameter $\alpha'$.

Let us now consider Eq. (1). We proceed as above and express all fields and derivatives as the multiplication of a non-dimensional number with a dimensional unit:

$$P = P'\frac{\eta}{\tau_p} \quad , \quad \mathbf{u} = \mathbf{u}'\frac{\ell_0}{\tau_p} \quad , \quad \boldsymbol{\sigma}^e = \boldsymbol{\sigma}^{e\prime}\alpha'\frac{\eta}{\tau_p} \quad , \quad \nabla_{\mathbf{x}} = \frac{1}{\ell_0}\nabla_{\mathbf{x}'} \quad , \quad \nabla^2_{\mathbf{x}} = \frac{1}{\ell_0^2}\nabla^2_{\mathbf{x}'} \quad . \tag{16}$$

Substituting these into Eq. (1), we have:

$$0 = -\nabla_{\mathbf{x}'}P' + \nabla^2_{\mathbf{x}'}\mathbf{u}' + \alpha'\nabla_{\mathbf{x}'}\cdot\boldsymbol{\sigma}^{e\prime} \tag{17}$$

where the non-dimensional parameter

$$\alpha' \equiv \kappa' G' \phi' \tag{18}$$

contains information about the microscopic model through $\kappa'$ and $G'$ and importantly is proportional to the (non-dimensional) number density $\phi'$ of Oldroyd-B particles in solution. The quantity $\alpha$ is a dimensionless parameter that effectively tunes the concentration of Oldroyd-B particles in solution or viscoelastic "strength" of the system.

Equations (M10) and (M11) are simply Eqns. (17) and (15) with the primes and the subscript $\mathbf{x}'$ on gradients removed.

#### A. Adding other fluids to the model

In our model description, in addition to the Stokes Oldroyd-B material in the slab (region $S$), we have a fluid layer (region $L$) and a fluid bath (region $B$), with viscosities $\eta_L$ and $\eta_B$, respectively. Consider the fluid layer $L$, where the Stokes equation holds for fields $P^+$ and $\mathbf{u}^+$ with viscosity $\eta_L$:

$$-\nabla_{\mathbf{x}}P^+ + \eta_L\nabla^2_{\mathbf{x}}\mathbf{u}^+ = 0 \quad . \tag{19}$$

When this equation is rescaled using $\eta$, $\ell_0$, and $\tau_p$, that is, when we substitute $P^+ = P^{+\prime}\eta/\tau_p$, $\eta_L = \eta_L'\eta$ and $\mathbf{u}^+ = \mathbf{u}^{+\prime}\ell_0/\tau_p$, use rescaled dimensionless derivatives, and eliminate common factors, we obtain the dimensionless equation:

$$-\nabla_{\mathbf{x}'}P^{+\prime} + \eta_L'\nabla^2_{\mathbf{x}'}\mathbf{u}^{+\prime} = 0 \tag{20}$$

We obtain a similar result if we consider the fluid bath $B$; there, the dimensionless form of the Stokes equation becomes:

$$-\nabla_{\mathbf{x}'} P^{-\prime} + \eta_B' \nabla_{\mathbf{x}'}^2 \mathbf{u}^{-\prime} = 0 \tag{21}$$

Equations (20) and (21) then become Eqns. (M9) and (M12) if all primes and the subscript $\mathbf{x}'$ on gradients are removed.

#### B. Driving forces

Given dimensionless Eqns. (17) and (15) in which all derivatives are dimensionless derivatives, then the Fourier Transform produces only dimensionless wavevectors, and driving forces will be naturally dimensionless since the stress jump between regions, for example:

$$[\boldsymbol{\sigma}'] = \mathbf{f}' \cos(\omega' t') s(\mathbf{x}') \tag{22}$$

produces only dimensionless values. When we vary the frequency and amplitude of driving in implementation, it is $\mathbf{f}'$ and $\omega'$ that we vary, and no additional non-dimensionalization needs to be considered. We can recover the dimensional forms of these dimensionless frequencies and amplitudes as:

$$\mathbf{f} = \mathbf{f}' \frac{\eta}{\tau_p} \quad , \quad \omega = \omega' \frac{1}{\tau_p} \quad . \tag{23}$$

Therefore, as long as we utilize dimensional estimates of $\eta$ and $\tau_p$ in S.I. units (via experiments or literature), we can convert our dimensionless force parameter values from numerical implementations to dimensional estimates.

#### C. Changing the intensive properties of the material

If we changed the material properties on which we rescale, for example $\tau_p \to \tau_p^{\mathrm{new}}$, then we would need to change the dimensionless constants. For example, dimensionless diffusion $D'$ is defined as:

$$D' = D \frac{\tau_p}{\ell_0^2} \tag{24}$$

where $D$ is a dimensional diffusion measurement, and $\ell_0$ is a length in the problem. To ensure that $\tau_p$ is altered while no other physical, dimensional parameters are altered, we need to recompute $D' = D\tau_p^{\text{new}}/\ell_0^2$ with the new $\tau_p$ while keeping $D$ and $\ell_0$ the same. This implies that if $\tau_p^{\text{new}} = 2\tau_p$ then $D'^{\text{new}} = 2D'$. The same is relevant for $\alpha'$; we had

$$\alpha' = \kappa G \phi \frac{\tau_p}{\eta} \quad . \tag{25}$$

So if $\tau_p^{\text{new}} = 2\tau_p$, then, since $\eta$, $\kappa$, $G$, $\phi$ do not change because as physical parameters, we also do not specify them to change, then we must have $\alpha'^{\text{new}} = 2\alpha'$. Non-dimensional parameters need to be changed whenever the intensive material parameters on which we rescale change.

### II. SOLUTION TO BULK EQUATIONS

#### A. Bulk velocity and pressure solutions given velocity boundary conditions

We solve the Fourier transformed Stokes equations, Eqns. (M21), (M22), and (M23) in the region $z \in [-h, 0]$. If velocities are $\tilde{\mathbf{V}}^1$ and $\tilde{\mathbf{V}}^2$ at $z = 0$ and $z = -h$, respectively, then the solution for $\tilde{w}$, the $z$-velocity in the bulk, is:

$$\begin{aligned} \tilde{w}(\mathbf{k}, z) =& \frac{i}{2}\left( (z+h)\frac{kh}{A}\left(e^{kz} - e^{-kz}\right) + (-z)\frac{\sinh(kh)}{A}\left(e^{k(z+h)} - e^{-k(z+h)}\right)\right)\mathbf{k}\cdot\tilde{\mathbf{V}}^1 \\ &+ \frac{i}{2}\left( (z+h)\frac{\sinh(kh)}{A}\left(e^{kz} - e^{-kz}\right) + (-z)\frac{kh}{A}\left(e^{k(z+h)} - e^{-k(z+h)}\right)\right)\mathbf{k}\cdot\tilde{\mathbf{V}}^2 \end{aligned} \tag{26}$$

$$A \equiv A(k, h) = \sinh(kh)^2 - (kh)^2 \tag{27}$$

where $A$, a parameter-free function of $(k, h)$, has been defined. In this region, the solution for pressure $\tilde{P}$ is:

$$\tilde{P}(\mathbf{k}, z) = \eta i\left(-\alpha_1 e^{k(z+h)} + \alpha_2 e^{-k(z+h)}\right)\mathbf{k}\cdot\tilde{\mathbf{V}}^1 + \eta i\left(\alpha_2 e^{kz} - \alpha_1 e^{-kz}\right)\mathbf{k}\cdot\tilde{\mathbf{V}}^2 \tag{28}$$

$$\begin{aligned} \alpha_1 \equiv \alpha_1(k, h) =& \left(\frac{\sinh(kh)}{A} - \frac{kh}{A}e^{-kh}\right) \\ \alpha_2 \equiv \alpha_2(k, h) =& \left(\frac{\sinh(kh)}{A} - \frac{kh}{A}e^{kh}\right) \end{aligned} \tag{29}$$

where $\alpha_1$ and $\alpha_2$, parameter-free functions of $(k, h)$, have been defined. Finally, in this region, the solution for $\tilde{\mathbf{v}}$, the $x, y$ velocity, is:

$$\tilde{\mathbf{v}}(\mathbf{k}, z) = \left(\tilde{\boldsymbol{\chi}}_A^1 e^{kz} + \tilde{\boldsymbol{\chi}}_B^1 e^{-kz}\right)\tilde{\mathbf{V}}^1 + \left(\tilde{\boldsymbol{\chi}}_A^2 e^{kz} + \tilde{\boldsymbol{\chi}}_B^2 e^{-kz}\right)\tilde{\mathbf{V}}^2 \tag{30}$$

where $\tilde{\boldsymbol{\chi}}$'s are matrices multiplying the $\tilde{\mathbf{V}}$ vectors given by:

$$\tilde{\boldsymbol{\chi}}_A^1 = \tilde{\boldsymbol{\chi}}_A^1(k,h) = \frac{1}{2\sinh(kh)}\left(\mathbb{1}e^{kh} - \hat{\mathbf{k}}\hat{\mathbf{k}}\frac{kh}{2}\alpha^0\right) + \frac{\alpha_1 e^{kh} kz}{2}\hat{\mathbf{k}}\hat{\mathbf{k}} \tag{31}$$

$$\tilde{\boldsymbol{\chi}}_B^1 = \tilde{\boldsymbol{\chi}}_B^1(k,h) = \frac{1}{2\sinh(kh)}\left(-\mathbb{1}e^{-kh} + \hat{\mathbf{k}}\hat{\mathbf{k}}\frac{kh}{2}\alpha^0\right) + \frac{\alpha_2 e^{-kh} kz}{2}\hat{\mathbf{k}}\hat{\mathbf{k}} \tag{32}$$

$$\tilde{\boldsymbol{\chi}}_A^2 = \tilde{\boldsymbol{\chi}}_A^2(k,h) = \frac{1}{2\sinh(kh)}\left(-\mathbb{1} + \hat{\mathbf{k}}\hat{\mathbf{k}}\frac{kh}{2}\alpha_C^0\right) - \frac{\alpha_2 kz}{2}\hat{\mathbf{k}}\hat{\mathbf{k}} \tag{33}$$

$$\tilde{\boldsymbol{\chi}}_B^2 = \tilde{\boldsymbol{\chi}}_B^2(k,h) = \frac{1}{2\sinh(kh)}\left(\mathbb{1} - \hat{\mathbf{k}}\hat{\mathbf{k}}\frac{kh}{2}\alpha_C^0\right) - \frac{\alpha_1 kz}{2}\hat{\mathbf{k}}\hat{\mathbf{k}} \tag{34}$$

and where $\alpha_C^0$ and $\alpha^0$ are parameter-free functions of $(k,h)$ defined by:

$$\begin{aligned} \alpha_C^0 \equiv& \alpha_C^0(k,h) \equiv \alpha_1 e^{kh} + \alpha_2 e^{-kh} = \frac{2}{A}(\sinh(kh)\cosh(kh) - kh) \\ \alpha^0 \equiv& \alpha^0(k,h) \equiv \alpha_1 + \alpha_2 = \frac{2}{A}(\sinh(kh) - kh\cosh(kh)) \quad . \end{aligned} \tag{35}$$

Here we have $(\hat{\mathbf{k}}\hat{\mathbf{k}})_{ij} = k_i k_j / k^2$.

#### *1. Velocity and pressure solutions in region S*

Region $S$ is exactly the region $z \in [-h, 0]$ for which solutions were presented in section II A. Hence, the bulk solutions for region $S$ are exactly as written in Eqns. (26), (28), and (30) only with the replacements $\tilde{\mathbf{V}}^1 \to \tilde{\mathbf{V}}^t$ and $\tilde{\mathbf{V}}^2 \to \tilde{\mathbf{V}}^b$.

#### *2. Velocity and pressure solutions in region L*

To obtain bulk pressure and velocity solutions for region $L$ (the fluid layer), we simply make replacements in Eqns. (26), (28), and (30) that $\eta \to \eta_L$ and $-h \to H$ and then set $\tilde{\mathbf{V}}^2 = \mathbf{0}$, as this will correspond to the velocity at the wall ($z = H$) after we have replaced $-h \to H$. It is helpful to notice as we change $h \to -H$, that from Eqns. (27), (29), and (35), we have:

$$A(k,-H) = A(k,H) \tag{36}$$

$$\alpha_1(k,-H) = -\alpha_2(k,H) \qquad , \qquad \alpha_2(k,-H) = -\alpha_1(k,H) \tag{37}$$

$$\alpha_C^0(k,-H) = -\alpha_C^0(k,H) \qquad , \qquad \alpha^0(k,-H) = -\alpha^0(k,H) \quad . \tag{38}$$

Explicitly, the solutions to the Stokes equations in region $L$ are as follows. For $\tilde{w}$ in region $L$:

$$\tilde{w}^{+}(\mathbf{k},z) = \frac{i}{2}\Big((z-H)\frac{-kH}{A(k,H)}\big(e^{kz}-e^{-kz}\big) + z\frac{\sinh(kH)}{A(k,H)}\big(e^{k(z-H)}-e^{-k(z-H)}\big)\Big)\mathbf{k}\cdot\tilde{\mathbf{V}}^{t} \quad . \tag{39}$$

For $\tilde{P}$ in region $L$:

$$\tilde{P}^{+}(\mathbf{k},z) = \eta_L i\big(\alpha_2(k,H)e^{k(z-H)} - \alpha_1(k,H)e^{-k(z-H)}\big)\mathbf{k}\cdot\tilde{\mathbf{V}}^{t} \quad . \tag{40}$$

For $\tilde{\mathbf{v}}$ in region $L$:

$$\tilde{\mathbf{v}}^{+}(\mathbf{k},z) = \big(\tilde{\boldsymbol{\chi}}^{1}_{A}(k,-H)e^{kz} + \tilde{\boldsymbol{\chi}}^{1}_{B}(k,-H)e^{-kz}\big)\tilde{\mathbf{V}}^{t} \quad , \tag{41}$$

where

$$\tilde{\boldsymbol{\chi}}^{1}_{A}(k,-H) = \frac{1}{2\sinh(kH)}\Big(-\mathbb{1}e^{-kH} + \hat{\mathbf{k}}\hat{\mathbf{k}}\frac{kH}{2}\alpha^{0}(k,H)\Big) - \frac{\alpha_2(k,H)e^{-kH}kz}{2}\hat{\mathbf{k}}\hat{\mathbf{k}} \tag{42}$$

$$\tilde{\boldsymbol{\chi}}^{1}_{B}(k,-H) = \frac{1}{2\sinh(kH)}\Big(\mathbb{1}e^{kH} - \hat{\mathbf{k}}\hat{\mathbf{k}}\frac{kH}{2}\alpha^{0}(k,H)\Big) - \frac{\alpha_1(k,H)e^{kH}kz}{2}\hat{\mathbf{k}}\hat{\mathbf{k}} \quad . \tag{43}$$

*3. Velocity and pressure solutions in region B*

To obtain bulk pressure and velocity solutions for region $B$ of the fluid bath, we make replacements and take limits in Eqns. (26), (28), and (30) as follows. First, we replace $\eta \to \eta_B$. Additionally, we take the limit $h \to \infty$ (or more intuitively $-h \to -\infty$) to represent that the bottom of the fluid bath is at negative infinity. We replace $\tilde{\mathbf{V}}^1 \to \tilde{\mathbf{V}}^b$ since the top surface of the fluid bath should match the velocity at the bottom of the slab $\tilde{\mathbf{V}}^b$. Furthermore, we set $\tilde{\mathbf{V}}^2 = \mathbf{0}$ as the velocity at the $z = -h \to -\infty$ side of the fluid bath must vanish. Finally, we replace the argument $z$ from Eqns. (26), (28), and (30) with $z+h$ due to the fact that the top of region $B$ at $z=-h$ should correspond to $z=0$ in the $z \in [-h, 0]$ geometry.

Explicitly, after making replacements and taking limits, the solutions to the Stokes equations in region $B$ are obtained from the solutions $z \in [-h, 0]$. For $\tilde{w}$ in region $B$:

$$\tilde{w}^{-}(\mathbf{k},z) = -ie^{k(z+h)}(z+h)\mathbf{k}\cdot\tilde{\mathbf{V}}^{b} \tag{44}$$

For $\tilde{P}$ in region $B$:

$$\tilde{P}^{-}(\mathbf{k},z) = -2\eta_B ie^{k(z+h)}\mathbf{k}\cdot\tilde{\mathbf{V}}^{b} \tag{45}$$

For $\tilde{\mathbf{v}}$ in region $B$:

$$\tilde{\mathbf{v}}^{-}(\mathbf{k},z) = \Big(\mathbb{1} + k(z+h)\hat{\mathbf{k}}\hat{\mathbf{k}}\Big)e^{k(z+h)}\tilde{\mathbf{V}}^{b} \tag{46}$$

#### B. Neumann-to-Dirichlet map of surface forces to surface velocities

Using the solutions for $\tilde{\mathbf{v}}$ and $\tilde{w}$ in Eqns. (30) and (26) and notating $\boldsymbol{\sigma} = -P\mathbb{1} + \eta\Big(\nabla\mathbf{u} + (\nabla\mathbf{u})^{T}\Big)$, shear stresses evaluated at the boundaries $z = 0$ and $z = -h$ are:

$$\mathbf{s}^{1} = \boldsymbol{\sigma}\hat{\mathbf{z}}\big|_{z=0,\text{2D}} = \eta\frac{\partial\tilde{\mathbf{v}}}{\partial z}\bigg|_{z=0} + \eta\begin{pmatrix}\frac{\partial\tilde{w}}{\partial x}\\ \frac{\partial\tilde{w}}{\partial y}\end{pmatrix}\Bigg|_{z=0} = \boldsymbol{\Theta}_1\tilde{\mathbf{V}}^{1} - \boldsymbol{\Theta}_2\tilde{\mathbf{V}}^{2} \tag{47}$$

$$\mathbf{s}^{2} = \boldsymbol{\sigma}\hat{\mathbf{z}}\big|_{z=-h,\text{2D}} = \eta\frac{\partial\hat{\mathbf{v}}}{\partial z}\bigg|_{z=-h} + \eta\begin{pmatrix}\frac{\partial\tilde{w}}{\partial x}\\ \frac{\partial\tilde{w}}{\partial y}\end{pmatrix}\Bigg|_{z=-h} = \boldsymbol{\Theta}_2\tilde{\mathbf{V}}^{1} - \boldsymbol{\Theta}_1\tilde{\mathbf{V}}^{2} \tag{48}$$

where

$$\begin{aligned}\boldsymbol{\Theta}_1 \equiv \boldsymbol{\Theta}_1(k,h,\eta) =& \eta k\left(\coth(kh)\mathbb{1} + \frac{\hat{\mathbf{k}}\hat{\mathbf{k}}}{2}\left(\frac{-kh\alpha^{0}}{\sinh(kh)} + \alpha_C^{0}\right)\right)\\ \boldsymbol{\Theta}_2 \equiv \boldsymbol{\Theta}_2(k,h,\eta) =& \eta k\left(\frac{1}{\sinh(kh)}\mathbb{1} + \frac{\hat{\mathbf{k}}\hat{\mathbf{k}}}{2}\left(\frac{-kh\alpha_C^{0}}{\sinh(kh)} + \alpha^{0}\right)\right) \quad .\end{aligned} \tag{49}$$

Here, we include the viscosity $\eta$ as an argument in $\boldsymbol{\Theta}_1$ and $\boldsymbol{\Theta}_2$ should we refer to a similar expression for $\boldsymbol{\Theta}_1$ or $\boldsymbol{\Theta}_2$ with a different viscosity. Equations (47) and (48) correspond to Eq. (M26) except with superscripts 1 and 2 labeling the velocities $\tilde{\mathbf{V}}^{1,2}$ and stresses $\mathbf{s}^{1,2}$. We keep these superscripts in this section because we wish to map the expressions in $z \in [-h, 0]$ to regions $L$ and $B$. Note that the derivatives involving $\tilde{w}$ in Eqns. (47) and (48) all vanish: $\frac{\partial}{\partial x}\tilde{w}\big|_{z=0} = \frac{\partial}{\partial y}\tilde{w}\big|_{z=0} = \frac{\partial}{\partial x}\tilde{w}\big|_{z=-h} = \frac{\partial}{\partial y}\tilde{w}\big|_{z=-h} = 0$.

##### *1. Neumann-to-Dirichlet map in region S*

Since we recognize that region $S$ is exactly $z \in [-h, 0]$, then all for all expressions, we need only to replace $\tilde{\mathbf{V}}^{1} \to \tilde{\mathbf{V}}^{t}$ and $\tilde{\mathbf{V}}^{2} \to \tilde{\mathbf{V}}^{b}$. Therefore, for the shear stress at the boundaries of $S$, from Eqns. (47), (48), and (49), we have:

$$\begin{aligned}\mathbf{s}_S^{t} =& \boldsymbol{\Theta}_1(k,h,\eta)\tilde{\mathbf{V}}^{t} - \boldsymbol{\Theta}_2(k,h,\eta)\tilde{\mathbf{V}}^{b}\\ \mathbf{s}_S^{b} =& \boldsymbol{\Theta}_2(k,h,\eta)\tilde{\mathbf{V}}^{t} - \boldsymbol{\Theta}_1(k,h,\eta)\tilde{\mathbf{V}}^{b}\end{aligned} \tag{50}$$

as stated by Eq. (M26).

*2. Neumann-to-Dirichlet map in region L*

As noted, to obtain expressions for region $L$, we simply needed to utilize expressions in $z \in [-h, 0]$ and make replacements $\eta \to \eta_L$ and $-h \to H$, and then set $\tilde{\mathbf{V}}^2 = \mathbf{0}$. These replacements applied to Eqns. (47) and (49) give the stress at the $z = 0$ boundary of the fluid layer $L$ as:

$$\mathbf{s}_L \equiv \boldsymbol{\sigma}^L \hat{\mathbf{z}}\Big|_{z=0,\text{2D}} = -\boldsymbol{\Theta}_1(k, H, \eta_L)\tilde{\mathbf{V}}^t \equiv -\boldsymbol{\Lambda}(k, H, \eta_L)\tilde{\mathbf{V}}^t \tag{51}$$

where we use $\boldsymbol{\Lambda}(k, H, \eta_L) \equiv \boldsymbol{\Theta}_1(k, H, \eta_L)$ to avoid confusion with $\boldsymbol{\Theta}_1(k, h, \eta)$. This corresponds to Eq. (M27).

*3. Neumann-to-Dirichlet map in region B*

We noted that to obtain expressions in region $B$, we utilize expressions in $z \in [-h, 0]$, replace $\eta \to \eta_B$, take the limit $h \to \infty$, replace $\tilde{\mathbf{V}}^1 \to \tilde{\mathbf{V}}^b$, set $\tilde{\mathbf{V}}^2 = \mathbf{0}$, and finally replace the argument $z$ with $z + h$. Therefore, for the shear stress at the top boundary of region $B$, from Eq. (47) and (49), we have that:

$$\mathbf{s}_B \equiv \boldsymbol{\sigma}^B \hat{\mathbf{z}}\Big|_{z=-h,\text{2D}} = \boldsymbol{\Theta}_1(k, h \to \infty, \eta_B)\tilde{\mathbf{V}}^b = \eta_B k\left(\mathbb{1} + \hat{\mathbf{k}}\hat{\mathbf{k}}\right)\tilde{\mathbf{V}}^b \equiv \boldsymbol{\beta}(k, \eta_B)\tilde{\mathbf{V}}^b \tag{52}$$

This corresponds to Eq. (M28). The quantity $\mathbf{s}_B$ can also be calculated using direct derivatives of Eqns. (46) and (44), i.e

$$\mathbf{s}_B = \eta_B \left( \frac{\partial \hat{\mathbf{v}}^-}{\partial z}\bigg|_{z=-h} + \begin{pmatrix} \frac{\partial \tilde{w}^-}{\partial x} \\ \frac{\partial \tilde{w}^-}{\partial y} \end{pmatrix}\bigg|_{z=-h} \right) \tag{53}$$

which will provide the same expression as Eq. (52).

*4. Relating active force to stress jump at the boundaries*

Listing all surface shear stresses, we have:

$$\begin{aligned}
\mathbf{s}_L &= -\boldsymbol{\Theta}_1(k, H, \eta_L)\tilde{\mathbf{V}}^t \equiv -\boldsymbol{\Lambda}(k, H, \eta_L)\tilde{\mathbf{V}}^t \\
\mathbf{s}_S^t &= \boldsymbol{\Theta}_1(k, h, \eta)\tilde{\mathbf{V}}^t - \boldsymbol{\Theta}_2(k, h, \eta)\tilde{\mathbf{V}}^b \\
\mathbf{s}_S^b &= \boldsymbol{\Theta}_2(k, h, \eta)\tilde{\mathbf{V}}^t - \boldsymbol{\Theta}_1(k, h, \eta)\tilde{\mathbf{V}}^b \\
\mathbf{s}_B &= \eta_B k\left(\mathbb{1} + \hat{\mathbf{k}}\hat{\mathbf{k}}\right)\tilde{\mathbf{V}}^b \equiv \boldsymbol{\beta}(k, \eta_B)\tilde{\mathbf{V}}^b
\end{aligned} \tag{54}$$

We assume that the shear stress jumps at $z = 0$ and $z = -h$ correspond to active forces $\mathbf{F}^t$ at $z = 0$ and $\mathbf{F}^b$ at $z = -h$. Then setting the shear stress jumps to active forces on the 2D surfaces gives:

$$\mathbf{s}_S^t - \mathbf{s}_L = \tilde{\mathbf{F}}^t \qquad \Longrightarrow \qquad \boldsymbol{\Theta}_1 \tilde{\mathbf{V}}^t - \boldsymbol{\Theta}_2 \tilde{\mathbf{V}}^b + \boldsymbol{\Lambda} \tilde{\mathbf{V}}^t = \tilde{\mathbf{F}}^t \tag{55}$$

$$\mathbf{s}_B - \mathbf{s}_S^b = \tilde{\mathbf{F}}^b \qquad \Longrightarrow \qquad \boldsymbol{\beta} \tilde{\mathbf{V}}^b - \boldsymbol{\Theta}_2 \tilde{\mathbf{V}}^t + \boldsymbol{\Theta}_1 \tilde{\mathbf{V}}^b = \tilde{\mathbf{F}}^b \quad . \tag{56}$$

Defining:

$$\boldsymbol{\Gamma}_B \equiv \boldsymbol{\Theta}_1 + \boldsymbol{\beta} \qquad , \qquad \boldsymbol{\Gamma}_L \equiv \boldsymbol{\Theta}_1 + \boldsymbol{\Lambda} \tag{57}$$

and inverting Eqns. (55) and (56), we obtain $\tilde{\mathbf{V}}^t$ and $\tilde{\mathbf{V}}^b$ in terms of $\tilde{\mathbf{F}}^t$ and $\tilde{\mathbf{F}}^b$:

$$\tilde{\mathbf{V}}^t = -\left(\boldsymbol{\Theta}_2 - \boldsymbol{\Gamma}_B \boldsymbol{\Theta}_2^{-1} \boldsymbol{\Gamma}_L\right)^{-1} \left(\tilde{\mathbf{F}}^b + \boldsymbol{\Gamma}_B \boldsymbol{\Theta}_2^{-1} \tilde{\mathbf{F}}^t\right) \tag{58}$$

$$\tilde{\mathbf{V}}^b = -\left(\boldsymbol{\Theta}_2 - \boldsymbol{\Gamma}_L \boldsymbol{\Theta}_2^{-1} \boldsymbol{\Gamma}_B\right)^{-1} \left(\tilde{\mathbf{F}}^t + \boldsymbol{\Gamma}_L \boldsymbol{\Theta}_2^{-1} \tilde{\mathbf{F}}^b\right) \tag{59}$$

which are the Neumann-to-Dirichlet maps (Eqns. (M31) and (M32)).

#### C. Calculation of the $z$-directional force densities at the boundaries

The force density in the $z$ direction on the $\hat{\mathbf{z}}$-directed surface in the pure Stokes solution is given by:

$$\sigma_{33} = 2\eta \frac{\partial}{\partial z} \tilde{w} - \tilde{P} \tag{60}$$

where $\eta$ is the viscosity of the Stokes fluid.

Since, in region $S$, we have that the solution to the Stokes equations are exactly Eqns. (26), (28), and (30) with $\tilde{\mathbf{V}}^1 \to \tilde{\mathbf{V}}^t$ and $\tilde{\mathbf{V}}^2 \to \tilde{\mathbf{V}}^b$, then using Eqns. (26) and (28), we have at $z = 0$:

$$\sigma_{33,S}^t = 2\eta \frac{\partial}{\partial z} \tilde{w} - \tilde{P} \bigg|_{z=0} = i2\eta \left(\frac{(kh)^2}{A(k,h)}\right) \mathbf{k} \cdot \tilde{\mathbf{V}}^t + i2\eta kh \left(\frac{\sinh(kh)}{A(k,h)}\right) \mathbf{k} \cdot \tilde{\mathbf{V}}^b \quad . \tag{61}$$

At the $z = -h$ boundary, we have:

$$\sigma_{33,S}^b = 2\eta \frac{\partial}{\partial z} \tilde{w} - \tilde{P} \bigg|_{z=-h} = i2\eta kh \left(\frac{\sinh(kh)}{A(k,h)}\right) \mathbf{k} \cdot \tilde{\mathbf{V}}^t + i2\eta \left(\frac{(kh)^2}{A(k,h)}\right) \mathbf{k} \cdot \tilde{\mathbf{V}}^b \tag{62}$$

where $A(k,h)$ is defined in Eq. (27).

For region $L$, we are interested in the force on the slab at $z = 0$, hence we calculate $\sigma_{33,L}\big|_{z=0}$. We obtain this from Eqns. (39) and (40) (or alternatively by taking $\sigma^t_{33,S}$ and replacing $h \to -H$, $\eta \to \eta_L$ and $\tilde{\mathbf{V}}^b \to 0$):

$$\sigma_{33,L} = 2\eta_L \frac{\partial}{\partial z}\tilde{w}^+ - \tilde{P}^+\Bigg|_{z=0} = i2\eta_L\left(\frac{(kH)^2}{A(k,H)}\right)\mathbf{k}\cdot\tilde{\mathbf{V}}^t \quad . \tag{63}$$

For region $B$, we are interested in the force on the slab at $z = -h$, hence we calculate $\sigma_{33,B}\big|_{z=-h}$. We obtain this from Eqns. (44) and (45):

$$\sigma_{33,B} = 2\eta_B \frac{\partial \tilde{w}^-}{\partial z} - \tilde{P}^-\Bigg|_{z=-h} = i2\eta_B(-k(z+h))\mathbf{k}\cdot\tilde{\mathbf{V}}^b\big|_{z=-h} = 0 \tag{64}$$

Interestingly, this is 0.

To compute the constraint force in the $z$ direction required to hold the surfaces flat in the Stokes model (with $\alpha = 0$), we demand that:

$$\begin{aligned}\sigma^t_{33,S} - \sigma_{33,L} =& \tilde{F}^{t,C}_z \\ \sigma_{33,B} - \sigma^b_{33,S} =& \tilde{F}^{b,C}_z\end{aligned} \tag{65}$$

where $\tilde{F}^{t,C}_z$ and $\tilde{F}^{b,C}_z$ are the constraint forces at the top and bottom of the slab, respectively. Using Eqns. (61), (62), (63), and (64), we obtain:

$$\begin{aligned}\tilde{F}^{t,C}_z =& i2\left(\eta\frac{(kh)^2}{A(k,h)} - \eta_L\frac{(kH)^2}{A(k,H)}\right)\mathbf{k}\cdot\tilde{\mathbf{V}}^t + i2\eta\frac{kh\sinh(kh)}{A(k,h)}\mathbf{k}\cdot\tilde{\mathbf{V}}^b \\ \tilde{F}^{b,C}_z =& -\sigma^b_{33,S} = -i2\eta\left(\frac{kh\sinh(kh)}{A(k,h)}\right)\mathbf{k}\cdot\tilde{\mathbf{V}}^t - i2\eta\left(\frac{(kh)^2}{A(k,h)}\right)\mathbf{k}\cdot\tilde{\mathbf{V}}^b \quad .\end{aligned} \tag{66}$$

If the Oldroyd-B extra stress were included in the model ($\alpha \neq 0$), then force densities $\sigma^t_{33,S}$ and $\sigma^b_{33,S}$ would be replaced by $\sigma^t_{33,S,\mathrm{OB}}$ and $\sigma^b_{33,S,\mathrm{OB}}$, respectively. Expressions $\sigma^t_{33,S,\mathrm{OB}}$ and $\sigma^b_{33,S,\mathrm{OB}}$ would include the contribution from the numerical particular solution $w_p$ and $P_p$:

$$\sigma_{33,S,\mathrm{OB}} = 2\eta\left(\frac{\partial\tilde{w}}{\partial z} + \frac{\partial\tilde{w}_p}{\partial z}\right) - (\tilde{P} + \tilde{P}_p) \quad . \tag{67}$$

Hence, the $z$-directional constraint force densities will be:

$$\tilde{F}^{t,C}_{z,\mathrm{OB}} = i2\left(\eta\frac{(kh)^2}{A(k,h)} - \eta_L\frac{(kH)^2}{A(k,H)}\right)\mathbf{k}\cdot\tilde{\mathbf{V}}^t + i2\eta\frac{kh\sinh(kh)}{A(k,h)}\mathbf{k}\cdot\tilde{\mathbf{V}}^b + \left(2\eta\frac{\partial\tilde{w}_p}{\partial z} - \tilde{P}_p\right)\Bigg|_{z=0} \tag{68}$$

$$\tilde{F}^{b,C}_{z,\mathrm{OB}} = -i2\eta\left(\frac{kh\sinh(kh)}{A(k,h)}\right)\mathbf{k}\cdot\tilde{\mathbf{V}}^t - i2\eta\left(\frac{(kh)^2}{A(k,h)}\right)\mathbf{k}\cdot\tilde{\mathbf{V}}^b - \left(2\eta\frac{\partial\tilde{w}_p}{\partial z} - \tilde{P}_p\right)\Bigg|_{z=-h} \tag{69}$$

For our implementation to find $F^{t,C}_{z,\mathrm{OB}}$ and $F^{b,C}_{z,\mathrm{OB}}$, we compute in $\mathbf{k}$ space the parts of the above not containing $w_p$ and $P_p$, we inverse Fourier transform into $(x, y)$ coordinates, and then add the contributions from numerical solutions $w_p(x, y)$ and $P_p(x, y)$. The quantities $F_z$(top) and $F_z$(bottom) discussed in Fig. M8 correspond to $-F^{t,C}_{z,\mathrm{OB}}$ and $-F^{b,C}_{z,\mathrm{OB}}$, respectively; they indicate the force on the tissue, applied against the constraint.

### III. CONSTRUCTING THE SOLUTION TO STOKES-OLDROYDB

If $\alpha \neq 0$, then the full forced Stokes-OldroydB equation must be solved in region $S$. This necessarily involves obtaining both a homogeneous solution (section (II A)) and a particular solution for Eq. (M10). Additionally, we must impose boundary conditions on the full solution. We claim that to satisfy all necessary boundary conditions on the full solution, we can choose the particular solution in region $S$ to satisfy *zero* velocity boundary conditions at the boundaries $z = 0$ and $z = -h$; we claim that this choice would leave us with a problem that is homogeneous and Newtonian everywhere, with familiar velocity and stress boundary conditions that we can satisfy using the analytical methods of section (II A). In the section below, we formally state and demonstrate this claim.

Consider regions $L$ and $S$. We seek solutions $P^+$, $\mathbf{u}+$, $P$, and $\mathbf{u}$ that satisfy Eqns. (M9) and (M10). Let the viscosity $\eta$ in region $S$ (rescaled to 1 using the non-dimensionalization in Eq. (9)) be reinstated here for clarity. The stress tensors in regions $L$ and $S$ are:

$$L: \quad \boldsymbol{\sigma}^L = \boldsymbol{\sigma}_N(P^+, \mathbf{u}^+, \eta_L) \tag{70}$$

$$S: \quad \boldsymbol{\sigma}^S = \boldsymbol{\sigma}_N(P, \mathbf{u}, \eta) + \alpha\boldsymbol{\sigma}^e \tag{71}$$

where the Newtonian stress has been notated as a function of pressure, velocity, and viscosity: $\boldsymbol{\sigma}_N(P, \mathbf{u}, \eta) \equiv -P\mathbb{1} + \eta(\nabla\mathbf{u} + \nabla\mathbf{u}^T)$.

In $L$, we have that Eq. (M9) is already homogenous and therefore $P^+$ and $\mathbf{u}^+$ solve:

$$-\nabla P^+ + \eta_L \nabla^2 \mathbf{u}^+ = 0 \tag{72}$$

in $L$ with $P^+$ and $\mathbf{u}^+$ as stated in section (II A 2); however boundary conditions have yet to be imposed.

In $S$, we construct both homogeneous (subscript $h$) and particular (subscript $p$) solutions. Let $P_h$ and $\mathbf{u}_h$ be the homogenous solution that solves:

$$-\nabla P_h + \eta \nabla^2 \mathbf{u}_h = 0 \tag{73}$$

and let $P_p$ and $\mathbf{u}_p$ be the particular solution that solves:

$$-\nabla P_p + \eta \nabla^2 \mathbf{u}_p + \alpha \nabla \cdot \boldsymbol{\sigma}^e = 0 \quad . \tag{74}$$

From Eqns. (73) and (74), the sums $P = P_h + P_p$ and $\mathbf{u} = \mathbf{u}_h + \mathbf{u}_p$ solve Eq. (M10) in $S$:

$$-\nabla P + \eta \nabla^2 \mathbf{u} + \alpha \nabla \cdot \boldsymbol{\sigma}^e = 0 \quad . \tag{75}$$

Note also that:

$$\boldsymbol{\sigma}_N(P, \mathbf{u}, \eta) = \boldsymbol{\sigma}_N(P_h, \mathbf{u}_h, \eta) + \boldsymbol{\sigma}_N(P_p, \mathbf{u}_p, \eta) \quad . \tag{76}$$

We construct particular solutions for $P_p$ and $\mathbf{u}_p$ numerically by solving Eq. (74) with *zero* velocity boundary conditions at the boundary between $L$ and $S$ at $z = 0$. That is, we obtain $P_p$ and $\mathbf{u}_p$ as numerical solutions satisfying:

$$-\nabla P_p + \eta \nabla^2 \mathbf{u}_p + \alpha \nabla \cdot \boldsymbol{\sigma}^e = 0 \quad \text{with} \quad \mathbf{u}_p\big|_{z=0} = \mathbf{0} \quad . \tag{77}$$

Claim: If the numerical solutions $P_p$ and $\mathbf{u}_p$ satisfy the boundary conditions in Eq. (77), then, to obtain the remaining fields $P^+$, $\mathbf{u}^+$, $P_h$, and $\mathbf{u}_h$ with all boundary conditions imposed (Eqns. (M14) and (M17)), we need only to solve a problem that is both homogeneous and Newtonian in regions $L$ and $S$ and that contains a familiar velocity boundary condition with a well-defined stress jump. This is the familiar problem for which we obtained the solution in section (II A).

To define the familiar problem, we impose the required boundary conditions for $P^+$, $\mathbf{u}^+$, $P$, and $\mathbf{u}$, the full solutions in $L$ and $S$. First, the velocity condition Eq. (M14) requires:

$$\mathbf{u}^+\big|_{z=0,2\mathrm{D}} = \mathbf{u}\big|_{z=0,2\mathrm{D}} = \mathbf{u}_h\big|_{z=0,2\mathrm{D}} + \mathbf{u}_p\big|_{z=0,2\mathrm{D}} = \mathbf{u}_h\big|_{z=0,2\mathrm{D}} + \mathbf{0} \tag{78}$$

$$\Longrightarrow \mathbf{u}^+\big|_{z=0,2\mathrm{D}} = \mathbf{u}_h\big|_{z=0,2\mathrm{D}} \equiv \mathbf{V}^t \tag{79}$$

since $\mathbf{u}_p\big|_{z=0,2\mathrm{D}} = \mathbf{0}$ by Eq. (77). Moreover, we have:

$$0 = w^+\big|_{z=0,2\mathrm{D}} = w\big|_{z=0,2\mathrm{D}} = w_h\big|_{z=0,2\mathrm{D}} + w_p\big|_{z=0,2\mathrm{D}} = w_h\big|_{z=0,2\mathrm{D}} + 0 \tag{80}$$

$$\Longrightarrow w^+\big|_{z=0,2\mathrm{D}} = w_h\big|_{z=0,2\mathrm{D}} = 0 \tag{81}$$

The conditions in Eqns. (79) and (81) say that planar ($x$ and $y$) velocities are continuous and $z$ velocities are 0 for $\mathbf{u}^+$ and $\mathbf{u}_h$ at the $z = 0$ boundary; this is the same velocity condition as Eq. (M14).

We similarly apply the stress jump condition (Eq. (M17)) to the full solutions $P^+$, $\mathbf{u}^+$, $P$, and $\mathbf{u}$:

$$\begin{aligned}\mathbf{F}^t =&(\boldsymbol{\sigma}^S - \boldsymbol{\sigma}^L)\hat{\mathbf{z}}\big|_{z=0,2\text{D}} = \big(\boldsymbol{\sigma}_N(P, \mathbf{u}, \eta) + \alpha\boldsymbol{\sigma}^e - \boldsymbol{\sigma}_N(P^+, \mathbf{u}^+, \nu)\big) \cdot \hat{\mathbf{z}}\big|_{z=0,2\text{D}} \\ =&\big(\boldsymbol{\sigma}_N(P_h, \mathbf{u}_h, \eta) + \boldsymbol{\sigma}_N(P_p, \mathbf{u}_p, \eta) + \alpha\boldsymbol{\sigma}^e - \boldsymbol{\sigma}_N(P^+, \mathbf{u}^+, \nu)\big) \cdot \hat{\mathbf{z}}\big|_{z=0,2\text{D}}\end{aligned}$$

where we used Eq. (76). Rearranging, we have:

$$\begin{aligned}\big(\boldsymbol{\sigma}_N(P_h, \mathbf{u}_h, \eta) - \boldsymbol{\sigma}_N(P^+, \mathbf{u}^+, \eta)\big)\hat{\mathbf{z}}\big|_{z=0,2\text{D}} =&\mathbf{F}^t - (\boldsymbol{\sigma}_N(P_p, \mathbf{u}_p, \eta) + \alpha\boldsymbol{\sigma}^e) \cdot \hat{\mathbf{z}}\big|_{z=0,2\text{D}} \\ \equiv&\mathbf{F}^t_{\text{OB}} \qquad (82)\end{aligned}$$

where $\mathbf{F}^t_{\text{OB}}$ is defined as the active force $\mathbf{F}^t$ on the surface plus corrections due to the extra stress $\boldsymbol{\sigma}^e$ and due to the Newtonian stress induced by the particular solution $\boldsymbol{\sigma}_N(P_p, \mathbf{u}_p, \eta)$. These corrections are numerically obtained. So Eq. (82) becomes a well-defined stress boundary condition for solutions of $P^+$, $\mathbf{u}^+$, $P_h$, and $\mathbf{u}_h$; this boundary condition is analogous to Eq. (M17). Hence, Eqns. (79), (81), and (82) are familiar boundary conditions that must be imposed on the fields $P^+$, $\mathbf{u}^+$, $P_h$, and $\mathbf{u}_h$ which all satisfy homogenous, Newtonian equations in the bulk (Eqns. (72) and (73)). This is as promised in the claim.

A similar analysis will show that the proposed numerical solutions for $\mathbf{u}_p$ and $P_p$ in Eq. (77) are also sufficient to define familiar velocity and stress boundary conditions for $P^-$, $\mathbf{u}^-$, $P_h$, and $\mathbf{u}_h$ between regions $S$ and $B$ analogous to Eqns. (79), (81), and (82), provided that $z = 0$ is replaced by $z = -h$ and that the active force $\mathbf{F}^b$ is amended to $\mathbf{F}^b_{\text{OB}}$ as indicated below. The quantities:

$$\mathbf{F}^t_{\text{OB}} =\mathbf{F}^t - (\boldsymbol{\sigma}_N(P_p, \mathbf{u}_p, \eta) + \alpha\boldsymbol{\sigma}^e) \cdot \hat{\mathbf{z}}\big|_{z=0,2\text{D}} \qquad (83)$$

$$\mathbf{F}^b_{\text{OB}} =\mathbf{F}^b + (\boldsymbol{\sigma}_N(P_p, \mathbf{u}_p, \eta) + \alpha\boldsymbol{\sigma}^e) \cdot \hat{\mathbf{z}}\big|_{z=-h,2\text{D}} \qquad (84)$$

are then Fourier transformed and substituted for $\tilde{\mathbf{F}}^t$ and $\tilde{\mathbf{F}}^b$ in Eqns. (M31) and (M32) to find the velocities $\tilde{\mathbf{V}}^t$ and $\tilde{\mathbf{V}}^b$. Explicitly:

$$\tilde{\mathbf{V}}^t = -\left(\boldsymbol{\Theta}_2 - \boldsymbol{\Gamma}_B\boldsymbol{\Theta}_2^{-1}\boldsymbol{\Gamma}_L\right)^{-1}\left(\tilde{\mathbf{F}}^b_{\text{OB}} + \boldsymbol{\Gamma}_B\boldsymbol{\Theta}_2^{-1}\tilde{\mathbf{F}}^t_{\text{OB}}\right) \qquad (85)$$

$$\tilde{\mathbf{V}}^b = -\left(\boldsymbol{\Theta}_2 - \boldsymbol{\Gamma}_L\boldsymbol{\Theta}_2^{-1}\boldsymbol{\Gamma}_B\right)^{-1}\left(\tilde{\mathbf{F}}^t_{\text{OB}} + \boldsymbol{\Gamma}_L\boldsymbol{\Theta}_2^{-1}\tilde{\mathbf{F}}^b_{\text{OB}}\right) \qquad (86)$$

as stated in Eqns. (M35) and (M36).

### IV. THE TRANSFER MATRICES

#### A. Expressing the action of $\boldsymbol{\tau}$ on $\tilde{\mathbf{F}}$ in terms of $\tilde{\mathbf{F}}$ and $\tilde{\mathbf{F}}^{\perp}$

To derive Eq. (M38), we use that the eigenvectors and eigenvalues of $\boldsymbol{\tau} = a(k)\mathbb{1} + b(k)\,\hat{\mathbf{k}}\hat{\mathbf{k}}$ are:

$$e_1 = a(k) + b(k) \qquad , \qquad v_1 = \hat{\mathbf{k}} \tag{87}$$

$$e_2 = a(k) \qquad , \qquad v_2 = \hat{\mathbf{k}}^{\perp} \equiv (-k_y, k_x)/k \quad . \tag{88}$$

We therefore write the action of $\boldsymbol{\tau}$ on an arbitrary force $\tilde{\mathbf{F}}$ as:

$$\boldsymbol{\tau}\tilde{\mathbf{F}} = \boldsymbol{\tau}\Big((\tilde{\mathbf{F}}\cdot\hat{\mathbf{k}})\hat{\mathbf{k}} + (\tilde{\mathbf{F}}\cdot\hat{\mathbf{k}}^{\perp})\hat{\mathbf{k}}^{\perp}\Big) \tag{89}$$

$$= (\tilde{\mathbf{F}}\cdot\hat{\mathbf{k}})(\boldsymbol{\tau}\hat{\mathbf{k}}) + (\tilde{\mathbf{F}}\cdot\hat{\mathbf{k}}^{\perp})(\boldsymbol{\tau}\hat{\mathbf{k}}^{\perp}) \tag{90}$$

$$= (\tilde{\mathbf{F}}\cdot\hat{\mathbf{k}})(a+b)\hat{\mathbf{k}} + (\tilde{\mathbf{F}}\cdot\hat{\mathbf{k}}^{\perp})(a)\hat{\mathbf{k}}^{\perp} \tag{91}$$

where the first equality decomposed $\tilde{\mathbf{F}}$ into $\tilde{\mathbf{F}} = (\tilde{\mathbf{F}}\cdot\hat{\mathbf{k}})\hat{\mathbf{k}} + (\tilde{\mathbf{F}}\cdot\hat{\mathbf{k}}^{\perp})\hat{\mathbf{k}}^{\perp}$ and the third equality used the fact that $\hat{\mathbf{k}}$ and $\hat{\mathbf{k}}^{\perp}$ are eigenvectors of $\boldsymbol{\tau}$ with eigenvalues $a+b$ and $a$, respectively.

In order to see how $\boldsymbol{\tau}$ acts in directions parallel to $\tilde{\mathbf{F}}$ and perpendicular to $\tilde{\mathbf{F}}$, we rewrite Eq. (91) with $\tilde{\mathbf{F}}$ and $\tilde{\mathbf{F}}^{\perp}$ as a basis instead of $\mathbf{k}$ and $\mathbf{k}^{\perp}$. This is done by decomposing $\hat{\mathbf{k}}$ and $\hat{\mathbf{k}}^{\perp}$ into components parallel and perpendicular to $\tilde{\mathbf{F}}$:

$$\hat{\mathbf{k}} = (\hat{\mathbf{k}}\cdot\hat{\tilde{\mathbf{F}}})\hat{\tilde{\mathbf{F}}} + (\hat{\mathbf{k}}\cdot\hat{\tilde{\mathbf{F}}}^{\perp})\hat{\tilde{\mathbf{F}}}^{\perp} \qquad , \qquad \hat{\mathbf{k}}^{\perp} = (\hat{\mathbf{k}}^{\perp}\cdot\hat{\tilde{\mathbf{F}}})\hat{\tilde{\mathbf{F}}} + (\hat{\mathbf{k}}^{\perp}\cdot\hat{\tilde{\mathbf{F}}}^{\perp})\hat{\tilde{\mathbf{F}}}^{\perp} \tag{92}$$

where $\hat{\tilde{\mathbf{F}}} = \tilde{\mathbf{F}}/|\tilde{\mathbf{F}}|$ is a unit vector in the direction of $\tilde{\mathbf{F}}$. Substituting these expressions into Eq. (91), we have:

$$\boldsymbol{\tau}\tilde{\mathbf{F}} = (\tilde{\mathbf{F}}\cdot\hat{\mathbf{k}})(a+b)\underbrace{\Big((\hat{\mathbf{k}}\cdot\hat{\tilde{\mathbf{F}}})\hat{\tilde{\mathbf{F}}} + (\hat{\mathbf{k}}\cdot\hat{\tilde{\mathbf{F}}}^{\perp})\hat{\tilde{\mathbf{F}}}^{\perp}\Big)}_{\hat{\mathbf{k}}} + (\tilde{\mathbf{F}}\cdot\hat{\mathbf{k}}^{\perp})(a)\underbrace{\Big((\hat{\mathbf{k}}^{\perp}\cdot\hat{\tilde{\mathbf{F}}})\hat{\tilde{\mathbf{F}}} + (\hat{\mathbf{k}}^{\perp}\cdot\hat{\tilde{\mathbf{F}}}^{\perp})\hat{\tilde{\mathbf{F}}}^{\perp}\Big)}_{\hat{\mathbf{k}}^{\perp}} \tag{93}$$

Collecting $\hat{\tilde{\mathbf{F}}}$ terms together and $\hat{\tilde{\mathbf{F}}}^{\perp}$ terms together, we have:

$$\boldsymbol{\tau}\tilde{\mathbf{F}} = \Big[\Big((\hat{\tilde{\mathbf{F}}}\cdot\hat{\mathbf{k}})^2 + (\hat{\tilde{\mathbf{F}}}\cdot\hat{\mathbf{k}}^{\perp})^2\Big)a + (\hat{\tilde{\mathbf{F}}}\cdot\hat{\mathbf{k}})^2 b\Big]\,|\tilde{\mathbf{F}}|\hat{\tilde{\mathbf{F}}} \tag{94}$$

$$+\Big[\Big((\hat{\tilde{\mathbf{F}}}\cdot\hat{\mathbf{k}})(\hat{\tilde{\mathbf{F}}}^{\perp}\cdot\hat{\mathbf{k}}) + (\hat{\tilde{\mathbf{F}}}\cdot\hat{\mathbf{k}}^{\perp})(\hat{\tilde{\mathbf{F}}}^{\perp}\cdot\hat{\mathbf{k}}^{\perp})\Big)a + (\hat{\tilde{\mathbf{F}}}\cdot\hat{\mathbf{k}})(\hat{\tilde{\mathbf{F}}}^{\perp}\cdot\hat{\mathbf{k}})b\Big]\,|\tilde{\mathbf{F}}|\hat{\tilde{\mathbf{F}}}^{\perp} \tag{95}$$

where in the first line of the above (expression (94)), we recognize that the coefficient of $a$ is equal to 1, i.e. $(\hat{\tilde{\mathbf{F}}}\cdot\hat{\mathbf{k}})^2+(\hat{\tilde{\mathbf{F}}}\cdot\hat{\mathbf{k}}^{\perp})^2=1$ since both $\hat{\tilde{\mathbf{F}}}$ and $\hat{\mathbf{k}}$ are unit vectors. In the second line of the above (expression (95)), we can show that the coefficient of $a$ is equal to 0, i.e. $(\hat{\tilde{\mathbf{F}}}\cdot\hat{\mathbf{k}})(\hat{\tilde{\mathbf{F}}}^{\perp}\cdot\hat{\mathbf{k}})+(\hat{\tilde{\mathbf{F}}}\cdot\hat{\mathbf{k}}^{\perp})(\hat{\tilde{\mathbf{F}}}^{\perp}\cdot\hat{\mathbf{k}}^{\perp})=0$, while the coefficient of $b$ is equal to $-(\hat{\tilde{\mathbf{F}}}\cdot\hat{\mathbf{k}})(\hat{\tilde{\mathbf{F}}}\cdot\hat{\mathbf{k}}^{\perp})$ since $\hat{\tilde{\mathbf{F}}}^{\perp}\cdot\hat{\mathbf{k}}=-\hat{\tilde{\mathbf{F}}}\cdot\hat{\mathbf{k}}^{\perp}$. These identities allow for simplifications, and we finally obtain Eq. (M38):

$$\boldsymbol{\tau}\tilde{\mathbf{F}}=\Big(a(k)+(\hat{\tilde{\mathbf{F}}}\cdot\hat{\mathbf{k}})^2 b(k)\Big)\tilde{\mathbf{F}}+\Big(-(\hat{\tilde{\mathbf{F}}}\cdot\hat{\mathbf{k}})(\hat{\tilde{\mathbf{F}}}\cdot\hat{\mathbf{k}}^{\perp})b(k)\Big)\tilde{\mathbf{F}}^{\perp} \quad . \tag{96}$$

Note that $\tilde{\mathbf{F}}$ and $\tilde{\mathbf{F}}^{\perp}$ both have magnitude $|\tilde{\mathbf{F}}|$, so the action of $\boldsymbol{\tau}$ on $\tilde{\mathbf{F}}$ is proportional to $|\tilde{\mathbf{F}}|$, as it should be.

#### B. Transfer matrices' dependencies on wavevector and other parameters

**I. Perpendicular transfer coefficients $\tau_{\perp}$, $\tau_{\perp}^{t,b}$ are *always* negative**. This is evident since the quantities $b, b^{t,b} < 0$, as plotted in Fig. M3 as functions of $k$ and in Fig. S1 as functions of $h$. Analogous to Fig. M3, the plots in Fig. S1 show the functions $a(h), a^{t,b}(h)$ (shades of red) and $b(h), b^{t,b}(h)$ (shades of brown) as functions of $h$ using the base parameters $\eta=\eta_B=\eta_L=1, H=0.5, k=2\pi/L, L=20$. Since $b, b^{t,b}<0$ and $k_x$ and $k_y$ are positive, then it follows from Eqns. (M39) and (M40) that $\tau_{\perp}$, $\tau_{\perp}^{t,b}$ are negative. The inequalities $b, b^{t,b}<0$ are additionally shown analytically in section (IV C).

Figure S1 also shows that $a(h), a^{t,b}(h) > 0$ when considered as functions of $h$ for a large set of parameter values. The inequalities $a, a^{t,b} < 0$ are additionally shown analytically in section (IV C). Furthermore, while matrix elements of opposite-side transfers $a$ and $b$ always decrease with $h$, those of same-side transfers $a^{t,b}$ and $b^{t,b}$ can increase with $h$.

**II. The quantities $\tau_{\parallel}^{t,b}$ satisfy $\tau_{\parallel}^{t,b} > 0$.** This is evident since the sums $a^{t,b}+b^{t,b}$ satisfy $a^{t,b}+b^{t,b}>0$. The quantities $a^{t,b}+b^{t,b}$ are plotted in Fig. S2(e-l) as functions of $k$ and in Fig. S3(e-l) as functions of $h$ to show that they are negative for a large set of parameter values. From Eq. (M40), the expressions $a^{t,b}+b^{t,b}>0$ are lower bounds to $\tau_{\parallel}^{t,b}$ since $b^{t,b}<0$. A positive lower bound on $\tau_{\parallel}^{t,b}$ implies that $\tau_{\parallel}^{t,b}>0$. The inequalities $a^{t,b}+b^{t,b}>0$ are also shown analytically in section (IV C).

**III. The quantity $\tau_{\parallel}$ is positive or negative depending on k.** This is evident since

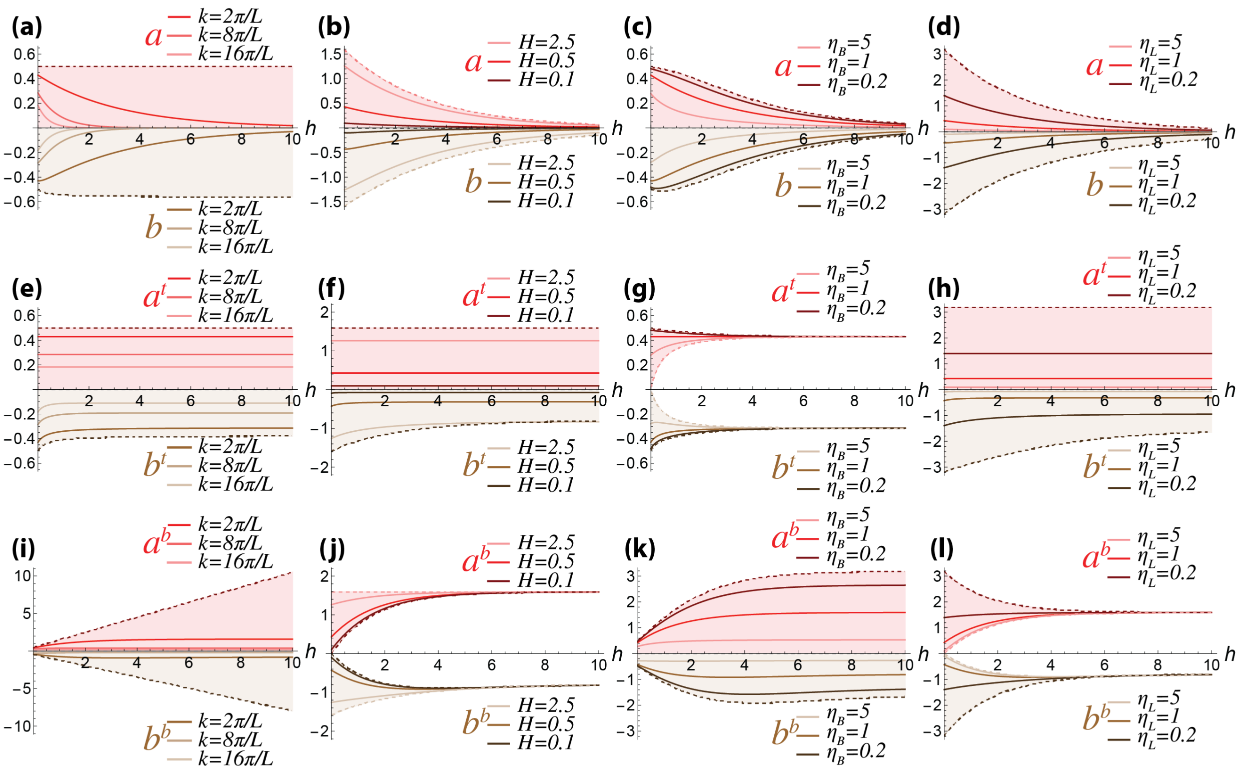


FIG. S1. Matrix elements of $\boldsymbol{\tau}, \boldsymbol{\tau}^{t,b}$. As functions of $h$, plots $a(h), a^{t,b}(h)$ and $b(h), b_k^{t,b}(h)$ considered as functions of $h$, the thickness of the slab, using the base parameters $\eta = \eta_B = \eta_L = 1, H = 0.5, L = 20$ with $k = 2\pi/L$, the magnitude of the fundamental mode. Here, each graph varies one of the parameters $k, H, \eta_B$, or $\eta_L$ by a numerical factor and plots the result in a different shade of red or brown. The range between which each of these parameters approaches 0 and $\infty$ are shaded lightly with the 0 limit indicated by dark dotted lines and the $\infty$ limit indicated by light dotted lines. (a-d) $a(h)$ and $b(h)$; (e-h) $a^t(h)$ and $b^t(h)$; (i-l) $a^b(h)$ and $b^b(h)$.

the sum $a + b$ satisfies $a + b < 0$. The quantity $a + b$ is plotted in Fig. S2(a-d) as a function of $k$ and in Fig. S3(a-d) as a function of $h$ to show that it is negative for a large set of parameter values. From Eq. (M39), the expression $a + b < 0$ is a lower bound to $\tau_{\|}$ since $a > 0$ and $b < 0$. A negative lower bound on $\tau_{\|}$ implies that $\tau_{\|}$ may be positive or negative depending on the wavevector $\mathbf{k}$. The inequality $a+b < 0$ is also shown analytically in section (IV C).

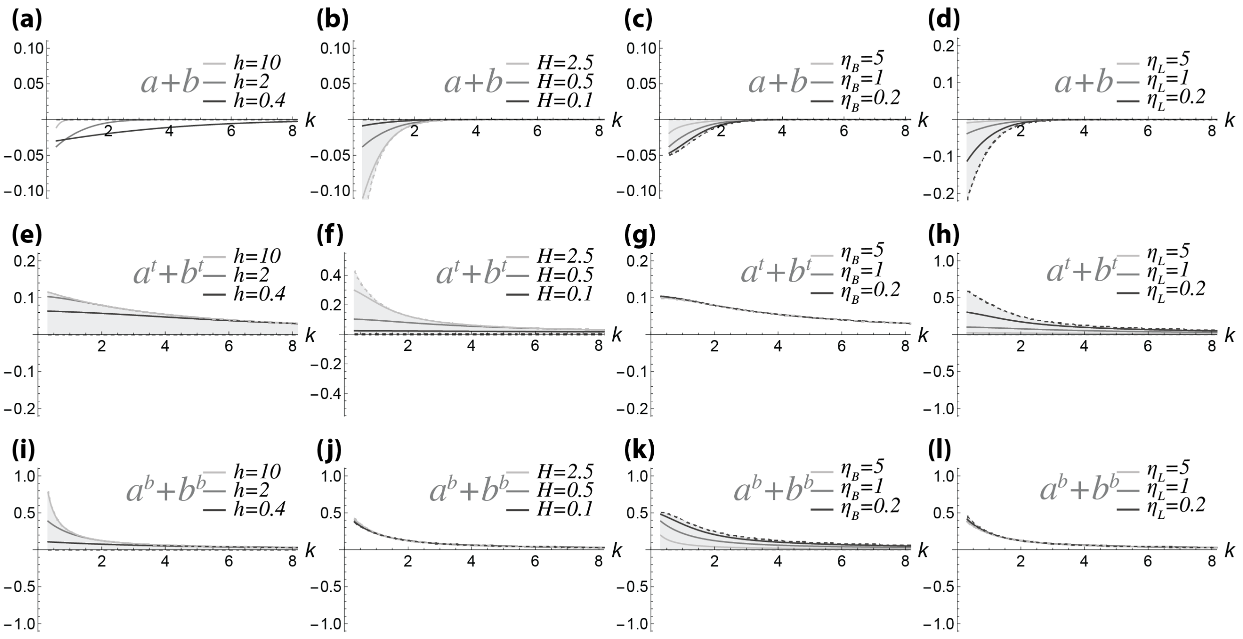


FIG. S2. As functions of $k$, plots the sums $a(k)+b(k)$ and $a^{t,b}(k)+b^{t,b}(k)$ using the base parameters $\eta = \eta_B = \eta_L = 1, H = 0.5, L = 20$ and $h = 2$. Each graph varies one of the parameters $h, H, \eta_B, \eta_L$ by a numerical factor and plots the result in a different shade of gray. The range between which each of these parameters approaches 0 and $\infty$ are shaded lightly with the 0 limit indicated by dark dotted lines and the $\infty$ limit indicated by light dotted lines. From this, we see that $a + b < 0$ and $a^{t,b} + b^{t,b} > 0$. (a-d) $a(k) + b(k)$; (e-h) $a^t(k) + b^t(k)$; (i-l) $a^b(k) + b^b(k)$.

#### *1. Parallel and perpendicular transfer coefficients*

Supplementary to Fig. M4, Figs. S4 and S5 indicate how the parallel and perpendicular transfer coefficients $\tau_\parallel$, $\tau_\perp$, $\tau_\parallel^{a,b}$, and $\tau_\perp^{a,b}$ depend on the wavevector $\mathbf{k}$ of the driving force $\tilde{F}_x$, as indicated in Eqns. (M39) and (M40). Figure S4 shows results for $h = 0.5\ell_0$, a short and flat cell, while Fig. S5 shows results for $h = 8\ell_0$, a tall cell more closely resembling the dimensions of a *Drosophila* cell in the ventral furrow and convergent-extension phases.

### C. Proofs that $a, a^{t,b} > 0$, $b, b^{t,b} < 0$, $a + b < 0$ and $a^{t,b} + b^{t,b} > 0$

In Figs. M3 and S1, we have plotted $a, b, a^{t,b}$ and $b^{t,b}$ and showed that at least for a large subset of parameters, we have $a, a^{t,b} > 0$ and $b, b^{t,b} < 0$. Additionally, in Figs. S2 and S3, we have plotted $a + b$ and $a^{t,b} + b^{t,b}$ and showed that at least for a large subset of parameters,

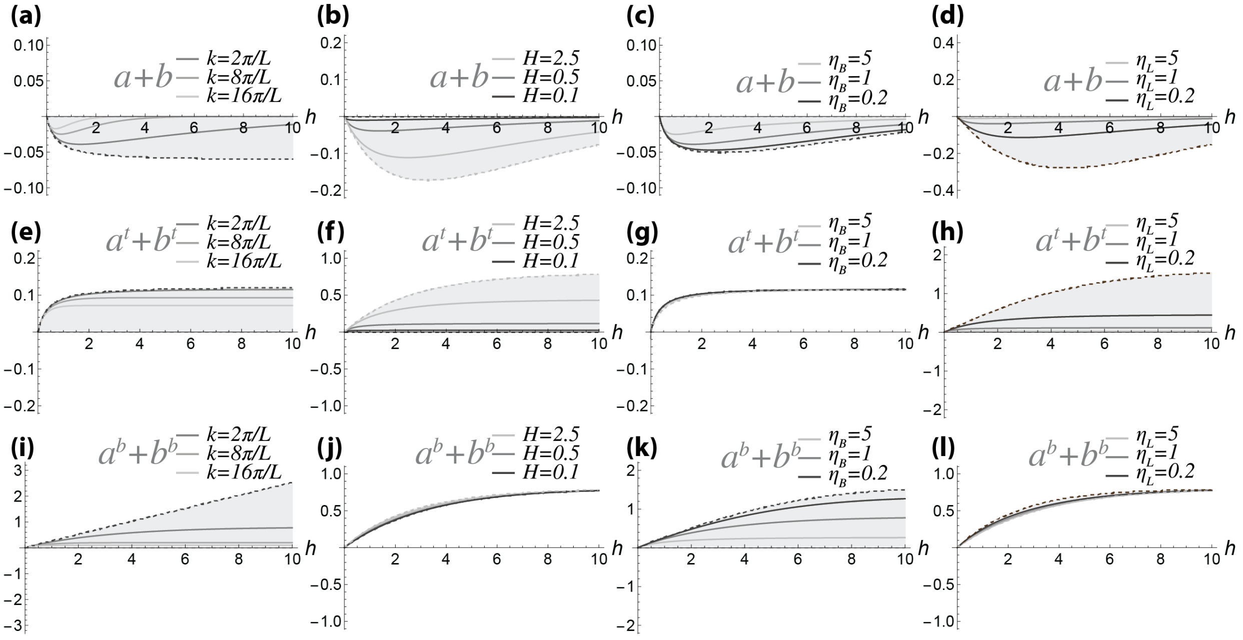

FIG. S3. As functions of $h$, plots the sums $a(h)+b(h)$ and $a^{t,b}(h)+b^{t,b}(h)$ using the base parameters $\eta = \eta_B = \eta_L = 1, H = 0.5, L = 20$ and $k = 2\pi/L$. Each graph varies one of the parameters $H, \eta_B, \eta_L$ by a numerical factor while $k$ is increased by numerical factors and plots the result in a different shade of gray. The range between which each of these parameters approaches 0 and $\infty$ are shaded lightly with the 0 limit indicated by dark dotted lines and the $\infty$ limit indicated by light dotted lines. From this, we see that $a + b < 0$ and $a^{t,b} + b^{t,b} > 0$. (a-d) $a(h) + b(h)$; (e-h) $a^t(h) + b^t(h)$; (i-l) $a^b(h) + b^b(h)$.

we have $a + b < 0$ and $a^{t,b} + b^{t,b} > 0$. Below, we prove that these statements are true for all physical values of the parameters.

Since the transfer matrices $\boldsymbol{\tau}$ and $\boldsymbol{\tau}^{t,b}$ are computed by multiplying, adding, and inverting matrices of the form $p(k)\mathbb{1} + q(k)\hat{\mathbf{k}}\hat{\mathbf{k}}$ (Eqs. (M31) and (M32)), then $a(k)$ and $b(k)$ are computed from various analytical functions of $k$ combined together by multiplication, division, and addition. Thus, all of $a, b, a^{t,b}$, and $b^{t,b}$ are analytical functions of $k$, and hence one should be able to tell by inspection whether any of them are positive or negative definite. However, the expressions $a, b, a^{t,b}, b^{t,b}$ are such complicated functions of $k$ that they are difficult to inspect. Our strategy to prove that any of these functions are positive or negative is to first show that the functions adding or multiplying to them are positive or negative, and then to deduce the signs of $a, a^{t,b}, b, b^{t,b}, a + b$, and $a^{t,b} + b^{t,b}$ from the the signs of these other

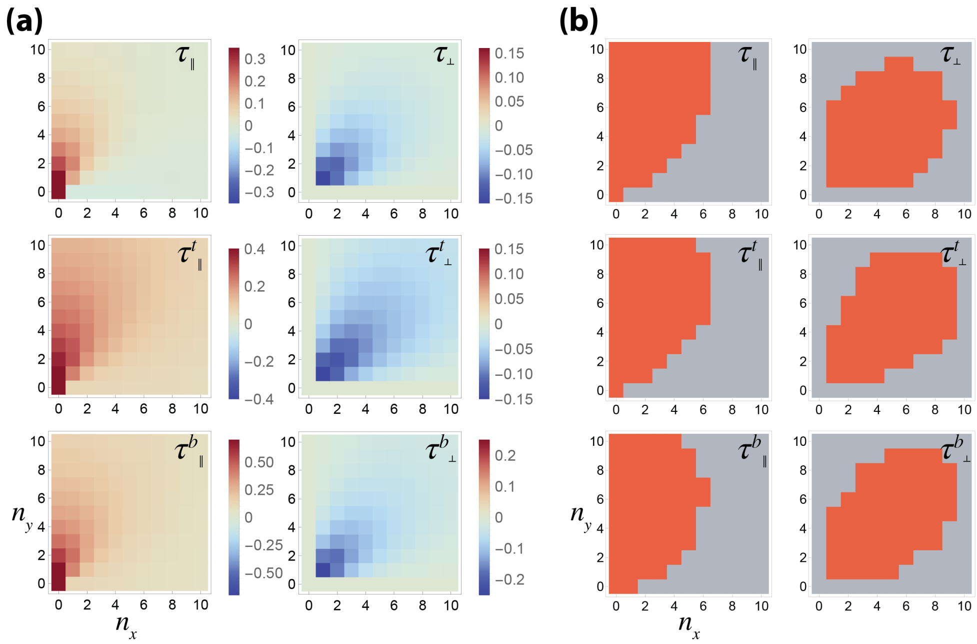


FIG. S4. Dependence of parallel and perpendicular transfer coefficients on wavevectors for $h = 0.5\ell_0$. (a) Coefficients of parallel and perpendicular transfer $\tau_\parallel$, $\tau_\perp$, $\tau_\parallel^{t,b}$, and $\tau_\perp^{t,b}$ plotted as functions of $\mathbf{k} = \frac{2\pi}{L}(n_x, n_y)$. (b) Wavevectors in which $|\tau_\parallel|$, $|\tau_\perp|$, $|\tau_\parallel^{t,b}|$, and $|\tau_\perp^{t,b}|$ are larger than their median values are highlighted in dark orange; for $\tau_\parallel$, only non-negative values greater than the median are highlighted.

functions. In other words, consider Eqns. (M31) and (M32) in which $\boldsymbol{\tau}, \boldsymbol{\tau}^{t,b}$ are given in terms of $\boldsymbol{\Gamma}$'s and $\boldsymbol{\Theta}$'s. We will show that matrix elements of $\boldsymbol{\tau}, \boldsymbol{\tau}^{t,b}$ are positive or negative based on the signs of the matrix elements of $\boldsymbol{\Gamma}$'s and $\boldsymbol{\Theta}$'s and their combinations.

First, let us simply notation. If we set $kh \equiv x$ and $kH \equiv xH/h$, then $x > 0$, and:

$$\boldsymbol{\Theta}_2(x, \eta) = p(x)\mathbb{1} + q(x)\hat{\mathbf{k}}\hat{\mathbf{k}} \tag{97}$$

$$\boldsymbol{\Theta}_1(x, \eta) = r(x)\mathbb{1} + s(x)\hat{\mathbf{k}}\hat{\mathbf{k}} \tag{98}$$

$$\boldsymbol{\Lambda}(x, \eta_L) = \boldsymbol{\Theta}_1\left(\frac{H}{h}x, \eta_L\right) = r^H(x)\mathbb{1} + s^H(x)\hat{\mathbf{k}}\hat{\mathbf{k}} \tag{99}$$

$$\boldsymbol{\Gamma}_L(x, \eta, \eta_L) = (r(x) + r^H(x))\mathbb{1} + (s(x) + s^H(x))\hat{\mathbf{k}}\hat{\mathbf{k}} \tag{100}$$

$$\boldsymbol{\Gamma}_B(x, \eta, \eta_B) = \left(r(x) + \eta_B\frac{x}{h}\right)\mathbb{1} + \left(s(x) + \eta_B\frac{x}{h}\right)\hat{\mathbf{k}}\hat{\mathbf{k}} \tag{101}$$

where we have absorbed the $\eta, \eta_L, \eta_B$ parameters into the definitions of the functions

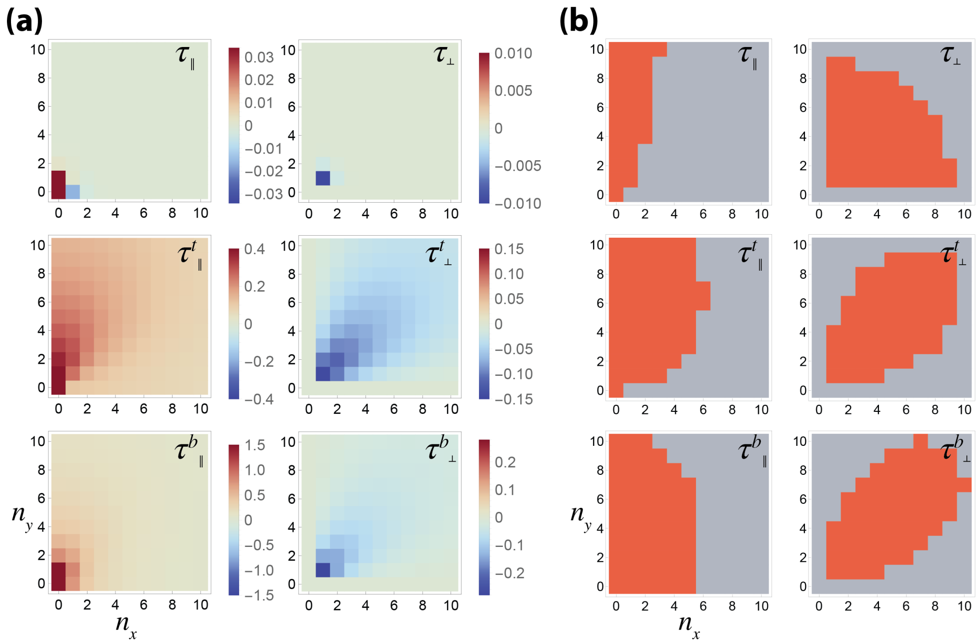


FIG. S5. Dependence of parallel and perpendicular transfer coefficients on wavevectors for $h = 8.0\ell_0$. (a) Coefficients of parallel and perpendicular transfer $\tau_\parallel$, $\tau_\perp$, $\tau_\parallel^{t,b}$, and $\tau_\perp^{t,b}$ plotted as functions of $\mathbf{k} = \frac{2\pi}{L}(n_x, n_y)$. (b) Wavevectors in which $|\tau_\parallel|$, $|\tau_\perp|$, $|\tau_\parallel^{t,b}|$, and $|\tau_\perp^{t,b}|$ are larger than their median values are highlighted in dark orange; for $\tau_\parallel$, only non-negative values greater than the median are highlighted.

$p, q, r, s, r^H$, and $s^H$, with

$$\begin{aligned} p(x) &= \frac{\eta x}{h\sinh(x)} \quad , \quad & q(x) =& \frac{\eta x}{2h}\left(\frac{-x\alpha_C^0(x)}{\sinh(x)} + \alpha^0(x)\right) \\ r(x) &= \frac{\eta x}{h}\coth(x) \quad , \quad & s(x) =& \frac{\eta x}{2h}\left(\frac{-x\alpha^0(x)}{\sinh(x)} + \alpha_C^0(x)\right) \\ r^H(x) &= \frac{\eta_L x}{h}\coth\left(\frac{H}{h}x\right) \quad , \quad & s^H(x) =& \frac{\eta_L x}{2h}\left(\frac{-xH\alpha^0\left(\frac{H}{h}x\right)}{h\sinh\left(\frac{H}{h}x\right)} + \alpha_C^0\left(\frac{H}{h}x\right)\right) \end{aligned} \tag{102}$$

where $\alpha^0(x)$ and $\alpha_C^0(x)$ are parameter-free functions of $x$, rewritten from Eq. (35) using the argument $x$:

$$\alpha_C^0(x) = 2\frac{\sinh(x)\cosh(x) - x}{\sinh(x)^2 - x^2} \quad , \quad \alpha^0(x) = 2\frac{\sinh(x) - x\cosh(x)}{\sinh(x)^2 - x^2} \tag{103}$$

The functions $\alpha^0\left(\frac{H}{h}x\right)$ and $\alpha_C^0\left(\frac{H}{h}x\right)$ contain the factor $H/h$ but this simply rescales the argument $x$ by a positive numerical factor and does not change the sign of $\alpha^0$ or $\alpha_C^0$.

To finally prove the positivity or negativity of $a, b, a^{t,b}, b^{t,b}$, etc, there are a few facts that we establish. Since $p, q, r, s$ are all equal to a positive factor $\eta/h$ multiplying a parameter-free function of $x \equiv kh > 0$, we have that:

$$\begin{aligned} p(x) >& 0 \quad , \quad q(x) < 0 \quad , \quad r(x) > 0 \quad , \quad s(x) > 0 \\ r^H(x) >& 0 \quad , \quad s^H(x) > 0 \quad . \end{aligned} \tag{104}$$

We establish all of these by inspection of the parameter-free functions of $x$ in $p, q, r, s$. Additionally, even though $r^H$ and $s^H$ contain factors of $H/h$ compared to $r$ and $s$, these simply scale the argument $x$, so they are shown to be positive by inspection as well. As an example of determining the sign of a function by inspection, we examine $q(x)$; after simplification, we find that:

$$q(x) = \frac{\eta x}{h \sinh(x)} \frac{x^2 - 2x\cosh(x)\sinh(x) + \sinh(x)^2}{\sinh(x)^2 - x^2} \tag{105}$$

The first factor of $q(x)$ is clearly positive since $x > 0$; one can also show that the numerator of the second factor is negative while the denominator of the second factor is positive; one can do this by series expansion. Hence the whole expression for $q(x)$ is negative. The same type of argument can be applied to $p, r, s, r^H$, and $s^H$ to determine their signs. Another useful fact is that

$$p(x) + q(x) = \frac{2\eta x}{h \sinh(x)} \frac{x\coth(x) - 1}{x^2 \operatorname{csch}(x)^2 - 1} < 0 \tag{106}$$

again derived by simplification of $p + q$ and then by inspection. Finally, a useful identity is that the inverse of $\mathbf{\Theta}_2$ is:

$$\mathbf{\Theta}_2^{-1} = \frac{1}{p}\left(\mathbb{1} + \frac{-q}{(p+q)}\hat{\mathbf{k}}\hat{\mathbf{k}}\right) \quad . \tag{107}$$

To eventually compute $\boldsymbol{\tau} = -\left(\mathbf{\Theta}_2 - \mathbf{\Gamma}_B \mathbf{\Theta}_2^{-1} \mathbf{\Gamma}_L\right)^{-1}$, one must first compute:

$$\mathbf{\Theta}_2^{-1}\mathbf{\Gamma}_L = \frac{1}{p}\left(\mathbb{1} + \frac{-q}{(p+q)}\hat{\mathbf{k}}\hat{\mathbf{k}}\right)\left((r + r^H)\mathbb{1} + (s + s^H)\hat{\mathbf{k}}\hat{\mathbf{k}}\right) \tag{108}$$

$$= \frac{r + r^H}{p}\mathbb{1} + \frac{1}{p}\left(\frac{-q}{p+q}(r + r^H) + \frac{p}{p+q}(s + s^H)\right)\hat{\mathbf{k}}\hat{\mathbf{k}} \tag{109}$$

$$\equiv \ell_a \mathbb{1} + \ell_b \hat{\mathbf{k}}\hat{\mathbf{k}} \tag{110}$$

where the coefficient of $\mathbb{1}$ is defined as $\ell_a$ and the coefficient of $\hat{\mathbf{k}}\hat{\mathbf{k}}$ is defined as $\ell_b$. Since $r, r^H, p > 0$, we deduce that

$$\ell_a = \frac{r + r^H}{p} > 0 \tag{111}$$

After simplification, we deduce that:

$$\ell_b = \frac{-q(r + r^H) + p(s + s^H)}{p(p + q)} < 0 \tag{112}$$

This is because $q < 0$, so all parts of the numerator are positive; also, since $p > 0$ and $p + q < 0$ from Eqns. (104) and (106), then the denominator is negative, making $\ell_b$ negative overall. We additionally deduce that the sum:

$$\ell_a + \ell_b = \frac{r + r^H + s + s^H}{p + q} < 0 \tag{113}$$

since $p + q < 0$ from Eq. (106) and the numerator is positive.

Now, as part of $\boldsymbol{\tau}$, we compute:

$$\begin{aligned}
\mathbf{\Gamma}_B \mathbf{\Theta}_2^{-1} \mathbf{\Gamma}_L =& \left( \left( r + \eta_B \frac{x}{h} \right) \mathbb{1} + \left( s + \eta_B \frac{x}{h} \right) \hat{\mathbf{k}}\hat{\mathbf{k}} \right) (\ell_a \mathbb{1} + \ell_b \hat{\mathbf{k}}\hat{\mathbf{k}}) & (114) \\
=& \left( r + \eta_B \frac{x}{h} \right) \ell_a \mathbb{1} + \left( \left( s + \eta_B \frac{x}{h} \right) (\ell_a + \ell_b) + \left( r + \eta_B \frac{x}{h} \right) \ell_b \right) \hat{\mathbf{k}}\hat{\mathbf{k}} & (115) \\
\equiv& g_a \mathbb{1} + g_b \hat{\mathbf{k}}\hat{\mathbf{k}} & (116)
\end{aligned}$$

where the coefficient of $\mathbb{1}$ is defined as $g_a$ and the coefficient of $\hat{\mathbf{k}}\hat{\mathbf{k}}$ is defined as $g_b$. Now,

$$g_a = \left( r + \eta_B \frac{x}{h} \right) \ell_a > 0 \tag{117}$$

because $\ell_a > 0$ from Eq. (111) and $r > 0$. Also we have:

$$g_b = \left( s + \eta_B \frac{x}{h} \right) (\ell_a + \ell_b) + \left( r + \eta_B \frac{x}{h} \right) \ell_b < 0 \tag{118}$$

because $\ell_a + \ell_b < 0$ from Eq. (113), $\ell_b < 0$ from Eq. (112), and the factors multiplying each of them are positive. Furthermore, after simplification, we have that the sum:

$$g_a + g_b = \frac{(r + r^H + s + s^H)(r + s + 2\frac{\eta_B}{h} x)}{p + q} < 0 \tag{119}$$

because $p + q < 0$ and the numerator is positive.

Here we compute:

$$\mathbf{\Theta}_2 - \mathbf{\Gamma}_B \mathbf{\Theta}_2^{-1} \mathbf{\Gamma}_L = (p - g_a) \mathbb{1} + (q - g_b) \hat{\mathbf{k}}\hat{\mathbf{k}} \equiv t_a \mathbb{1} + t_b \hat{\mathbf{k}}\hat{\mathbf{k}} \tag{120}$$

where the coefficient of $\mathbb{1}$ is defined as $t_a$ and the coefficient of $\hat{\mathbf{k}}\hat{\mathbf{k}}$ is defined as $t_b$. We will show that $t_a < 0$. Now since both $p > 0$ and $g_a > 0$, it is difficult to tell the sign of $p - g_a$ right away. However, we can expand $g_a$ using Eqns. (117) and (111):

$$t_a = p - g_a = p - \left( r + \eta_B \frac{x}{h} \right) \left( \frac{r + r^H}{p} \right) = p - \frac{r^2}{p} - \text{positive numbers} < p - \frac{r^2}{p} \quad . \tag{121}$$

In the above, we noted that $p - g_a$ is upper bounded by $p - r^2/p$ since if we expand the multiplication:

$$\left(r + \eta_B \frac{x}{h}\right)\left(\frac{r + r^H}{p}\right) = \frac{r^2 + \eta_B \frac{x}{h} r + r r^H + \eta_B \frac{x}{h} r^H}{p} \quad , \tag{122}$$

then all terms are positive; however, they are all subtracted from $p$, so that the result of keeping all terms must be less than that of truncating these terms. Substituting expressions for $p$ and $r$ from Eq. (102), we find that the upper bound of $t_a$ is negative and hence $t_a < 0$:

$$t_a < p - \frac{r^2}{p} = \frac{-\eta}{h} x \sinh(x) < 0 \quad . \tag{123}$$

We will show that $t_a + t_b$ is positive.

$$t_a + t_b = \frac{(p+q)^2 - (r + s + r^H + s^H)(r + s + 2\eta_B \frac{x}{h})}{p+q} \tag{124}$$

We already see that the denominator is negative since $p + q < 0$; if we determine the sign of the numerator, then we will know the sign of $t_a + t_b$. The numerator is upper bounded:

$$\text{numerator} = (p+q)^2 - (r + r^H + s + s^H)(r + s + 2\eta_B x/h) < (p+q)^2 - (r+s)^2 \tag{125}$$

Again, we use explicit expressions for $p, q, r$, and $s$ from Eq. (102) to obtain:

$$\text{numerator} < (p+q)^2 - (r+s)^2 = \frac{4\eta^2 x^2}{h^2(x^2 \operatorname{csch}(x)^2 - 1)} < 0 \tag{126}$$

Since the numerator of $t_a + t_b$ is upper bounded by a negative function, then the numerator is negative; since the denominator of $t_a + t_b$ is also negative, we deduce that:

$$t_a + t_b > 0 \tag{127}$$

Since we have already shown that $t_a < 0$ and $t_a + t_b > 0$, then we find that $t_b > 0$ because:

$$t_b = (t_a + t_b) - (t_a) = \text{positive} - \text{negative} > 0 \quad . \tag{128}$$

Showing that $a > 0, b < 0$ and $a + b < 0$:

Using the results so far in this section, we are now ready to calculate $\boldsymbol{\tau}$:

$$\boldsymbol{\tau} = -\left(\boldsymbol{\Theta}_2 - \boldsymbol{\Gamma}_B \boldsymbol{\Theta}_2^{-1} \boldsymbol{\Gamma}_L\right)^{-1} = -\left(t_a \mathbb{1} + t_b \hat{\mathbf{k}}\hat{\mathbf{k}}\right)^{-1} = \frac{-1}{t_a}\mathbb{1} + \frac{t_b}{t_a(t_a + t_b)}\hat{\mathbf{k}}\hat{\mathbf{k}} \tag{129}$$

$$\equiv a + b\hat{\mathbf{k}}\hat{\mathbf{k}} \tag{130}$$

where $a$ and $b$ are defined here as in the main text, as the coefficient of $\mathbb{1}$ and $\hat{\mathbf{k}}\hat{\mathbf{k}}$, respectively, in the matrix $\boldsymbol{\tau}$. Since we have already deduced the signs of $t_a$, $t_b$, and $t_a$+$t_b$ from Eqns. (123), (128), and (127), then it is easy to see that:

$$a = \frac{-1}{t_a} > 0 \qquad , \qquad b = \frac{t_b}{t_a(t_a + t_b)} < 0 \quad . \tag{131}$$

Meanwhile, the sum:

$$a + b = \frac{-1}{t_a + t_b} < 0 \quad . \tag{132}$$

<u>Showing that $a^t > 0, b^t < 0$ and $a^t + b^t > 0$:</u>

Since $\boldsymbol{\tau}^t = \boldsymbol{\tau}\boldsymbol{\Gamma}_B\boldsymbol{\Theta}_2^{-1}$, we first need to calculate $\boldsymbol{\Gamma}_B\boldsymbol{\Theta}_2^{-1}$:

$$\begin{aligned}
\boldsymbol{\Gamma}_B\boldsymbol{\Theta}_2^{-1} =& \boldsymbol{\Theta}_2^{-1}\boldsymbol{\Gamma}_B = \frac{1}{p}\left(\mathbb{1} + \frac{-q}{(p+q)}\hat{\mathbf{k}}\hat{\mathbf{k}}\right)\left(\left(r + \eta_B\frac{x}{h}\right)\mathbb{1} + \left(s + \eta_B\frac{x}{h}\right)\hat{\mathbf{k}}\hat{\mathbf{k}}\right) && (133)\\
=& \frac{\left(r + \eta_B\frac{x}{h}\right)}{p}\mathbb{1} + \left(\frac{-q}{p(p+q)}\left(r + s + 2\eta_B\frac{x}{h}\right) + \frac{\left(s + \eta_B\frac{x}{h}\right)}{p}\right)\hat{\mathbf{k}}\hat{\mathbf{k}} && (134)\\
\equiv& f_a\mathbb{1} + f_b\hat{\mathbf{k}}\hat{\mathbf{k}} && (135)
\end{aligned}$$

where $f_a$ and $f_b$ are defined as the coefficient of $\mathbb{1}$ and $\hat{\mathbf{k}}\hat{\mathbf{k}}$, respectively. We conclude that:

$$f_a = \frac{\left(r + \eta_B\frac{x}{h}\right)}{p} > 0 \tag{136}$$

because $p, r > 0$. Additionally, after simplifying,

$$f_b = \frac{-q(r + \eta_B\frac{x}{h}) + p(s + \eta_B\frac{x}{h})}{p(p+q)} < 0 \tag{137}$$

This is because $q < 0$, so the numerator is positive; meanwhile, since $p > 0$ and $p + q < 0$, the denominator is negative. So $f_b < 0$ overall. For the sum $f_a + f_b$ we have, after simplification:

$$f_a + f_b = \frac{r + s + 2\eta_B\frac{x}{h}}{p+q} < 0 \tag{138}$$

because the numerator is positive while the denominator $p + q < 0$.

Knowing the signs of $f_a, f_b$ and $f_a + f_b$ from Eqns. (136), (137), and (138), one can deduce the signs of the matrix elements of $\boldsymbol{\tau}^t$ as follows:

$$\boldsymbol{\tau}^t = \boldsymbol{\tau}\left(\boldsymbol{\Gamma}_B\boldsymbol{\Theta}_2^{-1}\right) = \left(a\mathbb{1} + b\hat{\mathbf{k}}\hat{\mathbf{k}}\right)\left(f_a\mathbb{1} + f_b\hat{\mathbf{k}}\hat{\mathbf{k}}\right) \tag{139}$$

$$= af_a\mathbb{1} + (b(f_a + f_b) + af_b)\hat{\mathbf{k}}\hat{\mathbf{k}} \tag{140}$$

$$\equiv a^t\mathbb{1} + b^t\hat{\mathbf{k}}\hat{\mathbf{k}} \tag{141}$$

where $a^t$ and $b^t$ are defined here as in the main text, as the coefficient of $\mathbb{1}$ and $\hat{\mathbf{k}}\hat{\mathbf{k}}$, respectively, in the matrix $\boldsymbol{\tau}^t$. We have:

$$a^t = af_a > 0 \tag{142}$$

since $a, f_a > 0$. Substituting Eqns. (138) and (137) for $f_a + f_b$ and $f_b$ in the expression for $b^t$ and simplifying, we have :

$$b^t = \frac{-q(r + \eta_B\frac{x}{h}) + p(s + \eta_B\frac{x}{h})}{p(p+q)} < 0 \quad . \tag{143}$$

This is because $-q > 0$ so the numerator of $b^t$ is positive, meanwhile the denominator is negative since $p + q < 0$. For the sum of $a^t + b^t$, we have:

$$a^t + b^t = (a + b)(f_a + f_b) > 0 \tag{144}$$

since $a+b < 0$ and $f_a+f_b < 0$. Hence, we have shown that $a^t > 0, b^t < 0$ and $a^t+b^t > 0$.

Showing that $a^b > 0, b^b < 0$ and $a^b + b^b > 0$:

Since $\boldsymbol{\tau}^t = \boldsymbol{\tau}\boldsymbol{\Gamma}_L\boldsymbol{\Theta}_2^{-1}$, we will use the expression for $\boldsymbol{\Gamma}_L\boldsymbol{\Theta}_2^{-1} = \boldsymbol{\Theta}_2^{-1}\boldsymbol{\Gamma}_L$ in Eq. (110):

$$\boldsymbol{\tau}^t = \boldsymbol{\tau}\left(\boldsymbol{\Gamma}_L\boldsymbol{\Theta}_2^{-1}\right) = \left(a\mathbb{1} + b\hat{\mathbf{k}}\hat{\mathbf{k}}\right)\left(\ell_a\mathbb{1} + \ell_b\hat{\mathbf{k}}\hat{\mathbf{k}}\right) \tag{145}$$

$$= \ell_a a\mathbb{1} + (b(\ell_a + \ell_b) + a\ell_b)\hat{\mathbf{k}}\hat{\mathbf{k}} \tag{146}$$

$$\equiv a^b\mathbb{1} + b^b\hat{\mathbf{k}}\hat{\mathbf{k}} \tag{147}$$

where $a^b$ and $b^b$ are defined here as in the main text, as the coefficient of $\mathbb{1}$ and $\hat{\mathbf{k}}\hat{\mathbf{k}}$, respectively, in the matrix $\boldsymbol{\tau}^b$. From above we see that:

$$a^b = \ell_a a > 0 \tag{148}$$

since both $\ell_a, a > 0$ from Eqns. (131) and (111). Additionally, we have:

$$a^b + b^b = (a + b)(\ell_a + \ell_b) > 0 \tag{149}$$

since $a+b<0$ and $\ell_a+\ell_b<0$ from Eqns. (132) and (113). Finally, to show that $b^b<0$, we write:

$$b^b = b(\ell_a+\ell_b)+a\ell_b = (a+b)\ell_b + b\ell_a \tag{150}$$

Expressing $a+b$ and $b$ in terms of $t_a$ and $t_b$ from Eqns. (132) and (131), we have:

$$b^b = \frac{-1}{t_a+t_b}\ell_b + \frac{t_b}{t_a(t_a+t_b)}\ell_a = \frac{-t_a\ell_b+t_b\ell_a}{t_a(t_a+t_b)} \tag{151}$$

Since $t_a<0$ and $t_a+t_b>0$ from Eqns. (123) and (127), then the denominator of $b^b$ is negative. If we can show that the numerator of $b^b$ is positive, i.e. $-t_a\ell_b+t_b\ell_a>0$, then we have shown that $b^b<0$. The signs of $t_a, \ell_a, t_b, \ell_b$ alone are not enough for us to deduce the sign of $-t_a\ell_b+t_b\ell_a$. So we proceed with substituting $-t_a\ell_b+t_b\ell_a$ in terms of $p, q, r, s, r^H, s^H$. After simplification, we have:

$$-t_a\ell_b+t_b\ell_a = -\frac{-2pq(r+r^H)+p^2(s+s^H)+(r+r^H)\big(-q^2+(r+r^H+s+s^H)(s+\eta_B\frac{x}{h})\big)}{p(p+q)} \tag{152}$$

Since the denominator of the above expression $p(p+q)<0$, then, considering the overall negative sign, to show that $-t_a\ell_b+t_b\ell_a>0$, we need only to show that the numerator of the above Eq. (152), without the overall $-$ sign, is positive. Now certainly $-2pq(r+r^H)>0$ since $p, r, r^H>0$ and $q<0$; additionally, we have $p^2(s+s^H)>0$. So the first two terms of the numerator are positive. If we can show that the last term of the numerator $(r+r^H)\big(-q^2+(r+r^H+s+s^H)(s+\eta_B\frac{x}{h})\big)>0$, then we are done. Since $(r+r^H)>0$, we really only need to show that $-q^2+(r+r^H+s+s^H)(s+\eta_B\frac{x}{h})>0$; we do this with a bounding argument. Certainly:

$$-q^2+(r+r^H+s+s^H)\Big(s+\eta_B\frac{x}{h}\Big) > -q^2+s^2 \tag{153}$$

because the terms in the expansion $(r+r^H+s+s^H)(s+\eta_B\frac{x}{h})$ are all positive. Evaluating $-q^2+s^2$ explicitly from Eq. (102), we find that

$$-q^2+s^2 = \frac{\eta^2x^2}{h^2} > 0 \tag{154}$$

Since the lower bound of $-q^2+(r+r^H+s+s^H)(s+\eta_B\frac{x}{h})$ is positive, then $-q^2+(r+r^H+s+s^H)(s+\eta_B\frac{x}{h})>0$. Hence

$$b^b > 0 \tag{155}$$

as promised. We have shown that $a^b>0$, $b^b<0$, and $a^b+b^b>0$.

### V. VELOCITY RESPONSE TO PERIODIC DRIVING WITH FULL OLDROYD-B MODEL: ANGULAR RESPONSE

Supplementary to Fig. M6, Fig. S6 plots the angular components of the velocity response (Eq. (M43)) under a surface driving force that is oscillatory in time (Eq. (M42)).

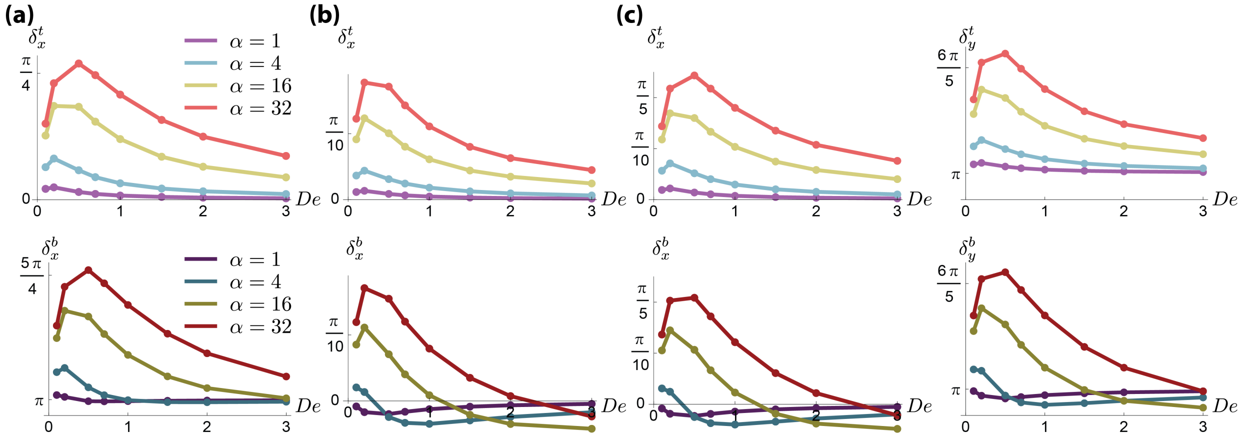


FIG. S6. Velocity response as a function of Deborah number for the driving force in Eq. (M42) with (a) wavevector $\mathbf{k} = \frac{2\pi}{L}(1,0)$, (b) wavevector $\mathbf{k} = \frac{2\pi}{L}(0,1)$, and (c) wavevector $\mathbf{k} = \frac{2\pi}{L}(1,1)$. Color key for (b) and (c) are same as in (a).

### VI. INSIGHT INTO VENTRAL FURROW FORMATION

#### A. Parameter plot for apical constriction with $\eta_L = 0.1\eta$

Supplementary to Fig. M8(b), we show a parameter space diagram in Fig. S7 for the case $\eta_L = 0.1$ with other parameters held same as in Fig. M8(b). Figure S7 shows that for the case $\eta_L = 0.1$, the magnitude of stress $S_0$ required to match experiments is around half of that for the case $\eta_L = 1.0$, assuming the same $\alpha$.

#### B. Decreasing viscosity in the fluid layer reduces tissue invagination

Supplementary to Fig. M8(d), Fig. S8 plots the $z$-directional force density on the tissue's top and bottom surface, $F_z(\text{top})$ and $F_z(\text{bottom})$, respectively, as $\eta_L$ is decreased (left to right panels) with other parameters held same as Fig. M8 ($\eta = \eta_B = 1, h = 8, H = 0.5$). As $\eta_L$ decreases, the resultant $z$-directional force on the tissue's top surface becomes less negative

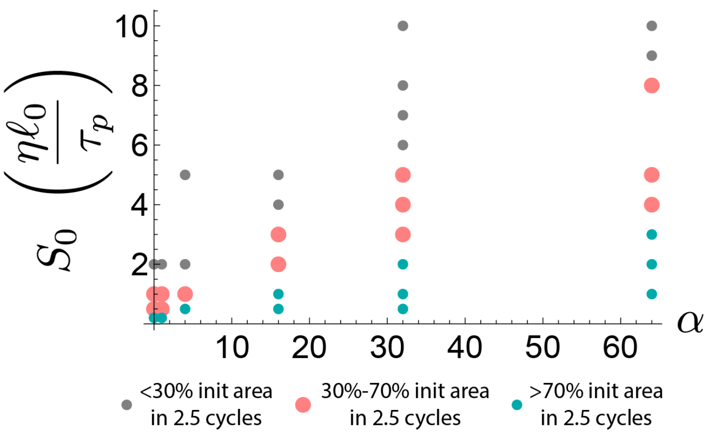


FIG. S7. Simulation of apical constriction: diagram of parameter space $\alpha$ and $S_0$; pink dots indicate simulations that agree with experiments. Parameters: $\eta = \eta_B = 1, \eta_L = 0.1, h = 8, H = 0.5$.

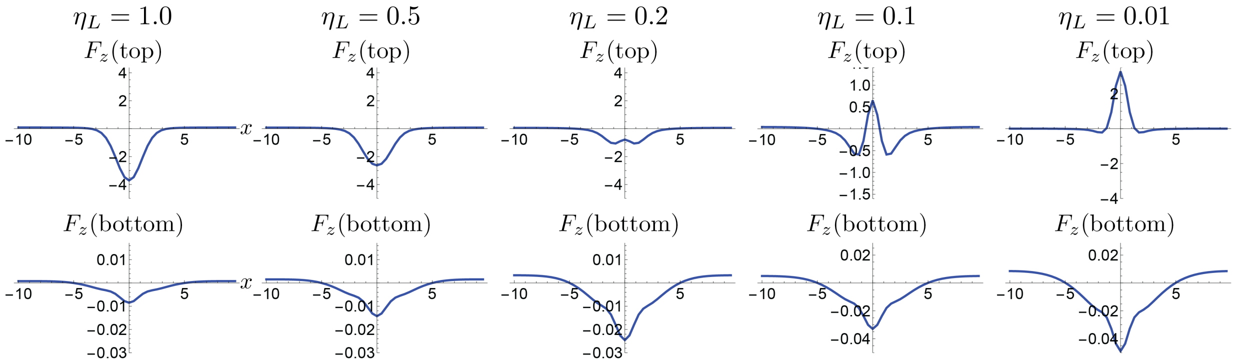


FIG. S8. As $\eta_L$ is decreased (left to right panels), the resultant $z$-directional force on the tissue's top surface becomes less negative and eventually positive, while the force on the tissue's bottom surface remains negative; that is, for sufficiently small values of $\eta_L$, the $z$-directional force would no longer cause invagination.

and eventually positive, meaning that the fluid layer pushes less strongly into the tissue as $\eta_L$ is decreased. Hence, with decreasing viscosity in the fluid layer, convergent active stresses at the tissue's top surface would act to make it locally thicker instead of causing it to invaginate. This situation is similar to increasing $H$, as shown in Fig. M8(d).

[1] M.B. Amar, A. Goriely, M.M. Müller, and L. Cugliandolo. *New Trends in the Physics and Mechanics of Biological Systems: Lecture Notes of the Les Houches Summer School: Volume 92, July 2009.* Lecture Notes of the Les Houch. OUP Oxford, 2011.
